# Neuro-musculoskeletal modeling reveals muscle-level neural dynamics of adaptive learning in sensorimotor cortex

**DOI:** 10.1101/2024.09.11.612513

**Authors:** Travis DeWolf, Steffen Schneider, Paul Soubiran, Adrian Roggenbach, Paolo Muratore, Mackenzie Weygandt Mathis

## Abstract

The neural activity of the brain is intimately coupled to the dynamics of the body. In order to predict and adapt to the sensorimotor consequences of our actions, compelling behavioral studies in humans, non-human primates, and in rodents have shown the existence of internal models – predictive models of our body in the environment. These internal models are theoretically used to compute an updated state estimate with prediction errors. Here, we directly test whether such errors are encoded in primary somatosensory (S1) or motor (M1) cortex during a motor adaptation task in mice. Using control theory-derived features that include prediction errors, we find that functionally distinct neurons are mapped onto specific computational motifs. We find that layer 2/3 population dynamics encode command-like signals and sensorimotor prediction errors (SPEs). S1 neurons encode SPEs more prominently than M1, and the neural latent dynamics change in S1 more than in M1 during this within-session learning. Then we asked, in which coordinate frameworks are such errors computed? To do this, we developed a novel 50-muscle model of the adult mouse forelimb that is capable of studying motor control and learning in a physics simulator. We identify both high-level 3D position and muscle spaces as coordinate frameworks for SPEs. Together, our results provide a new model of how neural dynamics in S1 enables adaptive learning.

## Introduction

Seamless execution of skilled actions requires animals to adapt in dynamic environments to maintain performance. Although many circuits from the cortex to the spinal cord are active and required to generate effective movements, it remains unclear how neural dynamics contribute to motor adaptation, the ability to learn from perturbations to restore performance. Several studies have shown that the primary somatosensory cortex (S1) (1–3) is critical for learning to adapt to motor perturbations, while others have argued that the motor cortex (M1) is critical for learning a new skill (4–6). Both insights may be mapped to mechanisms that support rapid plasticity (4, 7, 8) and updating of internal models (1, 9, 10). Internal models are updated when expected sensory (or motor) feedback does not match the forward prediction from the model. This “mismatch” in expected sensory feedback vs. actual feedback is used to compute a state estimate that then updates both motor commands and can be used to recalibrate the internal model if the error is persistent across movement attempts, hinting that the model was incorrect.

In human motor learning studies, there is ample evidence supporting the existence of internal models that guide adaptation across trials. This has been demonstrated in various tasks, such as a physical perturbation of the limb, or tasks that decouple visual feedback from movements of the limb or the eye (4, 11–17). In 2D planar reaching forelimb tasks in humans, upon introduction of a force field perturbation, hand movements are initially deviated from the baseline trajectory. Yet with repeated exposure to the same perturbation, one can compensate for the force field and restore performance similar to the baseline level by, theoretically, computing so called sensory prediction errors (SPEs) that are used to update an internal model (12, 18–23). Of particular interest are paradigms that use a delayed-onset force field such that there is a period of movement before the onset of a force-based perturbation to the hand (1, 20). Although this does not allow for restoration to baseline trajectories (sudden onset perturbations are difficult to predict (1, 20)), this has the advantage of observing the expected behavioral response to a perturbation of the limb – if you expect your hand to be deviated to the right, already counter to the left.

Ultimately, the large open question is where these internal models are stored and what they look like in the neural code. One path towards answering this question is to measure the SPEs that theoretically update the models. Many studies have focused on the visual system with tasks that decouple visual feedback from self-generated motion, causing a large mismatch signal (24, 25). Critically, SPEs have been shown to be in layer 2/3 excitatory neurons and not in other layers (such as the output layers 5 or 6). The detailed circuit defined by molecular type is an active area of study (25–28), but an emerging picture is that layer 2/3 neurons encode SPEs and these are used to update representations in layer 5/6 (29), or are shared across cortical columns (30) for rapid recalibration of internal models (29, 31). Moreover, since SPEs have recently been reported in the auditory cortex (32, 33), this indicates a potential canonical computation across the sensory areas in the cortex.

Notably though, it remains unclear in what coordinate frameworks such SPEs would be in during motor learning, despite this having implications for their mechanism. For example, in visual cortex errors could be computed when textures or edges are not correctly predicted, or if a higher-order feature like an object or face does not match what was predicted. In sensorimotor areas such as S1, this could be when the proprioceptive feedback from muscle spindles does not match the prediction or a more derived feature, such as when the 3D position of the hand does not match the predicted location. Thus, in *what* coordinate frameworks – kinematics, joint torques, or musculoskeletal features (34) –, and *how* the neural dynamics change in S1 during motor adaptation remains an open question. It could be that many internal models exist for separate coordinate frameworks. Thus, SPEs in different frameworks could update separate internal models and be used for direct motor command (policy) optimization (35–38). Moreover, it is unclear whether the motor cortex itself computes SPEs or changes its dynamics in a coordinate-specific manner, or if SPEs are only in sensory cortical areas, i.e., those receiving direct sensory thalamic input.

Thus, in order to ultimately answer in which coordinate frameworks such errors would be computed, the basic encoding properties of the circuit in study must be delineated. In motor adaptation tasks, most of the single-cell resolution data has come from studies in M1. A myriad of studies in primates have shown that single neurons in M1 can encode high-order kinematic features such as the hand direction, position and/or velocity (39, 40). However, several studies also demonstrated that much of M1 activity is related to lower-level features (41, 42), which cautions against assuming that M1 activity is best interpreted solely in terms of high-level end-point variables. In M1 of mice, a majority of deep layer (5) neurons increase their firing around forearm EMG movement events (43, 44). In L2/3 excitatory neurons in mice M1 show that approximately 40-50% of the neurons are tuned to events surrounding movements, such as licking or lever pressing (45–47), but their specific kinematic tuning properties were not described. In S1, in both primates and rodents, recent evidence points to encoding of the whole arm during movement (48–50) and S1 has been shown to increase firing immediately after peripheral perturbations (51, 52).

Only a few works in non-human primates have begun to explore the single-cell resolution neural basis of motor adaptation, focusing on premotor areas and motor cortex (M1) (4, 53). However, they have focused on higher-level encoding features, such as changes in the population preferred direction tuning and subspace neural dynamics during and after forelimb perturbations. While this is highly informative, it remains unknown if there are SPEs, particularly in S1, and if the neural dynamics encode specific higher- and lower-level (muscle features, proprioception) coordinate frameworks.

Given the evidence in different visual, auditory, and touch tasks for mismatch or SPEs coding in layer 2/3 (24, 28, 32, 33), we hypothesize that S1 is computing SPEs (1) in a task- and coordinate-relevant manner for learning to adapt. To test this hypothesis, we recorded from M1 and S1 in layer 2/3 excitatory neurons during a motor adaptation task for mice. Using unsupervised clustering, we discover that nearly 20% of the population encode prediction errors. Then, to test which coordinate frameworks are reflected in SPE-encoding neurons, we developed a novel *in silico* musculoskeletal-biomechanical model of adult mouse forelimb that can be controlled with an analytical solver to match the real kinematic behavior of mice performing a skilled reach, grasp, and pull task. In brief, we can control this system by solving the inverse kinematics and analytically calculating the muscle activations to follow a trajectory in joint space using muscle-based control: this specifically allows us to measure high- and low-level coordinate frameworks that would otherwise not be accessible.

## Results

### Movement encoding across the forelimb sensorimotor cortex during a joystick task

Mice have become an excellent model system for neural circuit studies during reaching and handling (1, 44, 45, 54–62). Here, we used a joystick-based reach, grasp, and pull task. In brief, head-fixed mice were trained to reach with their dominant hand to a vertical joystick that has 2 degrees of freedom (i.e., 360 degree movement) that was placed approximately 8 mm from the platform they stood on (see Methods, Figure 1a). They learned to move the joystick handle into a target zone that was programmatically delineated in order to receive a liquid reward. As with planar manipulandum tasks in humans and non-human primates (12, 20, 48), the advantage of head fixation is a more limited range of whole-body movements, yet unlike in most primate planar-movement tasks, here the mice must reach out, grasp, and learn to move the joystick to a reward zone, making the behavior 3D (not planar 2D). Over the course of 5-10 days they learned to control the robotic joystick to receive liquid rewards (65% of trials were rewarded during the two-photon imaging sessions), which also resulted in controlled bell-shaped velocity profiles during pulls (Figure 1b Videos 1-3).

**Figure 1.**
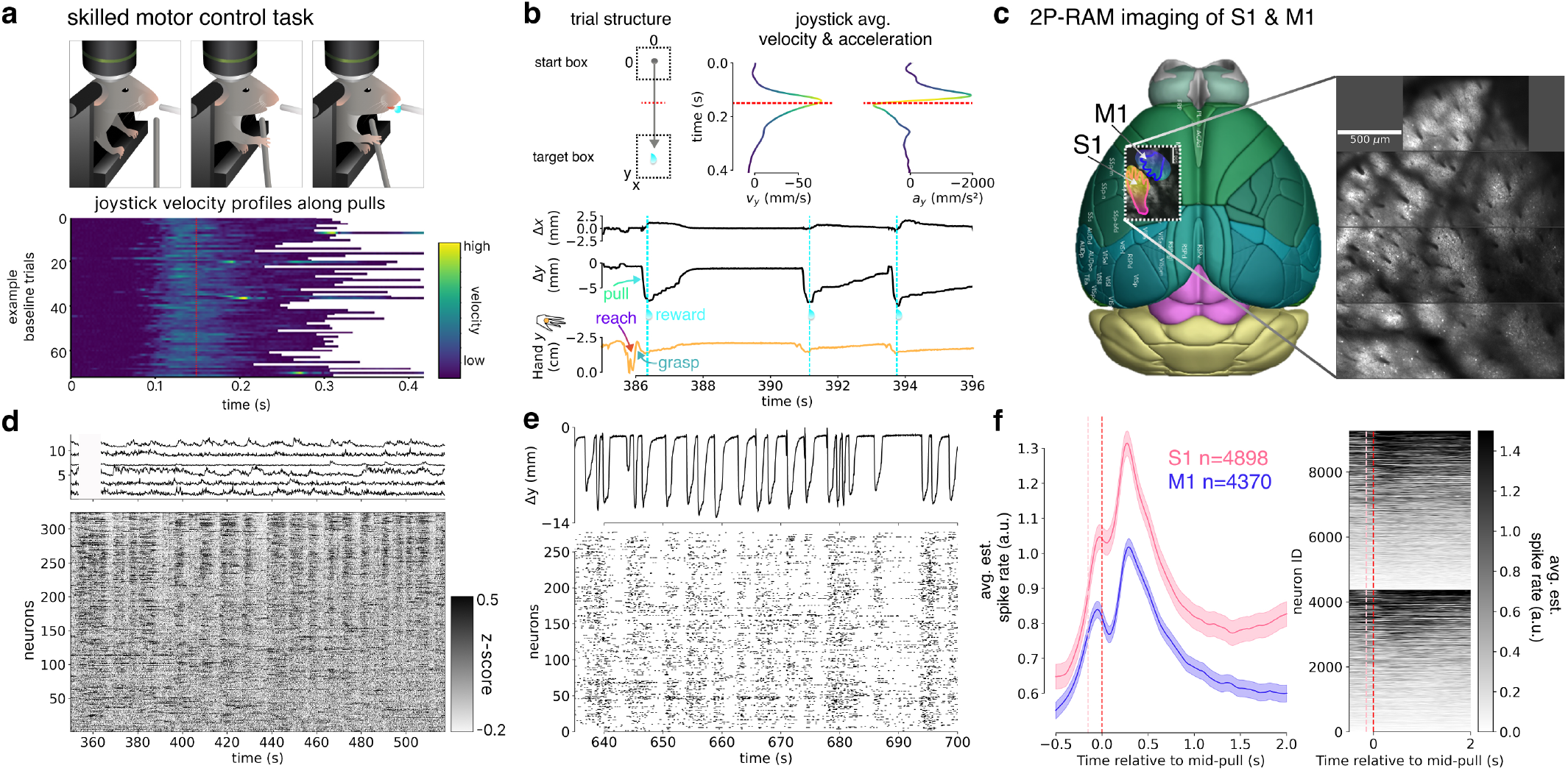
Activity of neurons in M1 and S1 during forelimb reach, grasp, and pull task. **a**, Schematic of the experimental setting showing a head-fixed mouse engaged in a reach, grab, and pull task, the session structure, with example velocity plots from a single session, aligned to a positional bin (highlighted in red). Cartoon by Julia Kuhl. **b**, Trial structure, average velocity (mean *±* SEM), and acceleration (mean *±* SEM) profiles across trials (n = 6 mice, 22 baseline sessions, n = 4,667 trials), and example joystick and hand trajectories from a individual mouse below. **c**, Example two-photon imaging area with the 2P-RAM system, covering forelimb S1 and M1 with example scan fields in a behaving mouse. Average intensity projection of four imaging fields recorded simultaneously at 5.8 Hz. **d**, A z-scored recording of ~400 neurons within a single baseline session, with example Δ*F/F* traces for approx. 1 min. **e**, Example deconvolved and binarized traces for neurons shown as a rastermap, plus joystick trajectory, for approx. 1 min. **f**, Total count of M1 and S1 neurons co-recorded, and peri-stimulus time histogram aligned to trial-based movement (red line, as in **a**, which is nearly peak-velocity) for all M1 and S1 neurons recorded, with the mean estimated spiking rates shown (see Methods). Right: heatmap of all neurons recorded in S1 and M1, order ranked by activity. The pink line denotes the estimated pull-onset.

To address the limited understanding of how neurons in the forelimb S1 encode movement and the potential involvement of both S1 and M1 in encoding SPEs during motor learning, we analyzed the activity of 9,268 excitatory L2/3 neurons expressing GCaMP6s using two-photon single-cell resolution mesoscope imaging (63). We recorded simultaneously in both contralateral M1 and S1 forelimb areas of mice performing this task (n=6 mice, Figure 1a-e) from layer 2/3 excitatory neurons given the hypothesis that during learning these neurons could encode SPEs. We computed behavior-triggered (joystick movement) trial averages across all neurons (Figure 1f), which showed that both M1 and S1 had movement-related activity. Thus, we proceeded to record in M1 and S1 during a motor learning task to test the hypothesis that they encode prediction errors during motor learning.

### Learning to adapt to external perturbations

The ability to rapidly update our movements based on external perturbations is an essential task for the nervous system. Humans, monkeys, and mice can adapt to perturbations of the forelimb (1, 3, 12, 18, 20). Although the role of the sensorimotor cortex as being essential for this behavior has been established by temporary inactivation of S1 (1, 3), the underlying neural dynamics have not been shown. Thus, we adapt the delayed-onset force field task that has been successfully used in humans and mice (1, 20) and after the baseline learning period as described above, we test their ability to respond to a force field applied to the forelimb.

Within a single session, after 75 baseline (null field) trials, we applied a force field in 100 trials that perturbed their hand path orthogonal to the pull axis, whose onset was at a spatially defined point midway through the pull (Figure 2a) and active for 100 ms. This delayed-onset force field does not allow them to pull straight again after learning the perturbation dynamics (it is difficult to predict a sudden onset force versus a gradually applied force), but has the advantage that it allows them to end up in the reward zone (Figure 2a b), and that we have an exact positional onset that is highly useful for aligning with neural data post hoc. Then, after 100 such perturbation trials, in an uncued manner change back to a block of 75 null field trials (commonly called the washout epoch (12, 20)). We found that throughout the perturbation trials the mice could adapt their motor output, measured by a significant reduction in perpendicular displacement (t-test ‘first_10p’ vs. ‘late_p’ p = 0.0006, Figure 2b-d). Furthermore, the measurement of the perpendicular displacement was well fit by a state-space generative model that relied on prediction errors (see Methods, Figure 2d e).

**Figure 2.**
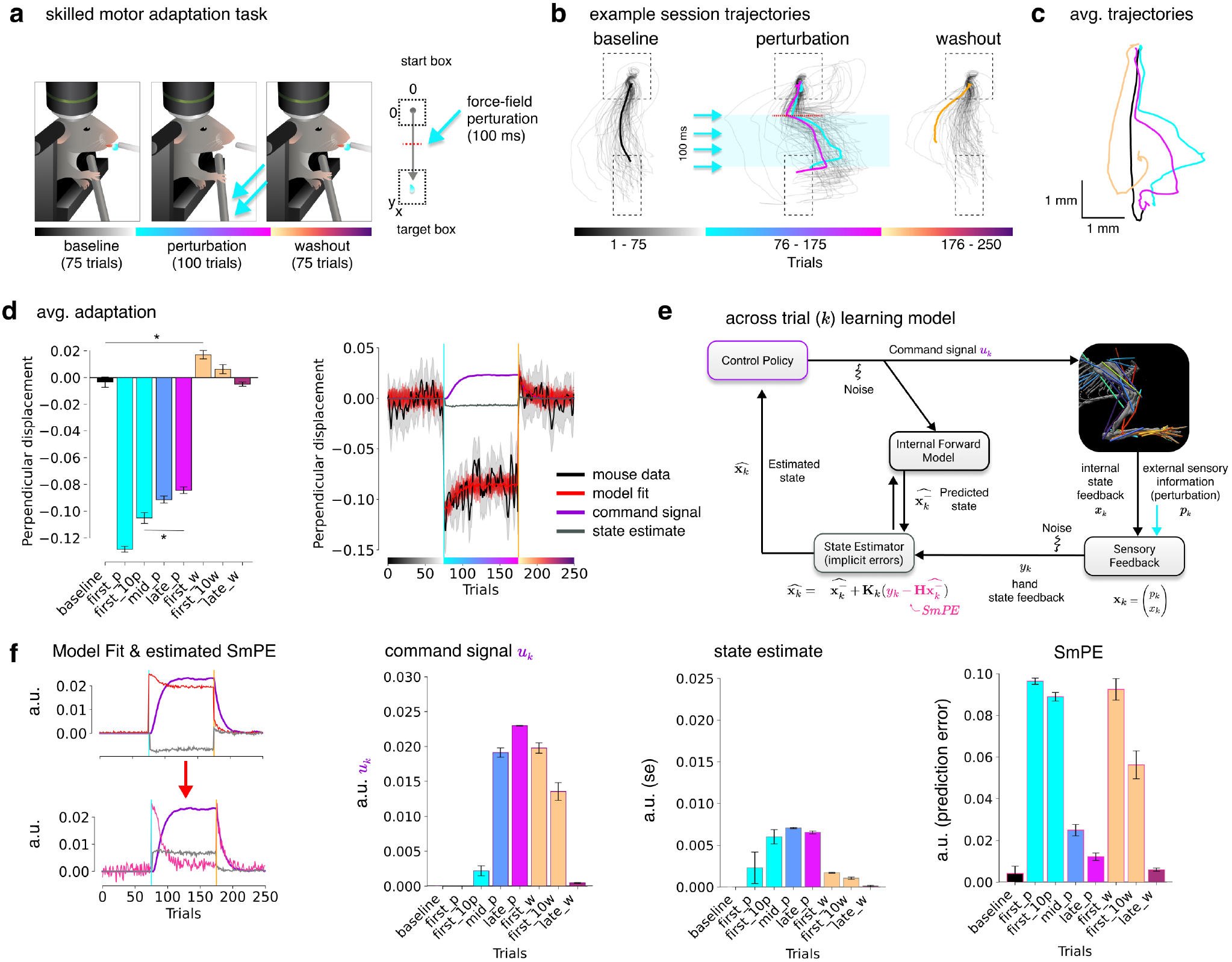
Mice learn to adapt to motor perturbations and sensorimotor cortex reflects changes in neural activity. **a**, Schematic of the experimental setting showing a head-fixed mouse engaged in a reach, grab, and pull task, where during specific trials a force field is applied to the joystick. Cartoon by Julia Kuhl. **b**, An example adaptation session showing pull trajectories; baseline average trajectory in black, avg. first 10 perturbations in cyan, last 10 in pink, avg. first 3 washout trials, and otherwise all trajectories per block are shown. **c**, Avg. trajectories across mice (n=4 mice). Black is the avg. baseline (during the last 25 trials, i.e., 50-75), cyan is the first 3 perturbations, magenta is the last 3 perturbations, and orange is the first 3 washouts. **d**, Left: bar plot of the average perpendicular displacement during the force field. Baseline is trials 50-75, first 3 perturbations, first 10 perturbations, mid-perturbations (trials 87-125), late perturbations (126-175), first 3 washout trials, first 10 washouts, and late washout (trials 230-250). Right: Perpendicular displacement across trials (n=4 mice, avg. *±* SEM), showing reduction in error during learning and an after-affect in washout. Data is smoothed (within block) with an Savitzky-Golay filter (polynomial order 3, window length 11). Median model fit is shown in red (with 10 runs in alpha), model-predicted *u*_*k*_ in purple and *x*_*k*_ in grey. **e**, Schematic of the across-trial learning model for state estimation computing internal signals across trials during adaptation (see Methods). Note, 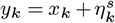 is the sensory feedback to the hand, while 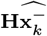 is the prediction from the internal model. **f**, Extracted model parameters based on fitting to the mouse behavioral data (the x-position of the joystick in a given spatial bin), binned as in **d**; this generates hypotheses of per-trial neural dynamics for a putative command signal, state estimator, and sensory prediction error. For interpretability, we also show how the model fit (red) gives rise to the hypothesized prediction error signal (pink). Note, the state estimator and prediction errors in the model are negative values thus, we show the absolute values. This transform is shown in the left panel for clarity.

As a secondary measure of learning, we tested if they had an aftereffect. This is where they overshoot in the opposite direction as the force field upon now entering into trials with an unexpected null field (return to baseline conditions after the learning block). This not only reveals the intended motor command (as the lack of perturbation is unpredicted), but also showed evidence of having an updated internal model (1, 12, 64). They showed a statistically significant aftereffect in the trials immediately after the force field trials, which rapidly returned to baseline levels, as expected (t-test ‘baseline’ and ‘first_w’: p = 0.0371, ‘baseline’ and ‘late_w’: p = 0.7238).

The state estimator model makes concrete predictions about latent computations that could drive learning and be reflected in the washout period (Figure 2e f). Specifically, the model makes predictions about prediction errors, a state estimator signal, and the command signal that would be required to adapt across trials; therefore, we computed these signals based on the model fit (see Methods, Figure 2f).

### Sensorimotor cortex encodes prediction errors

We asked if we could find neural signatures of these latent computations in our layer 2/3 neurons. We computed peri-stimulus time histograms (PSTH) triggered on the onset of the force field for the initial, early, middle and late perturbation trials, and for the early and late washout trials (Figure 3a). We found that the neural population during this task significantly increased their mean firing during perturbations and in washout compared to baseline firing rates (Figure 3a; t-test, baseline vs. all other bins individually tested *p <* 0.0001). Moreover, we found that compared to the baseline spiking activity, both M1 and S1 showed this signature (Suppl. Figure S1a b) as measured in the PSTH’s and as a change from baseline using the area under the curve (Suppl. Figure S1c). As a control, we considered the baseline only sessions that have the same 250 trial structure but without any perturbation (as shown in Figure 1) and found no significant changes across the session (Suppl. Figure S1d-g).

**Figure 3.**
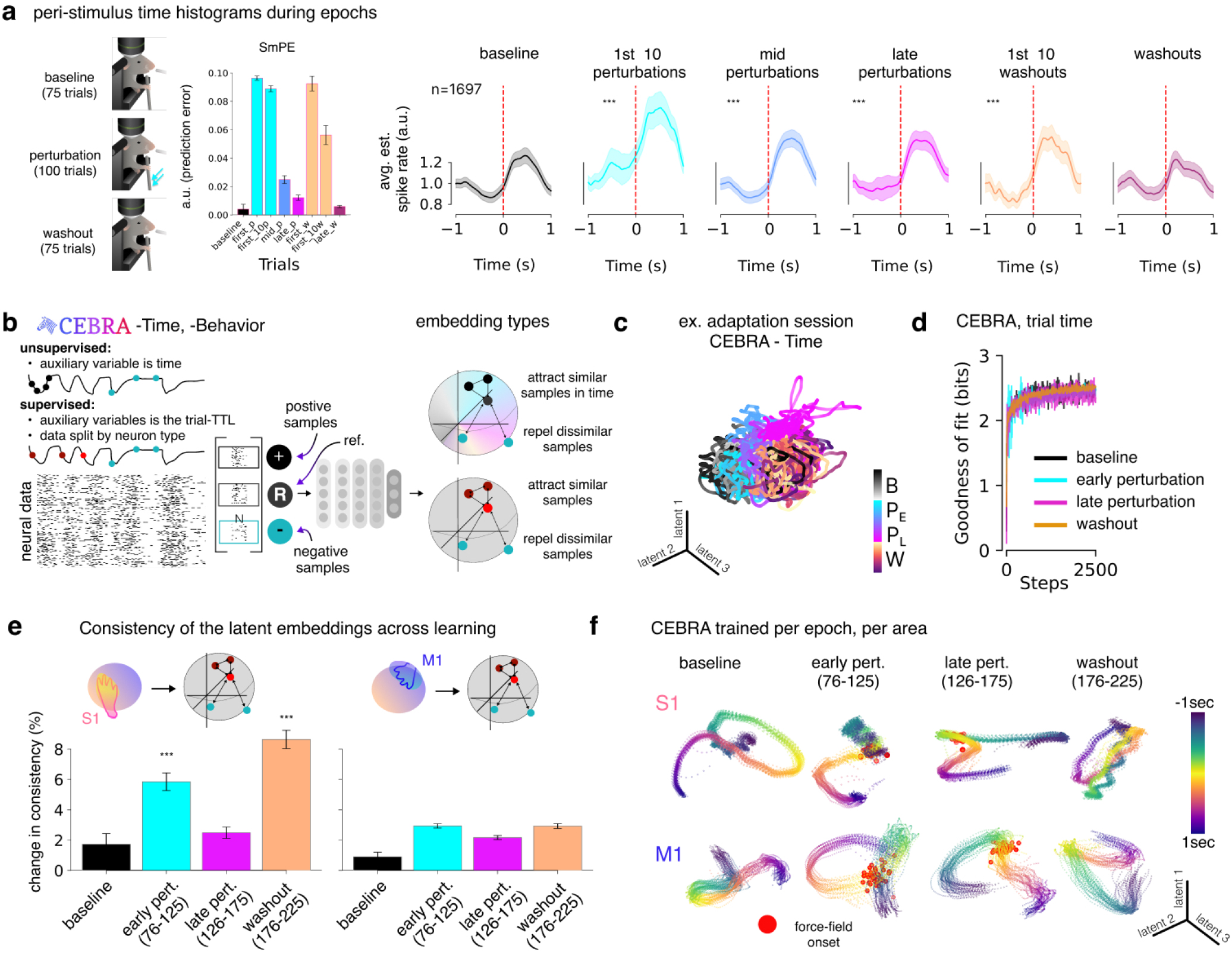
Sensorimotor cortex reflects the neural dynamics of learning to adapt. **a**, Left: diagram of the task epochs and the model estimation for sensorimotor prediction errors (from Figure 2f). Right: Normalized avg. estimated spiking rates PSTH plots over all neurons for a listed trial-bin. *** bins are significantly different than baseline *p <* 0.0001, paired t-test. The red dashed line denotes force-field onset time. The colors denotes the trials used to compute each block, as written. **b**, Diagram of how we used CEBRA in an self-supervised and supervised manner to quantify changes in the latent dynamics across learning. We use CEBRA-Time on each individual session (see c, d) then build single encoder models from across all mice in a neuron and trial dependent manner (see e, f). **c**, An example CEBRA-Time (offset-10-model-mse, 8D) colored by baseline perturbation, and washout. The first three latent variables are shown (no ranking). **d**, We trained a single encoder model per block: 50 baseline trials, 50 early perturbation, 50 late perturbation, and 50 washout, and measure the change in the latent space between blocks with a consistency metric (shown in matrix, see Methods). Goodness of fit for all models showed there was no difference in their ability to train. **e**, Quantification of CEBRA models trained per epoch and area. S1 n=908 neurons, M1 n=789 neurons; each showing the output embeddings and consistency matrices. CEBRA was run n=10 for each epoch, then the change in consistency vs. baseline across epochs was computed (mean *±* SEM shown; see statistics in Suppl. Figure S9 Table S1). **f**, Visualization of example CEBRA embeddings from **e**.

Such punitive prediction errors are key in state estimation, and therefore in the updating of internal models. This is hypothesized to be implemented as a dynamical system (1, 17, 18, and see Methods). Therefore, we leveraged CEBRA (65) to model nonlinear changes in the neural dynamics. Using self-supervised CEBRA models, we observed changes in the latent space across learning (Figure 3b c, Suppl. Figure S9a). To quantify the changes in the latent spaces, we trained individual encoder models across each epoch (Figure 3d Suppl. Figure S9b Video 3). Importantly, we could equally fit each epoch with the same number of latent dimensions, model parameters, and training time (Figure 3d), which is an important control before quantifying changes across models (i.e., with a consistency metric). Next, we measure the change in the latent dynamics between epochs.

We built M1- or S1-only CEBRA models and found that the latent space in S1 significantly changed compared to baseline in the early perturbation and washout phases (Figure 3e f; see statistics in Suppl. Figure S9c d and Suppl. Table S1). The latents returned toward baseline during late perturbation trials (despite the trajectories remaining different from baseline). Notably, encoder models trained across mice with only M1 data showed less change in the consistency of latent neural dynamics between blocks (i.e., 2% early perturbation vs. baseline), while S1 activity shifted nearly twice as much during perturbation (i.e., 4.1% early perturbation vs. baseline; Figure 3e f, Suppl. Figure S9c d). Thus, the relative change from baseline activity was significantly higher in S1 and not in M1 (Suppl. Figure S9d). This suggests that when the internal models are updated, even though baseline movements are restored in washout, the cortical dynamics remained more changed in S1.

In summary, across the sensorimotor cortex we see signatures of prediction errors, yet critically we see larger changes in S1 dynamics, which is the region known to be essential for adaptation in humans and mice. However, a central question remained: in what reference frameworks were these prediction errors computed?

### A realistic adult mouse musculoskeletal biomechanical arm model

We next aimed to test the hypothesis that the sensorimotor cortex encodes coordinate-specific prediction errors. Thus, our objective was to establish which coordinate frameworks are encoded in S1 and M1. In primates, it has been shown that both high-level (position, direction) and low-level (muscles) can be encoded in M1 and S1 (40, 66). However, in mice, while excellent work has shown neural activity is strongly correlated with EMGs from a few forelimb muscles (44), the space of high-level and low-level coding has not been established. In an ideal setting, one could record all the forelimb muscles and from each muscle we could record proprioceptors to measure feedback to S1. The recording of muscles (with EMG) has been very effective in providing important signals to better understand forelimb reaching (6, 10, 44, 66, 67), yet recording from all muscles in the arm remains technically impossible to do in any animal, and particularly challenging in mice due to their small size. Although emerging technology is being developed to record up to 8 muscles in the mouse (68), there are 52 muscles in the adult mouse forelimb that spans the trunk and head to the shoulder, arm, and hand (69, 70). Thus, here we developed a new approach for normative modeling of the sensorimotor system by developing a neuro-musculoskeletal model of the mouse in order to then extract the high-level and low-level features to compare with neural data.

Musculoskeletal models in humans have been critical for sports science, medical applications (such as modeling for prosthetics), and robotics, but are less widely used in neuroscience (71–74). Exceptions have been to model the role of proprioception (50, 75) or motor control in humans and non-human primates (76–79). Recent progress in the development of rodent biomechanical models has already shown promise (80), but a remaining gap has been the lack of a complete adult musculoskeletal forelimb.

We developed a novel model of adult mouse musculoskeletal biomechanical forelimbs by performing micro-CT scans in various postures (including reaching and grasping as in the task we use) and MRI-guided analysis to determine muscle insertion points in 50 muscles (we did not model the two tiny ‘thumb’ muscles; Figure 4a-d, Suppl. Figure S2a b; see Methods). Moreover, we developed a method using machine learning to optimize the final placement of the muscle attachment point by maximizing the amount of force that can be exerted on the arm while recapitulating movements from experimental recordings (Suppl. Figure S2c; see Methods).

**Figure 4.**
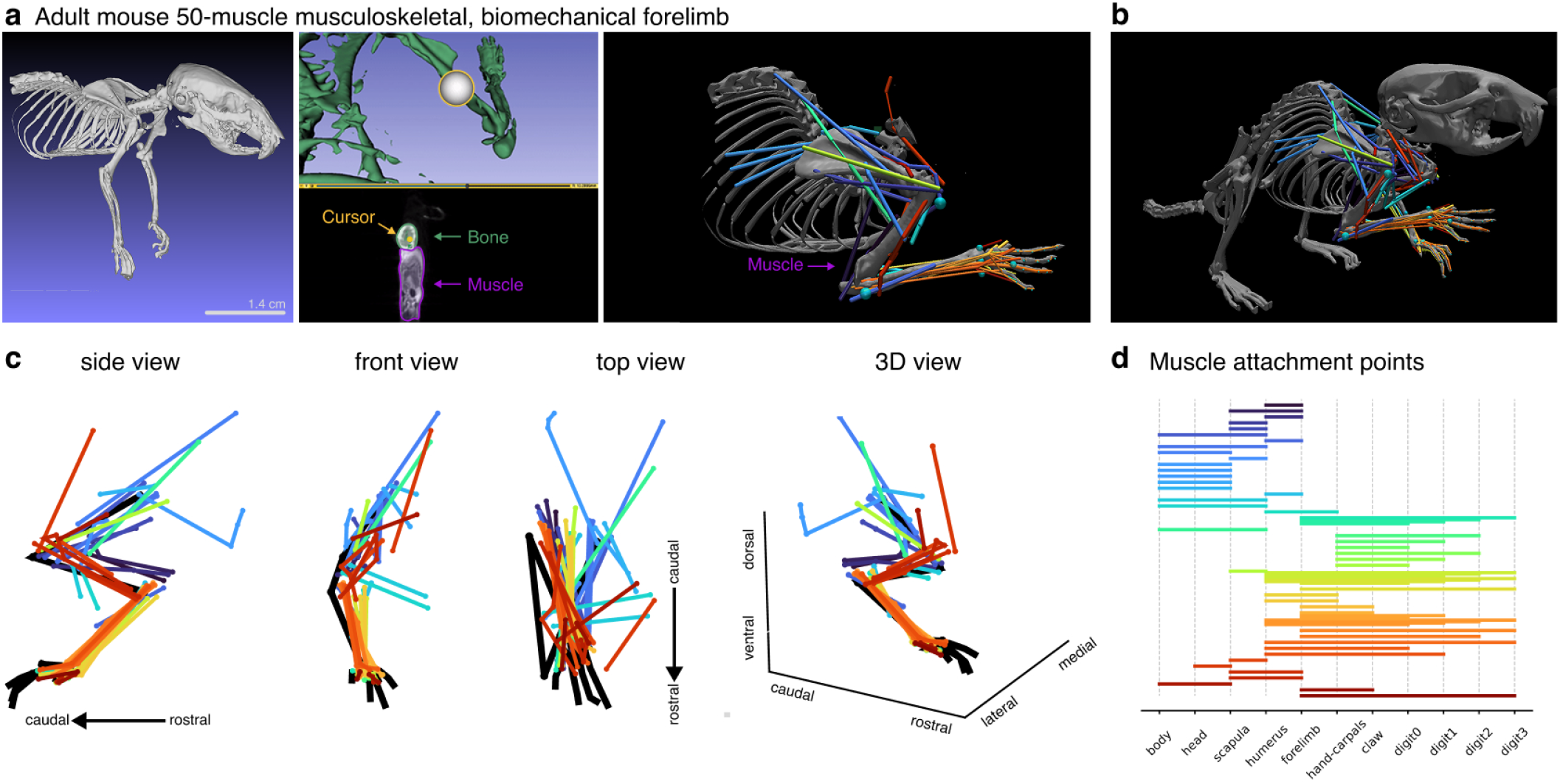
Musculoskeletal biomechanical model of the adult mouse forelimb (MusBioMaus). **a**, Development of the model: micro-CT scans were acquired in reaching, standing, and t-pose positions (see Methods). Muscle attachment (insertion) points were manually extracted from CT (right panel) and MRI scans then optimized with a neural network. In total, there are 50 muscles in the MuJoCo model. **b**, Example image of full model of the mouse musculoskeletal biomechanical arms and shoulders. **c**, The full 50-muscle arm, viewed from the side, front, and top view. **d**, Clustering of functional groups of muscles, delineating the scapula, shoulder, elbow, wrist, all digits.

Our musculoskeletal model was developed in MuJoCo (74), a physics engine designed for the efficient simulation of articulated structures interacting with the environment, plus the ability to accurately model tendon dynamics. We aimed to fully model our reach, grasp, and joystick pull task for mice. Thus, we generate 3D pose estimation data with DeepLabCut (81–83) (Video 1; see Methods) and use inverse kinematics optimizations to identify joint trajectories that best reproduced the 3D pose with our model in MuJoCo (Figure 5a Video 2).

**Figure 5.**
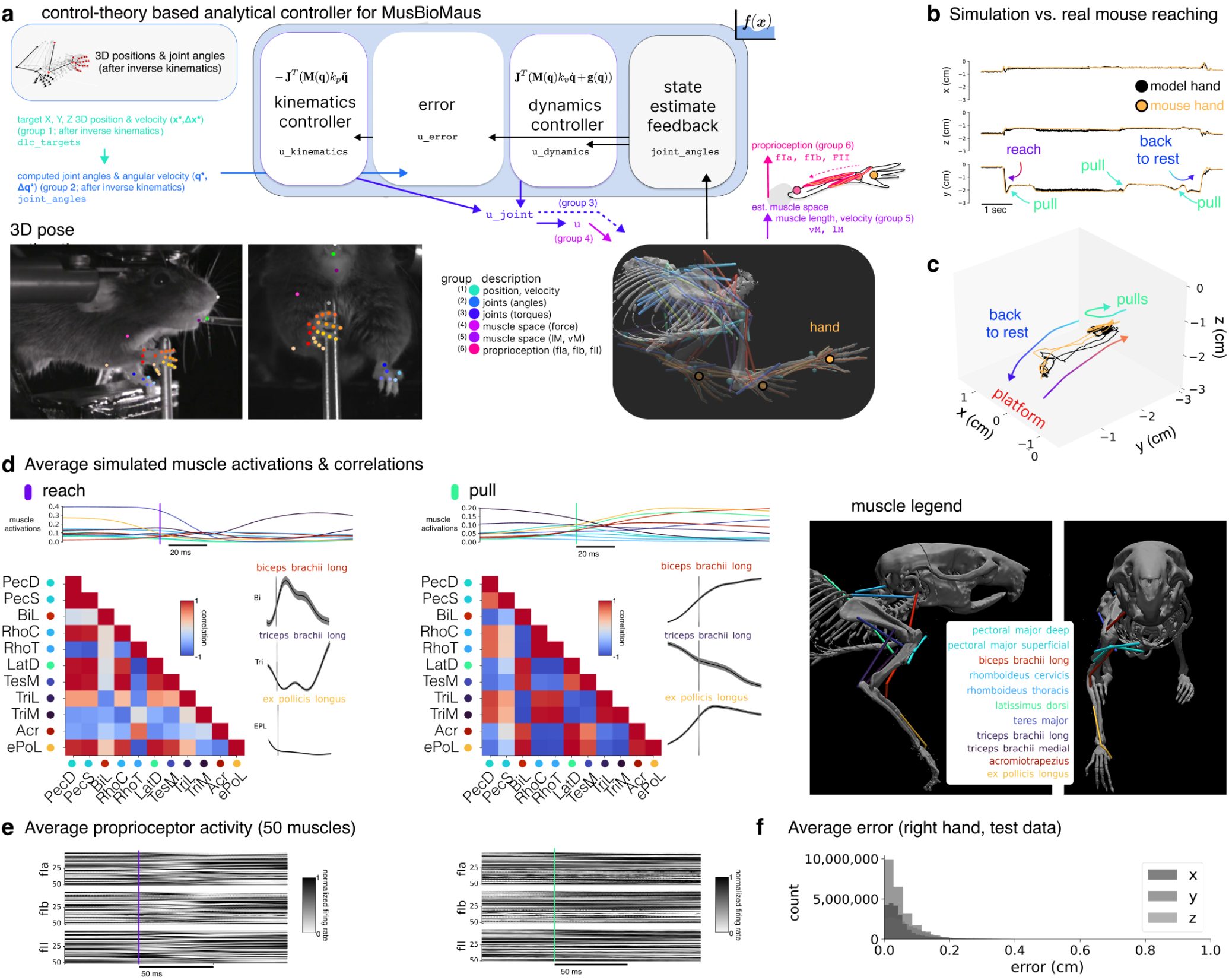
*In silico* modeling of reaching and pulling motor behaviors for mice. **a**, A diagram of the analytical controller. The kinematics and dynamics controllers, error calculation, and state estimate feedback modules directly execute the mathematical equations of their associated functions, developed to match behavioral level phenomena (see Methods). The model can be set to control joint torques and generate muscle-activations to drive the 50-muscle arm model for full musculoskeletal control. **b**, We simulate the same reach, grasp, pull task that real mice perform; here is an example test session reproducing a 6 second sequence from experimental data using our muscle-based arm controller, showing the x, y, z trajectories of the hand, as well as in **c**, a 3D plot of the action. To note, we trained our controller on only 128.86 seconds of data (the test set was therefore 27,796.05 seconds). **d**, We compute the average muscle activations in a set of anatomically-grouped muscles (key on the right) during stereotyped (left) reach (n=342) and (right) pull actions (n=435, see Methods). The top shows the average muscle activations over time. The matrix shows the correlation during this time window, and to the right is the generated “EMG” signals from our model for three muscles. All muscle results can be found in Suppl. Figures S3, S8. **e**, Muscle spindle (fIIa, fII) and GTO (fIb) outputs (from all 50 muscles) for the same time epoch as in **d** is shown. **f**, Quantification of the 3D simulated movements vs. real mice movements across all actions of the hand (n=22 sessions; 118,523,145 time steps in total).

### An analytical control-theoretic approach for animating MusBioMaus

We developed a novel analytical approach to generate torques and muscle activations that drive movements that match the behavioral data recorded in our joystick task setting (Figure 5a see Methods, n=26 sessions). Specifically, we control our muscle-based arm model by generating joint torques using a proportional-derivative (PD) controller to follow a pre-planned joint angle trajectory (while accounting for the forces of inertia and gravity), and transform these forces into muscle activations. This transformation is performed through a dot product calculation with the Jacobian between muscle space and joint space. The resulting forces exerted on the arm are the best approximation of the torques specified by the joint space PD controller, given the arm’s muscle configurations and current states, and the precise attachment points we found to be critical for reliable force control (see Methods, and Suppl. Figure S2c-e).

This approach can be considered a ‘white-box’, mechanistic approach, where the underlying computations performed by the system are well-understood, without the standard trappings of machine learning methods (Figure 5a). Critically, our approach also allows us to *in silico* record internal control signals – from kinematics to muscle-level control and sensory feedback (Figure 5b). Collectively, we refer to our musculoskeletal biomechanical mouse model and the control-theoretic analytical controller as “MusBioMaus” (**Mus**culoskeletal **Bio**mechanical **Maus**; Figure 5a Video 3).

We validated our model by computing the model vs. real-mouse trajectories and showing that they are highly similar (mean hand error in x is 0.06 cm *±*0.07 (STD); y = 0.05 cm *±*0.08, z =0.08 cm *±*0.07; for reference, the forearm length is 1.38 cm (Table 1; Figure 4a)). We computed the correlation of antagonist and agonist muscle groups based on example electromyographic (EMG) recordings in mice, rats, cats, and primates (44, 68–70, 84). Namely, we grouped the muscles (69, 70) (see Methods) and computed the correlation of co-activation during limb movements (Figure 5d Suppl. Figure S3). We also extracted the outputs of the proprioceptor models for use in downstream analyses (Figure 5e). Next, we compared mouse forelimb EMG data (44) and confirmed expected results (i.e., muscles such as biceps should contract, triceps relax during pulling; Suppl. Figure S4), but given that not all muscles have been experimentally recorded in mice, others are predictions (to note, many are very small and current EMG technology would prevent their recording).

**Table 1.** Bones of the Mouse Arm.

| Bone Name | Length (cm) | Bone Name | Length (cm) |
| --- | --- | --- | --- |
| finger0_0 | 0.190855 | finger2_0 | 0.208992 |
| finger0_1 | 0.213156 | finger2_1 | 0.192786 |
| finger0_2 | 0.13053 | finger2_2 | 0.163729 |
| finger0_3 | 0.0993316 | finger2_3 | 0.073004 |
| finger1_0 | 0.222189 | finger3_0 | 0.140258 |
| finger1_1 | 0.251804 | finger3_1 | 0.149808 |
| finger1_2 | 0.137059 | finger3_2 | 0.113512 |
| finger1_3 | 0.0794417 | finger3_3 | 0.0701662 |
| metacarpals | 0.185991 | thumb_0 | 0.0777345 |
| forearm (ulna + radius) | 1.3834 | thumb_1 | 0.0984265 |
| humerus | 1.24144 | scapula | 1.19748 |

### Neural population dynamics encode 3D kinematics and muscle-level features

Next, in order to establish which high-level and low-level features are represented in M1 and S1 in mice, we first examined the tuning properties of individual neurons with Generalized Linear Models (GLMs, Figure 6a-d and Suppl. Figure S5a-e; see Methods) that included the task-relevant, kinetic, and kinematic features. Given that kinetic or kinematic features can be correlated at different points in the behavior (Suppl. Figures S6 S7), we thematically grouped the regressors to measure their relative effects. Specifically, we built a global design matrix of groups that captured the 3D position & velocity (group 1), joint position & velocity (group 2), joint torques (group 3), muscle-space activations (group 4), muscle length & velocity (group 5), and proprioceptive signals (muscle spindles & Golgi tendon organs, group 6).

**Figure 6.**
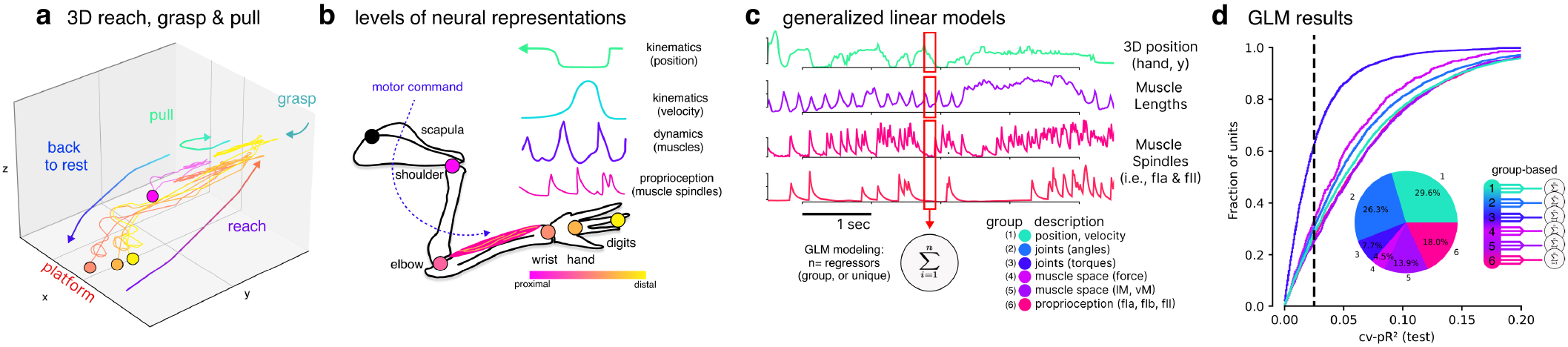
Neurons reflect posture and muscle-level features. **a**, 3D plot of an example reach, grasp, and pull (and retraction to platform) by a mouse, highlighting the wrist, hand, and finger (colors as in **b**). **b**, Our 3D data allows us to extract kinematics and dynamics across joints (and with our model mouse, muscles). **c**, Diagram of the GLMs design matrix (with only example regressors such as hand, muscle length, and spindles) for measuring unique contributions and grouped by major levels (see Table 6 for full description). **d**, Cumulative variance explained vs. pseudo-*R*^2^ for all neurons, showing our cutoff of 0.025 for downstream analysis. Inset pie chart shows the percentage of neurons tuned for a specific level (where any score over 0.025 is counted for that level and then we normalize to equal 100%).

We simulated each real-mouse session with MusBioMaus to record 3D kinematics, dynamical features of internal muscle control, and proprioceptive feedback in the same *in silico* framework. For computed proprioceptive features we used models for two spindle types (fIa, fII) and one for Golgi tendon organs (fIb, see Methods). For GLMs, we computed a cross-validated pseudo-*R*^2^ and only further considered neurons with performance over 0.025 (Suppl. Figure S5a-c), which accounted for 55.4% of the population (Figure 6d).

The GLMs show that both high-level egocentric 3D position and lower-level proprioception and the muscles that support the movements of the forearm and hand were well represented in the sensorimotor cortex (Suppl. Figure S5e). This suggests that the forelimb area (fS1 and part of M1 (85)) in mice encode both high-level postural information, muscle, and related proprioceptive features, similar to Brodmann’s area 3a in primates (40, 86). These signals would be essential for effective state estimation – knowing where in 3D space one’s body is, plus knowing the muscle-space coordinates that enable movement.

### Computational motifs map to neural ensembles

The neural dynamics of the sensorimotor cortex richly encodes high-level kinematics and low-level muscle kinematic features that are hypothesized to be critical signals related to an internal model of the world. We therefore aimed to map computational motifs – prediction errors, state estimation, and command signals – to the neural dynamics and compute the related coordinate frameworks. This is only possible given our neuro-musculoskeletal model-guided analysis.

We used a data-driven approach to compute PSTH’s for each neuron across a set of binned trials across the session, then used this per-neuron signature to perform hierarchical clustering in order to quantify how neurons changed across learning (Figure 7a). Specifically, we performed principal component analysis (PCA) followed by hierarchical clustering (Figure 7a), which identified 5 clusters that was significantly different from random clustering (Adjusted Rand Score, n = 10,000 permutations, p = 0.00). We observed that the sensorimotor prediction error^0^ dynamics, which showed an increase in firing during the early perturbation, a decrease over learning, and again a change in firing during washout, was in two clusters (cluster 0 and 3; Figure 7b). These clusters represented 22.5% of the neurons recorded (n = 382 of 1697, Figure 7). We also noted that two other clusters could be mapped to a command-like (*u*) signal – which presents as an increase over learning and a decrease in later washout (clusters 1 and 4 Figure 7b n = 104 neurons, 6.1%). As an additional control, if we clustered baseline-only sessions with this unsupervised method, we did not find these computational motifs (Suppl. Figure S10a).

**Figure 7.**
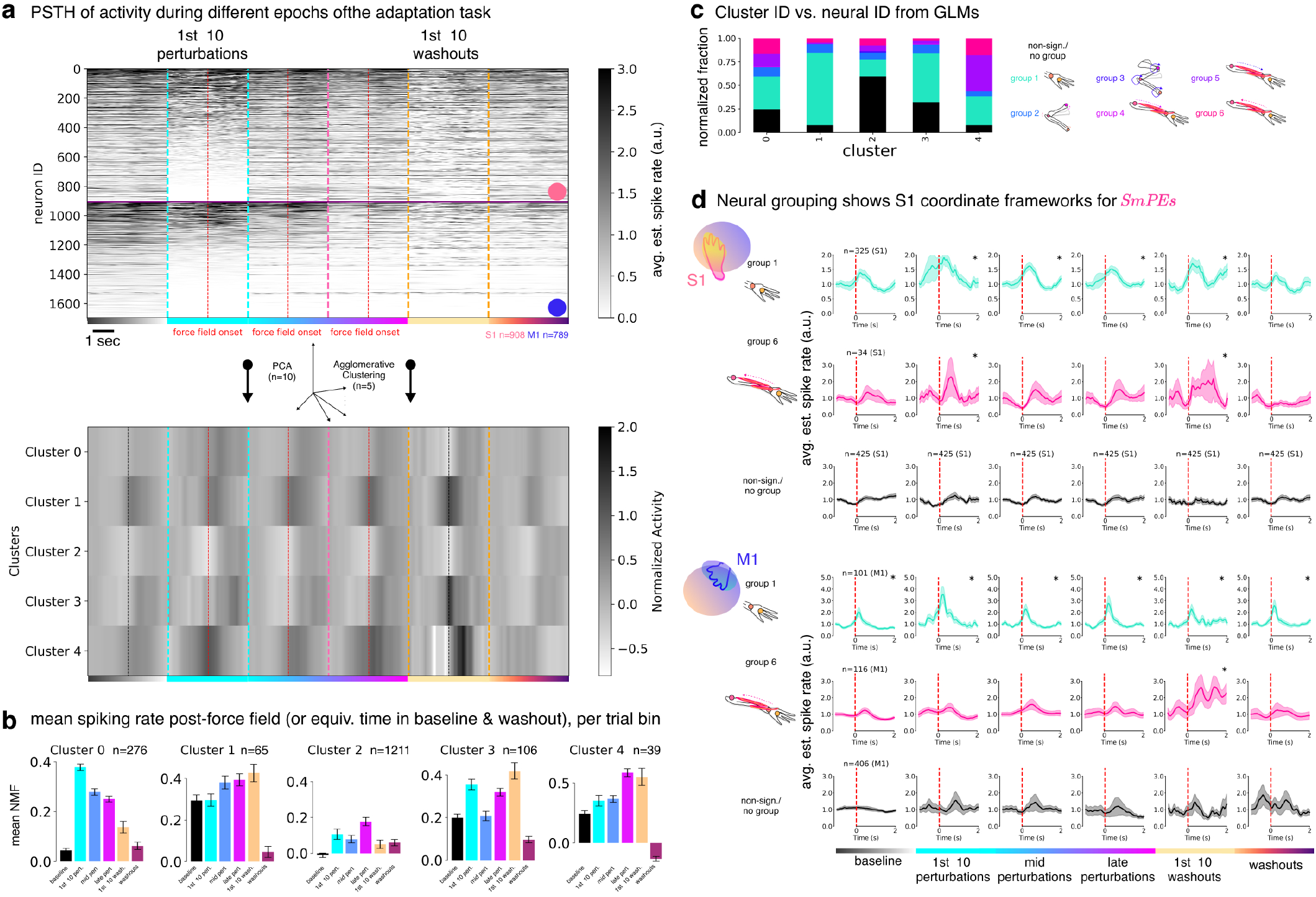
Neural populations reflect computational motifs that can be mapped onto specific neural types. **a**, PSTHs of neurons across each epoch is shown for −1 to 2 sec around trial start/force field onset (red line). The colored dashed lines denote the block change. Then, we performed clustering of these per neuron PSTH across blocks with PCA (n=10) followed by hierarchical clustering (n=5). We computed the adjusted rand score across 10,000 permutations and find this clustering is significantly different than random (p=0.0). **b**, We computed the mean estimated spiking rates after the force-field onset (red line) or the equivalent bin in baseline and washout (black line) per bin for each of the 5 clusters. The number of neurons per cluster is noted. Clusters are organized by grouping: 0 and 3 show a SPE signature, 2 none, and 1 and 4 show a command-like signature. To see the clustering PSTHs steps, see Suppl. Figure S10. **c**, Using the GLM-based neuron ID we plot the normalized count per cluster with the relative proportion of each group. **d**, Using the neural grouping, we re-compute the PSTH per bin (as noted) for S1 (top) and M1 (bottom). We computed the mean estimated spiking rates and SEM per time bin. *denotes a significant difference across multiple timesteps using a permutation test of baseline (using all neurons) vs. each combination of block:region:group 1 or 6. Full statistics are available in Suppl. Tables S2, S3.

Lastly, to ask which coordinate framework these neurons were in, we assigned each neuron a group ID. While this does not imply that a neuron exclusively represents a single group, it enables us to ask whether a functionally defined neuron type preferentially maps to the SPE signature. In the SPE clusters (0 and 3), we note that the neurons assigned showed a mixture of tuning (neural ID was assigned based on the GLM analysis), but notably group 1 is most represented (Figure 7c). If we clustered all neurons with the same group ID, we find that group 1 and group 6 reflect the signature SPE in S1, but this was not in M1 neurons (Figure 7d Suppl. Tables S2, S3, and see Suppl. Figure S10c for the other groups). Thus, while sensorimotor prediction errors could be computed in multiple reference frameworks, the data suggest it is enriched for 3D kinematics and muscle-features (Figure 7d).

Taken together, our analyzes show that the neural dynamics in S1 changes more strongly than M1, and S1 encodes sensorimotor prediction errors. This could mean that neurons within S1 layer 2/3 are using sensorimotor prediction errors, primarily in 3D kinematics and muscle-feature space, to build an optimal state estimator that can be used for adaptive learning.

## Discussion

Understanding how neural dynamics enable adaptive, dynamic control of the body is a fundamental goal of neuroscience, and has implications for improving applications in robotics, artificial intelligence, and neuroprosthetics. We report how sensori-motor cortex supports skilled motor control and learning by encoding multi-dimensional representations of kinematics (3D position and joints) and dynamics (muscular and proprioceptive) information. We focused on recording in S1 from excitatory layer 2/3 pyramidal neurons, which are postulated to be critical integrators of top-down (pre-motor) and bottom-up (thalamic, brainstem) inputs (25), namely motor efference and proprioceptive feedback, and due to S1 having a known essential role for the ability to adapt in mice (1) and humans (2).

Firstly, we find that the sensorimotor cortex shows the signature of sensorimotor prediction errors: an increase in firing when an unexpected perturbation occurs that reduces over learning, then a change in firing again during the unexpected removal of the force field. This was measured by both classical PSTHs, and with nonlinear machine learning-based encoding models (Figures 2a e, f). Interestingly, we find that S1 changes more than M1 (Figures 2e f). We then aimed to study two aspects of these dynamics; first, how they computationally could encode hypothesized signals used in optimal feedback control (Figure 2 and secondly, what frameworks these neurons might be computing in (Figure 6.

To map neural dynamics to optimal feedback control, we inferred relevant variables from joystick trajectories (behavior) with a generative state estimator model (Figure 2d-f). This allowed us to get a trial-by-trial estimate of the command signal, the state estimator, and the sensorimotor prediction error. Then, we computed PSTHs for all neurons across trials, binned them into appropriate epochs, and performed unsupervised hierarchical clustering. The five resulting clusters showed signatures that resembled the command signal (6.1% of neurons) and a sensorimotor prediction error (22.5% of neurons; Figure 7a b). This bias towards more SPE-like clusters agrees with the general overall SPE signature that is reflected in the PSTHs and in the neural dynamics as measured by nonlinear encoding models. Thus, we found that functional ensembles of neurons can be mapped to computational motifs that support optimal feedback control.

In order to better understand the coordinate frameworks that these neural motifs reflect – i.e., 3D posture to muscle spaces – we needed to develop an anatomically accurate articulated adult mouse arm model. Prior efforts have used estimates from embryonic mice (80) or did not include muscles (87). Here we combined CT scans, MRI in posed arm configurations, and a neural network-based optimization method (see Methods) to ensure we captured movements that were similar to our experimental conditions, then finally animated the mouse arm in a physics engine. This allowed us to assign neurons to various features important for motor control in this task.

Interestingly, when computing PSTHs using assigned neural groups, we find that S1 neurons encoding both high-level (3D kinematics) and low-level proprioceptive (muscle spindle) signals exhibit signatures of sensorimotor prediction errors during motor adaptation. Why might they be in both coordinate frameworks? We believe that this combination could be the essential ingredients for state estimation across spatial scales. The state estimator is the combination of the current internal model 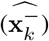 plus the uncertainty-weighted actual vs. predicted feedback 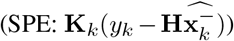. We hypothesize that proprioceptive-tuned populations may be used more for fine motor control, while high-level 3D kinematics is less positionally refined but critical for detecting global, task-relevant trajectory deviations. Thus, having both low-and high-level represented in S1 could be advantageous for spatial refinement, and also why repeated exposure to perturbations is needed to refine the motor plan. Analogous to how the cerebellum and basal ganglia mediate fast and slow learning (88), S1 may resolve spatial error components, supporting accurate feedback to the controller (putatively M1).

Previous work has shown that S1 responds to peripheral sensory stimulation before M1 and the cerebellar cortex (89, 90), which would support the idea that S1 could compute the state estimation that is essential for internal forward model-based control, where signals from S1 are then sent to M1 and the cerebellum. The role of S1 and the parietal cortex (area A5) in primates was shown to be crucial for volitional limb control, and inactivation of A5 suggested that it likely housed a state estimator (52, 91). These finds are complementary to finding that the neural dynamics reflect sensorimotor prediction errors and state estimation in S1 (note that mice do not have an equivalent area to A5 that projects back to the spinal cord, and a close homologue might be S1). This also synergistically adds support for the underlying neural basis of prior work, which found that S1 was essential for learning to adapt (1, 2).

Another interesting observation is that S1 changes more than M1 during adaptation. Why might M1 change less despite its role in adaptation? Prior work on potent vs. null spaces suggests that local changes on the neural manifold can produce large behavioral effects (67). Thus, we hypothesize that even small changes in M1 could drive behavioral output if they occur in the potent space (67, 92, 93). This could also explain why, although less strong, we do see significant changes in M1 throughout adaptation. A small change in M1 was reported with different methods in primates during motor adaptation (4, 94). Providing another perspective is also the finding that the tangling in M1 is less than other areas such as SMA (95) or S1 (96). Notably, the degree of tangling in S1 was closer to EMG signals than M1 (96). Collectively, if the key change in neural activity during our adaptation task is how S1 computes a change in state based on changes in sensory feedback (namely, SPEs) this change in computation would be reflected in the neural activity, while M1 can continue to perform a similar computation (i.e., an adapted command signal) which could result in more subtle changes in the population dynamics.

Lastly, while this task focused on forelimb adaptation, an important future direction will be to extend this model to other tasks that include whole body movements to test the generalizability of the sensorimotor prediction errors. Another important direction is to causally perturb these precise neural ensembles during behavior in order to test their ability to drive adaptation. Ongoing efforts for real-time neural encoder-decoders that would allow for subspace identification of specific neurons in real-time will greatly aid in these important future directions (97, 98).

In summary, our work provides a theory-driven computational framework for studying motor adaptation. We used our musculoskeletal biomechanical mouse model to tackle a long-standing question about the nature of neural representations in the sensorimotor cortex during learning. We were able to find both kinematic and muscle-level features with both classical methods and new neural dynamical systems approaches, and show that they can be mapped to computational functional motifs – i.e., sensorimotor prediction errors – in S1 that could directly support motor control.

## Contributions

T.D. developed the MuJoCo musculoskeletal model and optimization pipeline, built the MusBioMaus controller, analyzed the biomechanics data, and contributed to editing the manuscript. St.S. Built additional neural data analysis tools and analyzed the data (GLMs and CEBRA), and contributed to editing the manuscript. P.S. Prepared and collected all the CT images, parsed the biomechanical skeleton, and analyzed muscle attachment data. A.H. Built the initial DataJoint data analysis pipeline and performed preliminary neural data analysis. P.M. contributed to GLM neural data analysis and interpretation of the results. M.W.M. Conceived the project, performed surgeries, mouse training, conducted 2P imaging, developed the DeepLabCut models, assisted in all data analysis, wrote the manuscript, co-prepared visualizations, acquired funding, and supervised the project. All authors contributed to visualizations, writing the methods, and approved the final manuscript.

## Acknowledgments

This was a project that spanned over many years in the Mathis Lab (since 2017), thus we are deeply grateful to all former and current members from Harvard and EPFL for their inputs. We greatly thank M. Black for providing the resources for the CT and MRI scans, S. Zuffi & S. Pujades for discussions on CT scans, and A. Mathis for ample discussions on motor control. We also thank T. Nath for helping developing the original DataJoint pipeline, M. Popova and N. Poulsen with numerical optimizations, M. Frey and W. Monteith for 3D GT labeling and assistance with 3D pose estimation, K. Sandbrink for early GLM analysis, S. Ye for reviewing and improving the mouse controller code base, S. Hausmann for rig CAD files, and E. Madsen at the Rowland Institute Machine Shop for the customized joystick handle. We thank A. Mathis, S. Scott, A. Tolias, M. Churchland, A. Perez Rotondo, A. Schneider, and S. Bowles for discussions and/or feedback on the manuscript.

## Materials & Correspondence

For all requests please contact the corresponding author.

## Data Availability

Two-photon and related behavioral data will be released with the publication of the paper.

## Code Availability

The MuJoCo-based biomechanical model, code, and analysis tools will be available upon publication at: https://github.com/AdaptiveMotorControlLab/MuJoCo-MusBioMaus. Code to reproduce the figures will be available at: https://github.com/AdaptiveMotorControlLab/DeWolf_Schneider_etal_2025. All other requests should be made to the corresponding author, Mackenzie W. Mathis.

## Declaration of generative AI and AI-assisted technologies in the writing process

During the preparation of this work the author(s) used GPT4o(-mini) in order to check grammar, format latex tables, and (within cursor.ai) to generate and review some code. After using this tool/service, the author(s) reviewed and edited the content as needed and take(s) full responsibility for the content of the publication.

## Methods

### Animals & Surgery

Data was collected from 6 adult (P90-P365) male and female from layer 2/3 neurons that were in mice that were either slc17a7 crossed to ai148 or Thy1-GCaMP6s-4.3 mice (Jackson Laboratory). Surgeries were performed under aseptic conditions with animals under isoflurane (1-2% at 1.0 L/min oxygen) anesthesia, and pre-operatively mice were given an anti-inflammatory (dexamethasone, 5 mg/kg, S.Q.). Buprenorphine (0.1 mg/kg, I.P. every 6-8 hours) was administered postoperatively for 48 hours. Mice were surgically implanted with a custom head plate (1) and a “crystal skull” glass cranial window (99). All surgical and experimental procedures were in accordance with the National Institutes of Health Guide for the Care and Use of Laboratory Animals and approved by the Harvard Institutional Animal Care Use Committee and/or Caton Geneva, under license number GE68.

All mice were housed on an inverted 12h dark/12h light cycle (dark from 09:00 AM - 09:00 PM) and each performed the behavioral task at approximately the same time of day. After surgery mice were individually housed and allowed to recover for 7 days. All mice were maintained above 90% of their starting weight once on water restriction. No mouse was given less than 1 mL of water per day. Only right-handed mice were used in this study.

### Joystick task for mice

Mice were water restricted and taught to manipulate a 2D of freedom joystick as described in Mathis et al (1). To note, the mice presented in this work are not the same mice previously used. In brief, we utilized a joystick system (https://github.com/AdaptiveMotorControlLab/JoystickControlSystem) and rewarded mice with a approx. 3 uL drop of water for pulling the joystick towards themselves 6-9 mm into a virtual target box (of 2 mm in width). Note, they grasped the joystick with their hand in a vertical fashion, not as a pulled lever (see Figure 4a). The training progressed in three stages, where at first if the mice moved the joystick towards them any distance they were rewarded (this persisted for two to four days until they could do this consistently). Then, in stage two, and in a fully automated fashion, if they pulled 6 mm into a virtual target box and waited 100 ms they were rewarded. Once the mice could do this for greater than 100 trials at 80% accuracy for at least two days in a row the task shifted to phase three where they needed to pull 10 mm into the target box (width 2 mm). This was considered as the full baseline task. The joystick system records the position of the joystick at 1kHz within LabView and is synchronized with the video and mesoscope imaging for post-hoc analysis. We collected twenty-two baseline sessions and four adaptation sessions, where the mice where perturbed with a systemically applied force-field (see below) during two-photon imaging (see below). In six baseline sessions we included randomly applied force-field on 10% of the trials to increase the coverage of the x-y joystick space, i.e., in only 150 trials of the 4,667 total baseline trials in the data set).

The adaptation session paradigm was adopted from Mathis et al. (1). In brief, after the mice are experts at the baseline task above, we challenge them in a single session with a delayed-onset force-field perturbation. The force-field is an approximately 0.2N force orthogonal (x) to the main pull axis (y, see Figures 1b and 2a). This was only applied in trials 76-175. Other trials (1-75 and 176-250) had no force-field.

### Pose estimation

To measure the pose and kinematics of the mice we used two Imaging Source cameras 640×480 pixels recorded at 75-150Hz placed in stereo-view around the mouse’s right arm and head. We then trained pose estimation models for either the front and side view (or a single combined view model) using DeepLabCut version 2.0.7 or 2.0.9 (81, 82) (https://github.com/DeepLabCut/DeepLabCut). In brief, we labeled 32 keypoints across approx. 500 2D frames (then 95% were used for training). We used a ResNet50-based neural network with default augmentation and training parameters for 1.2M training iterations. We then used a p-cutoff of 0.9 to condition the x,y coordinates for 3D triangulation. This network was then used to analyze videos from similar experimental settings. To filter any noisy 3D triangulated data, we additionally used a manual labeling step to obtain ground truth 3D information for a subset of approx. 500 frames.

### Lifting model for transforming 2D to 3D space

We trained a simple transformer model that took as input the 2D DeepLab-Cut extracted coordinates from *one* camera and extracts the 3D coordinates through a linear output layer (83, 100). The input to the transformer consists of sequences (of length T=2 frames) of 2D coordinates representing the keypoint positions. Each keypoint is represented by an x and y position. We then encode the flattened joint coordinates using a transformer layer and then project the encoder output to a three-dimensional space using a fully connected layer. We trained this with a triangulation loss, continuity loss, and ground truth loss. For more details see Frey et al. (83) (https://github.com/AdaptiveMotorControlLab/mousearmtransformer). The transformer model is trained to output 3D coordinates in the MuJoCo reference frame, using the linear transformation parameters identified by aligning keypoints on the experimental platform (see below). In this reference frame, the origin (0, 0, 0) is located as the base of the head. We trained the model across 10 sessions where two cameras are available, and used the resulting weights to generate 3D predictions for sessions with only one camera.

### Two-Photon imaging and analysis

To record neural activity across several brain regions a commercially available two-photon mesoscope was used (2P-RAM system, Thorlabs). This microscope is designed to scan multiple fields at arbitrary locations within a 5 × 5 mm field of view and depths up to 1 mm (63). In brief, a resonant scanner provides fast line scanning (24 kHz line rate) over a range of 0.6 mm in the fast scanning direction, while the position of the scan field and movement along the slow scan axis is controlled by 3 galvo scanners. Fast scanning of different depths is achieved by a remote focusing unit, which shifts the focal point in z-direction by moving a lightweight mirror on a voice coil. High instantaneous photon densities to drive two-photon absorption are provided by a pulsed Ti:Sapphire laser with a tunable wavelength between 700 and 1100 nm (Tiberius laser, Thorlabs). All recordings were acquired using wavelengths between 960 and 970 nm and with laser powers below 60 mW after the objective.

The locations of the scan fields in the sensorimotor cortex were selected in the microscope control software (ScanImage 2018-2021, Vidrio, Alexandria, VA, USA) based on relative coordinates from Bregma (centered over +0 mm Bregma, −2 mm lateral, as defined in Mathis et al. (1), Tennant et al. (56), and the Allen Brain Atlas). Bregma was located in each recording by matching the vessel pattern visible in the full 5 mm field of view to the bregma location relative to the vessel pattern during the cranial window surgery. Depending on the size of the scanned fields, scanning rates varied between 5.7 to 16.2 Hz.

All post-recording analysis code was managed within a relational database using DataJoint (101). To extract neural signals from the acquired two-photon scans, a publicly available template for a two-photon processing pipeline from the Tolias Lab (https://github.com/cajal/pipeline) was adapted (102). The processing steps from the recorded image stacks to neural activity traces include raster correction, motion correction, segmentation, mask classification, and deconvolution. To correct for possible shifts of pixels scanned in the forward and backward direction of the resonant scanner, raster correction was performed. The shift was computed by shifting even and odd lines of the scan with respect to each other and calculating the position of maximal correlation between neighboring lines. All detected shifts were below 1 pixel due to the manual raster correction performed in the microscope control software (ScanImage 2018-2021, Vidrio, Alexandria, VA, USA).

All recorded frames were motion corrected for each scan field separately using a Discrete Fourier Transform (DFT) algorithm to register each frame to a template by x-y shifting (103). The template for each field was created by Anscombe transforming, averaging, and smoothing (Gaussian filter, σ = 0.7 pixel) a section of 2000 frames from the middle of the scan.

The x- and y-shifts were determined by finding the maximum of the convolution between each frame and the template. Shifts larger than 20 *µm* were considered outliers and were linearly interpolated. Then, the x- and y-shifts were median filtered (kernel width: 3) to remove spurious jumps of x- or y-shifts on single frames. Datasets with excessive z movement were excluded from further analysis. To detect the localized neuronal signal sources from the whole recording, the image stack was segmented using the CaImAn package (104) (https://github.com/flatironinstitute/CaImAn). The algorithm computes a constrained non-negative matrix factorization (CNMF) to extract the time course of localized sources (105). The recorded data **Y** ∈ ℝ^*d×T*^ (*d*: number of pixels, *T* : time) was decomposed into three parts:

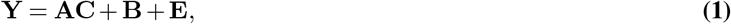

with the matrix **A** ∈ ℝ^*d×n*^ containing *n* extracted localized masks and the matrix **C** ∈ ℝ^*n×T*^ representing the corresponding temporal components. **B** ∈ ℝ^*d×T*^ is the extracted background activity and **E** ∈ ℝ^*d×T*^ is the residual activity that cannot be decomposed (104).

After segmentation, the mask shapes were evaluated with a convolutional neural network. Only shapes classified as a soma (p > 0.75) were further considered for analysis. The masks were then manually reviewed and rare edge artifacts that were classified as somas were removed. In total, we collected 9,268 neurons: 4,370 from M1 and 4,898 S1 from n=6 mice. Neuron ID was assigned based on the visible imaging window in relation to bregma using the CCF atlas.

To normalize the recorded fluorescent signal, the relative fluorescence change Δ*F/F* was calculated. The background fluorescence *F* for each neuron was estimated by median filtering (kernel size 120 seconds, padding with median values at the edges) the time course of the background signal extracted by the CNMF segmentation at the location of the neuron mask. The fluorescent increase Δ*F* was the extracted signal time course from the CNMF algorithm.

To extract estimates of spiking activity, all signals used for analysis were deconvolved by fitting an autoregressive process to the recorded fluorescent signal using the CaImAn pipeline (105). This deconvolution algorithm estimates the underlying neural activity that best explains the recorded fluorescent signal based on the calcium indicator dynamics. The assumption of this model is that the number of currently fluorescent molecules *c*(*t*) is generated by a number of spikes *s*(*t*) at times *t* paired with a decay of the signal *c*(*t*) = *γ*_1_*c*(*t* − 1) + *γ*_2_*c*(*t* − 2) + *s*(*t*) with the decay constants *γ*_1_ and *γ*_2_. The recorded fluorescent signal *y*(*t*) results from *c*(*t*) according to *y*(*t*) = *α*(*c*(*t*) + *b*) + ϵ (*t*) with a non-negative scalar *α* to scale the signal, the baseline fluorescence *b* and zero mean Gaussian noise *E*(*t*) (105). We used the output of this step as the estimated spiking rate.

### Building a functional, biologically realistic muscle-based mouse arm model

#### Building an anatomically accurate forelimb model

Following the modeling process of (106, 107), contrast-enhanced micro-computed tomography (CT) scanning was used to create a 3D model of the mouse’s skeletal structure, which was then separated digitally into component parts. Eight mice were imaged using an Inveon Multimodality CT scanner using the following parameters: 360 projects in 1° steps, 80 kV peak tube voltage, 500 µA per projection, exposure time of 250 ms, and a resultant effective pixel size of 103.80 µm. The reconstruction of the CT scans was performed using the ordered subset expectation maximization (108) (Cobra version 6.3.39; Exxim Computing Corporation) algorithm with 10 iterations, slight noise reduction, and no filter or down-sampling. The reconstructed voxel size was 103.80 µm^3^. Segmentation of the mouse skeleton was performed in 3D Slicer (109) using a threshold function applied to the reconstructed DICOM files. For all mice, a minimum value of 450 Hounsfield Unit (HU) was set for segmenting the skeleton and then manually fine-tuned.

The model was divided into 23 bone segments to allow forelimb articulation around 7 major joints (Tables 1 and 2). Specifically, the model was segmented into the main body, scapula, humerus, radius-ulna (treated as a single segment), carpals, and separate digit components. These constituent parts were then reassembled to form the arm model in MuJoCo, using the xml-based MJCF API. A slide joint and two hinge joints connect the scapula to the main body. The slide joint is aligned to allow movement along the proximal-distal axis, and the hinge joints allow posterior-anterior tilting and medial-lateral rotation. A ball joint is used for the shoulder joint connecting the scapula to the humerus, and a ball joint is used for the elbow joint connecting the humerus to the radius-ulna. There are two joints at the wrist, allowing for flexion-extension and abduction-adduction, and each of the finger joints are modeled using single hinge joints that allows flexion-extension.

**Table 2.** Mouse Arm Joint Parameters List.

| Joint | Joint Degree-of-Freedom | Joint Range-of-Motion |
| --- | --- | --- |
| Scapula1 | 1 (slide) | -0.4 to 0.1 cm |
| Scapula2 | 1 (hinge) | -8° to 17° |
| Scapula3 | 1 (hinge) | 0° to 6° |
| Shoulder | 3 (ball) | – |
| Elbow | 3 (ball) | – |
| Wrist 1 | 1 (hinge) | -115° to 115° |
| Wrist 2 | 1 (hinge) | -40° to 40° |
| Fingers 1 - 4 | 1 each (hinge) | -6° to 42° |

In total there are 50 muscles defined in the model (Table 3), representing nearly all of the muscles in the mouse shoulder and arm and including all of the principle muscles affecting movement of the scapula (69, 70). The muscle origin and insertion points, which were defined as single points, were determined on the resulting 3D skeletons based on descriptions and diagrams from (69, 70).

**Table 3.**
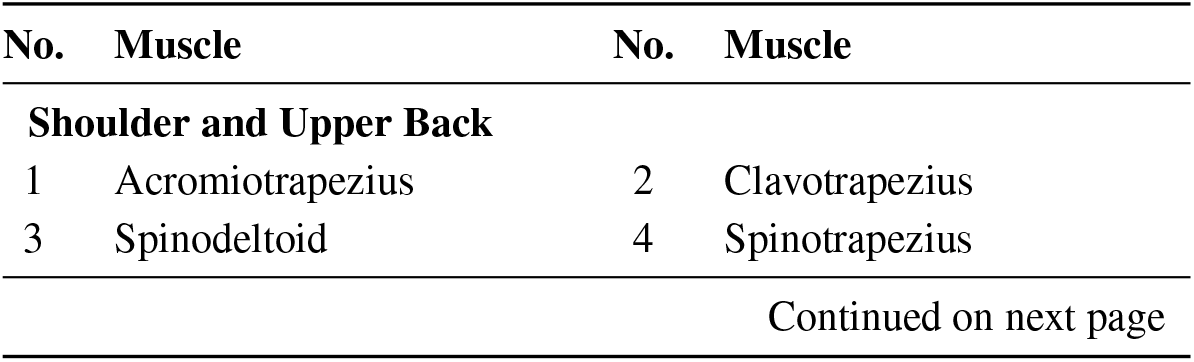
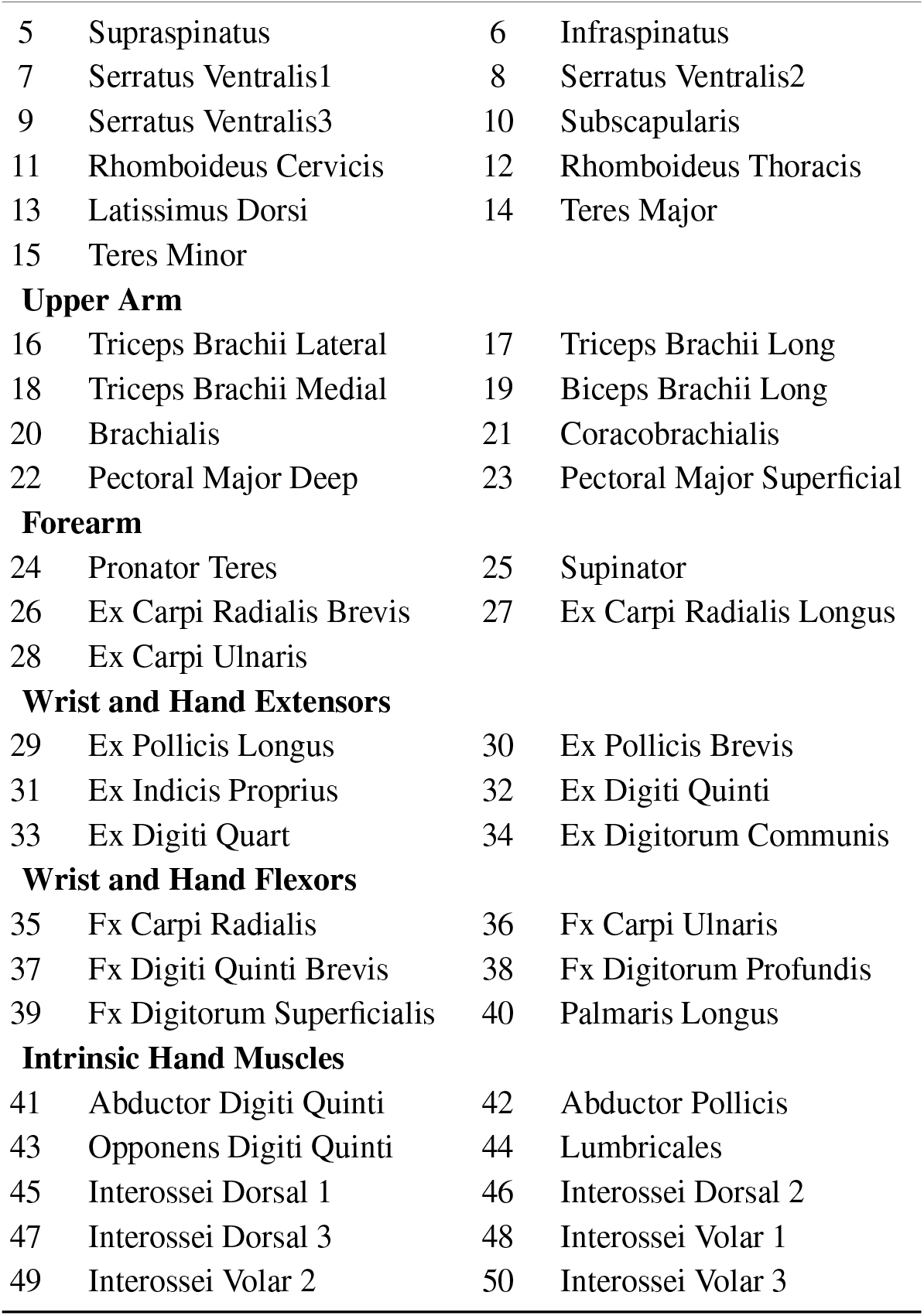
Mouse Arm 50 Muscle List - Grouped by Anatomical Location.

| No. | Muscle | No. | Muscle |
| --- | --- | --- | --- |
| **Shoulder and Upper Back** |  |  |  |
| 1 | Acromiotrapezius | 2 | Clavotrapezius |
| 3 | Spinodeltoid | 4 | Spinotrapezius |
| Continued on next page |  |  |  |
| 5 | Supraspinatus | 6 | Infraspinatus |
| 7 | Serratus Ventralis1 | 8 | Serratus Ventralis2 |
| 9 | Serratus Ventralis3 | 10 | Subscapularis |
| 11 | Rhomboideus Cervicis | 12 | Rhomboideus Thoracis |
| 13 | Latissimus Dorsi | 14 | Teres Major |
| 15 | Teres Minor |  |  |
| **Upper Arm** |  |  |  |
| 16 | Triceps Brachii Lateral | 17 | Triceps Brachii Long |
| 18 | Triceps Brachii Medial | 19 | Biceps Brachii Long |
| 20 | Brachialis | 21 | Coracobrachialis |
| 22 | Pectoral Major Deep | 23 | Pectoral Major Superficial |
| **Forearm** |  |  |  |
| 24 | Pronator Teres | 25 | Supinator |
| 26 | Ex Carpi Radialis Brevis | 27 | Ex Carpi Radialis Longus |
| 28 | Ex Carpi Ulnaris |  |  |
| **Wrist and Hand Extensors** |  |  |  |
| 29 | Ex Pollicis Longus | 30 | Ex Pollicis Brevis |
| 31 | Ex Indicis Proprius | 32 | Ex Digiti Quinti |
| 33 | Ex Digiti Quart | 34 | Ex Digitorum Communis |
| **Wrist and Hand Flexors** |  |  |  |
| 35 | Fx Carpi Radialis | 36 | Fx Carpi Ulnaris |
| 37 | Fx Digiti Quinti Brevis | 38 | Fx Digitorum Profundis |
| 39 | Fx Digitorum Superficialis | 40 | Palmaris Longus |
| **Intrinsic Hand Muscles** |  |  |  |
| 41 | Abductor Digiti Quinti | 42 | Abductor Pollicis |
| 43 | Opponens Digiti Quinti | 44 | Lumbricales |
| 45 | Interossei Dorsal 1 | 46 | Interossei Dorsal 2 |
| 47 | Interossei Dorsal 3 | 48 | Interossei Volar 1 |
| 49 | Interossei Volar 2 | 50 | Interossei Volar 3 |

The fIa and fII muscle spindles and fIb Golgi tendon organ functions from Kibleur et al. (107), Prochazka and Gorassini (110, 111) and Moraud et al. (112) were used to generate the proprioceptive firing rate feedback. The mean parameter values were chosen to match those reported in Kibleur et al. (107), and different proprioceptors (i.e., we simplify to one for each type per muscle) were created by setting parameters using a Gaussian distribution with a standard deviation of 0.1 times the mean. Specifically, we implemented these models as the following:

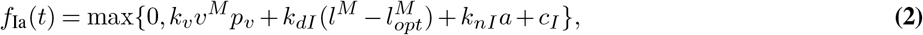

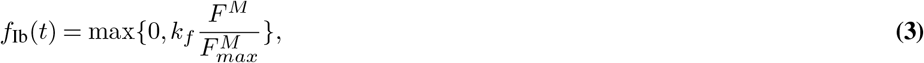

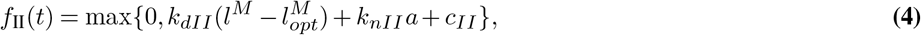

where *l*^*M*^ is the fiber length, 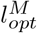 is the optimal fiber length, *v*^*M*^ is the muscle fiber contraction velocity, *a* is the normalized muscle activity, *F*^*M*^ is the force exerted by the muscle fiber, 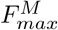 is the maximal isometric force, and *c*_*\**_, *k*_*\**_, and *p*_*\**_ are numerical coefficients determined in Prochazka (113). The *l*^*M*^, *v*^*M*^, *F*^*M*^, and *a* parameters are read from MuJoCo at each time step. The 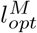 parameter is calculated on model instantiation, and is a function of the length ranges of each muscle, as described see below. The 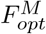 parameter for each muscle is determined through an optimization process, also described below.

#### Building an analytical muscle-based controller

We will initially sketch the idea and then detail the mathematics below. First, we calculate a proportional-derivative (PD) control signal in joint-space, given a target trajectory for each of the joint angles of the arm. We linearize the control of each arm segment by accounting for the inertial and gravitational forces at each joint, which prevents the movement of other arm segments and gravity from affecting control at each joint. Next, to convert this joint torque control signal into a set of muscle activations, we multiply by the negative of the tendon Jacobian (transforming between joint space and tendon space), provided by the MuJoCo simulation engine (74). The resulting muscle activations implement the joint torques specified by the PD controller as best as possible given the current state of the arm, tendon configuration, and muscle parameters.

We define a vector **q** containing the joint angles of the scapula, shoulder, elbow, wrist, and all digits (see Figure 4c), and 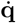 containing their respective derivatives:

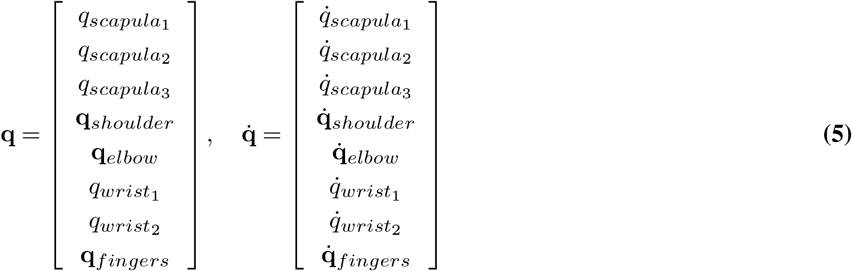

The units and ranges for the scalar elements in this vector are outlined in Table 2. Note that **q**_*shoulder*_ ∈ ℍ and **q**_*elbow*_ ∈ ℍ are quaternions, where ℍ is the 4-dimensional unit hypersphere, and that the derivatives of quaternions are 3-dimensional.

The error signal, 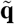, is defined as a function of the target joint angles **q**\* and current joint angles **q**. The error signal is calculated per joint, based on the type of joint. For each of the slide or hinge joints, the error calculation is simply the difference between the target and current positions. For the two ball joints in the arm model (shoulder and elbow), whose position in **q** is represented by 4D quaternions, the error is calculated as

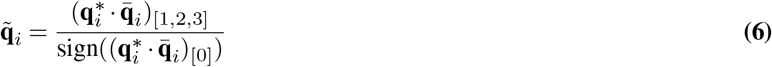

where **q**_*i*_ represents the 4D quaternions (corresponding to the shoulder or elbow joints), · represents quaternion multiplication, and 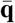 denotes the conjugate of the quaternion **q**. Note that the output of the error calculation for ball joints is a 3-dimensional vector, even though the input are 4-dimensional quaternions. The full error signal is constructed by concatenating the 1D position errors for the three scapula joints, the 3D position error for the shoulder, the 3D position error for the elbow, and then the 1D position errors for the two wrist joints and each finger:

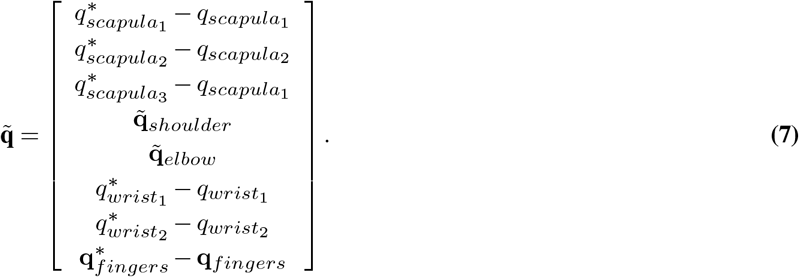

We can now define the control signals driving the model. Given the PD gain terms *k*_*p*_ and *k*_*v*_, the joint space control signal **u**_joint_ is calculated as

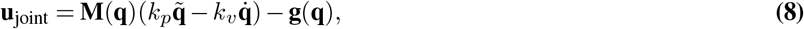

where **M**(**q**) denotes the inertia matrix and **g**(**q**) is the vector of gravitational torques. The joint space control signal **u**_joint_ is then transformed into muscle space using the Jacobian between joints and tendons, **J**(**q**), which captures how the movement of each tendon affects the movement of each joint:

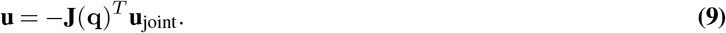

The Jacobian **J**(**q**) is computed analytically based on the configuration of joints and the muscle state and attachment points in the MuJoCo model. The last step of our control signal generation process is to check controller output for overflow values, and set any that are found to zero to prevent crashing the simulation.

Note that instead of muscle-based control, it is also possible to leverage **u**_joint_ as a control signal for direct joint-based control, using the same logic described above. It is also possible to generate a muscle-level control signal based on controlling a specific point on the arm, e.g., the end-effector. To do this requires defining our control signal in end-effector space, and then transforming from end-effector space into joint space using the Jacobian relating joint and end-effector space. From here, the process is the same as above, where compensation for inertia, gravity, and velocity is added, and the signal is transformed to muscle activations. Formally, the full control signal for muscle-based end-effector control is specified:

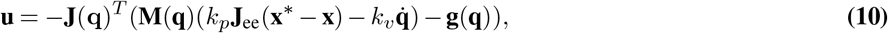

where **J**_ee_ is the end-effector Jacobian, and **x**\* and **x** are the target state and current state of the end-effector. Note that only the calculation of the positional control term changes in this formulation, and compensation for gravity and velocities remains the same. The interface with MuJoCo and the control algorithms are implemented using the ‘abr_control’ package (https://github.com/abr/abr_control).

#### Constraining kinematics with experimental behavior

There are three steps to our process of generating joint angle trajectories for the model mouse arm to follow that recapitulate the mouse movements from experimental sessions. The first step is using DeepLabCut (81, 82) to retrieve 3D estimates of keypoints on the mouse arm from experimental videos (see above), and identify corresponding sites on the mouse arm model in MuJoCo. The chosen keypoints were the shoulder, elbow, three points on the wrist, back of the hand, and each of the digits of the mouse.

Second, the frame-of-reference for the DeepLabCut keypoints needed to be aligned with the MuJoCo frame-of-reference. To do this requires static reference points that are consistent between the experimental videos and the MuJoCo model, that can be used to determine the linear transformations required to move between these frames of reference. We chose to use points on the apparatus the mice stood on and were head-mounted to during the experiment as reference points for alignment. This required implementing a replica of the experimental platform in MuJoCo, which was done using our 3D design CAD files of our apparatus. After identifying the 3D positions of the chosen keypoints from the experimental videos and in the model, the Procrustes linear transformation method (114) was used to transform the 3D pose data into the MuJoCo model reference frame.

Lastly, once the experimental data are aligned with the model reference frame, the inverse kinematics for each experimental trajectory are calculated using a minimization optimization to determine the joint angles that best align the experimental and model 3D keypoints at each point in time. Specifically, we use the ‘scipy.optimize.minimize’ function with the limited-memory Broyden-Fletcher-Goldfarb-Shanno algorithm with an *E* = 0.001 step size. The DeepLabCut keypoints define the configuration of the mouse arm fully enough that the optimization process will converge to the same minima every time it is run. To ensure this, however, we set the initial configuration of the arm during the optimization process to be the resting state angles of the arm, and the cost function includes a term that penalizes distance from this resting configuration. Each subsequent time step is warm-started with the solution from the previous time step to ensure a smooth final trajectory. Formally, our cost function is defined:

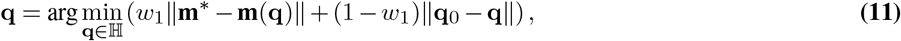

where **m**\* denotes the 3D keypoints from the aligned experimental data, **m**(**q**) is the set of 3D keypoints from the model given the joint angles **q, q**_0_ represents the resting configuration of the arm, and *w*_1_ is a scalar weighting the relative contribution of each term, set to 0.9.

#### Tuning the forelimb model for functionality

There are a total of 511 parameters to tune in the arm model, including stiffness and damping terms for the major joints (digit joints are excluded and use default values), 50 tendon length range upper and lower bounds, and 50 muscle strength terms for each muscle. Additionally, the exact position of the attachment points for each tendon (muscle) can have an effect on the ability of the corresponding muscle to manipulate the arm. The number of attachment points for each tendon can vary between 2 and 5. In total, 115 attachment points are tuned, each with *x, y*, and *z* values. In the controller there are only 2 parameters to tune, the control gains *k*_*p*_ and *k*_*v*_.

The muscle length ranges determine the minimum and maximum length of the model muscles during simulation. To set the muscle length ranges, we took the inverse kinematics across the 26 sessions of experimental recordings (approximately 1,000 minutes of data), set the arm configuration to match the specified inverse kinematic joint angles at each time step, and recording the length of each muscles at every time step. The range of muscle lengths spanned throughout all movements was calculated for each muscle, and minimum and maximum length ranges were set to the minimum and maximum values throughout the movement *±* half the range.

The remaining 411 parameters of the arm model and controller were optimized for functionality in 3 stages: 1) controller gains, 2) tendon attachment points, and 3) joint stiffness, damping, and muscle strengths. The first two stages are optimized independently, and the third stage optimization is dependent on the results of the first two stages.

To tune the model and controller for biologically realistic movements, a set of joint angle trajectories were identified to be reenacted by the arm and controller. Nineteen target trajectories were chosen by analyzing a set of labeled experimental videos and identifying sequences between 1 and 10 seconds where the mouse starts with its hand on the platform, reaches to the joystick, pulls it, releases the joystick, and returns its hand to the platform. After each stage of optimization ran until convergence, we tested the ability of the tuned model to recapitulate full experimental trajectories. Sections in these reenacted full experimental trajectories where the model performed poorly were identified, and one to two second segments from these times were added to the trajectories used during optimization. The optimization process for the joint stiffness and damping and muscle strengths was then performed again with the larger set of target trajectories. We performed four such iterations, using a total of 44 trajectories between all the iterations to optimize the joint stiffness and damping and muscle strengths in the end. Our train error across all markers (excluding the lower-wrist marker, which had a high 3D error due to occlusions) was 0.256 cm *±* 0.183, and test error was 0.201 cm *±* 0.151. For reference, the hand marker train error was 0.190 cm *±* 0.176, and the test error was 0.130 cm *±* 0.119.

To note, running all 44 video clips (2.70136 *±* 0.3764 sec each) it takes roughly 45 minutes (on a workstation, Ubuntu 22.04.4 64-bit with Ryzen 9 5950x 16-core processor x 32 and 64GB memory) to evaluate a hyperparameter set. To speed up evaluation, we generated random parameter sets from a uniform distribution across the search space and used the EPFL RCP cluster to warm-start our hyperparameter sweep - which uses the Tree-Structured Parzens Estimator. In our final iteration of searching, we evaluated 12,628 parameter sets to identify the final parameters.

In the first stage, we tune the PD controller gains using joint torque actuators to control the arm instead of muscle-based actuators. We do this because joint torque control is more precise than muscle-based control, and represents the best possible performance achievable by the muscle-based controller. The cost function (see below) used during this stage is based on the difference between the experimental and simulated joint angles and 3D keypoints, and the simulate joint velocities. We include the 3D keypoints on the arm in the cost function to ensure that deviations at joints earlier in the kinematic tree (e.g., the shoulder or elbow) are weighted with a heavier penalty. The result of a discrepancy at the shoulder has a strong downstream effect, and can result in a trajectory that does not overlap with the experimental trajectory at all, even if all downstream joints follow their target trajectory perfectly. Additionally, we include the weighted sum of velocity at each joint throughout the movement in the cost term, to prioritize parameter sets that more “smoothly” while following the experimental trajectory. Formally, the cost function is:

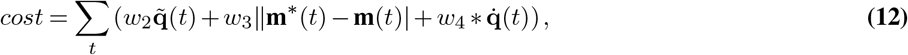

where **q**\* and **q** are the target and actual joint angle trajectories, **m**\* and **m** are the target and actual 3D key-point positions on the arm, **q**? is the joint angles velocities, and *w*_2_, *w*_3_, *w*_4_ are gain terms. The gain terms were set to normalize the angle and 3D key-point costs relative to each other and the velocity term to 1/5the based on values collected during a joint torque controlled movement: *w*_2_ = 0.01212, *w*_3_ = 0.0144, *w*_4_ = 2.94*e* − 6.

In the second stage, the *x, y*, and *z* locations of the 115 attachment points are tuned. The initial placements were made by matching the positions identified by the CT scan analysis as precisely as possible. To account for noise and small errors in the joint modeling and attachment point placement process, a hyperparameter search was conducted across each tendon’s attachment points inside a 3 cubic millimeter space around the initial locations. This fine-tuning is an important step, because the effect each tendon is able to exert over the arm is tightly coupled with their exact attachment point locations relative to positions of joints they cross over.

To evaluate each set of attachment points we moved the arm through the test set of joint angle trajectories, calculated the manipulability term (115) for the tendon Jacobian at each point in time, and summed the manipulability across all time steps and movements. The manipulability term is a scalar measure of the efficacy of actuators on the target task, calculated as the square root of the Gram determinant of the Jacobian. Formally,

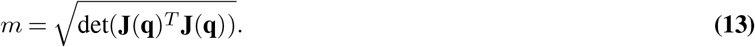

Calculating the manipulability term of the tendon Jacobian in our hyperparameter search provides a metric to evaluate how small variations in the position of a tendon affect the amount of force a muscle is able to exert on the joints it crosses. The units of this metric are referred to as ‘manipulability units’, and we denote them as ‘m’ when reporting numbers in Suppl. Figure S2e.

Lastly, in the third stage of parameter tuning of the arm, the joint damping and stiffness and the muscle strengths were tuned. These results are dependent on the controller gains and attachment points from the first two stages of optimization. Technically, there are stiffness and damping terms for each of the tendons, in addition to the joint damping and stiffness and the muscle strengths terms. In our initial attempts at optimizing the arm parameters, we performed hyperparameter sweeps across 166 joint and muscle parameters. This proved to be too large a problem for an effective search given the viable range for each of the parameters and the available software and computational resources. To overcome this obstacle, the parameter space was heuristically reduced by optimizing for stiffness and damping only at the joints (and setting the stiffness and damping of each tendon to 0), and grouping muscles with similar positioning and effect. This reduction and grouping lead to a much more tractable search problem over 46 parameters (i.e., 3.61x fewer parameters). The muscle groups used during our hyperparameter sweeps are listed in Table 4.

**Table 4.** Muscle groupings. used during optimization of muscle strength parameters.

| Muscle group | Muscles |
| --- | --- |
| pectoral | pectoral major deep, pectoral major superficial |
| serratus ventralis | serratus ventralis 1, serratus ventralis 2, serratus ventralis 3 |
| rhomboideus | rhomboideus cervicis rhomboideus thoracis |
| trapezius | acromiotrapezius, spinatrapezius, clavotrapezius |
| spinatus | supraspinatus, infraspinatus |
| pollicis | ex pollicis brevis, ex pollicis longus |
| interossei dorsal | interossei dorsal 1, interossei dorsal 2, interossei dorsal 3 |
| interossei volar | interossei volar 1, interossei volar 2, interossei volar 3 |

We used the same cost function defined in Eq. (12) for this stage of optimization. The search over these parameters was conducted iteratively, by analyzing the hyperparameter sets from a given search that achieved the best results and adjusting the parameter ranges when there was clustering at a limit.

For all of the parameter sweeps, the Neural Network Intelligence 2.5 (NNI; 116) software was used to conduct the search, which provides access to optimized hyperparameter-space search methods. The controller gain terms used in the final model were *k*_*p*_ = 10000 and *k*_*v*_ = 100. The final model’s joint damping, stiffness, and muscle strength terms are listed in Table 5.

**Table 5.** Joint and Muscle Parameters List. Scale is in relation to the normalized muscle activation profile in MuJoCo.

| Name | Stiffness range (N/m) | Damping range | Length range (cm) | Scale range (N) |
| --- | --- | --- | --- | --- |
| joint scapula1 | 200 - 2330 | 4430 - 4710 | 0 | 0 |
| joint scapula2 | 200 - 2330 | 4430 - 4710 | 0 | 0 |
| joint scapula2 | 200 - 2330 | 4430 - 4710 | 0 | 0 |
| joint shoulder | 3160 - 4570 | 940 - 1280 | 0 | 0 |
| joint elbow | 3350 - 7470 | 1810 - 2320 | 0 | 0 |
| joint wrist1 | 3920 - 8740 | 1440 - 1580 | 0 | 0 |
| joint wrist2 | 3920 - 8740 | 1440 - 1580 | 0 | 0 |
| latissimus dorsi | 0 | 0 | 0.880 - 3.309 | 13800 - 19600 |
| teres major | 0 | 0 | 0.501 - 1.182 | 1600 - 5600 |
| teres minor | 0 | 0 | 0.640 - 1.238 | 500 - 1800 |
| infraspinatus | 0 | 0 | 0.995 - 1.648 | 1500 - 3200 |
| corachobrachialis | 0 | 0 | 0.578 - 0.836 | 1500 - 2000 |
| brachialis | 0 | 0 | 0.846 - 1.398 | 41000 - 47300 |
| triceps brachii long | 0 | 0 | 0.753 - 1.526 | 500 - 2000 |
| triceps brachii lateral | 0 | 0 | 0.897 - 1.213 | 6800 - 27400 |
| triceps brachii medial | 0 | 0 | 0.633 - 0.839 | 2600 - 12800 |
| biceps brachii long | 0 | 0 | 0.784 - 1.419 | 7800 - 11400 |
| spinodeltoid | 0 | 0 | 0.388 - 1.217 | 4100 - 8900 |
| supinator | 0 | 0 | 0.602 - 0.876 | 20300 - 32600 |
| abductor digiti quinti | 0 | 0 | 0.247 - 0.376 | 500 - 2000 |
| ex indicis proprio | 0 | 0 | 1.931 - 2.100 | 3000 - 4100 |
| fx digiti quinti brevis | 0 | 0 | 0.254 - 0.374 | 22800 - 22900 |
| opponens digiti quinti | 0 | 0 | 0.270 - 0.346 | 20600 - 28200 |
| lumbricales | 0 | 0 | 0.622 - 0.881 | 23700 - 32700 |
| ex carpi ulnaris | 0 | 0 | 1.134 - 1.397 | 400 - 3500 |
| fx carpi ulnaris | 0 | 0 | 1.035 - 1.514 | 36100 - 57200 |
| ex digiti quarti | 0 | 0 | 1.809 - 1.974 | 100 - 2000 |

| Name | Stiffness range | Damping range | Length range | Scale range |
| --- | --- | --- | --- | --- |
| palmaris longus | 0 | 0 | 0.929 - 1.497 | 41500 - 69100 |
| ex digiti quinti | 0 | 0 | 1.300 - 1.570 | 900 - 2000 |
| fx digitorum superficialis | 0 | 0 | 1.878 - 2.463 | 5000 - 23100 |
| ex digitorum communis | 0 | 0 | 3.733 - 4.032 | 18500 - 30600 |
| fx digitorum profundis | 0 | 0 | 2.107 - 2.515 | 4400 - 5800 |
| ex carpi radialis brevis | 0 | 0 | 1.302 - 1.513 | 200 - 1400 |
| fx carpi radialis | 0 | 0 | 0.997 - 1.582 | 200 - 200 |
| ex carpi radialis longus | 0 | 0 | 1.277 - 1.488 | 400 - 800 |
| pronator teres | 0 | 0 | 0.397 - 0.944 | 19700 - 24600 |
| pectoral major deep | 0 | 0 | 0.298 - 1.705 | 19600 - 20700 |
| pectoral major superficial | 0 | 0 | 0.119 - 1.705 | 19600 - 20700 |
| serratus ventralis1 | 0 | 0 | -0.519 - 2.248 | 600 - 2000 |
| serratus ventralis2 | 0 | 0 | -0.340 - 2.135 | 600 - 2000 |
| serratus ventralis3 | 0 | 0 | -0.161 - 2.082 | 600 - 2000 |
| rhomboides cervicis | 0 | 0 | -0.136 - 2.471 | 700 - 2000 |
| rhomboides thoracis | 0 | 0 | 0.009 - 2.503 | 700 - 2000 |
| acromiotrapezius | 0 | 0 | 0.557 - 1.581 | 6000 - 8900 |
| spiniotrapezius | 0 | 0 | 0.212 - 3.421 | 6000 - 8900 |
| clavotrapezius | 0 | 0 | 0.455 - 1.926 | 6000 - 8900 |
| supraspinatus | 0 | 0 | 1.006 - 1.327 | 1500 - 3200 |
| abductor pollicis | 0 | 0 | 0.109 - 0.290 | 11300 - 21100 |
| adductor pollicis | 0 | 0 | 0.069 - 0.109 | 11300 - 21100 |
| fx pollicis brevis | 0 | 0 | 0.089 - 0.129 | 10300 - 21100 |
| ex pollicis brevis | 0 | 0 | 0.632 - 0.789 | 2900 - 4100 |
| ex pollicis longus | 0 | 0 | 0.746 - 0.903 | 2900 - 4100 |
| interossei dorsal 1 | 0 | 0 | 0.382 - 0.451 | 3500 - 4100 |
| interossei dorsal 2 | 0 | 0 | 0.452 - 0.481 | 3500 - 4100 |
| interossei dorsal 3 | 0 | 0 | 0.412 - 0.446 | 3500 - 4100 |
| interossei volar 1 | 0 | 0 | 0.226 - 0.423 | 1900 - 5400 |
| interossei volar 2 | 0 | 0 | 0.286 - 0.447 | 1900 - 5400 |
| interossei volar 3 | 0 | 0 | 0.276 - 0.430 | 1900 - 5400 |

#### State Estimator Modeling

The force field operates orthogonal to the pull axis. Motor adaptation can thus be analyzed by modeling the changes of the perpendicular deviation from the pull axis midway through the pull (i.e., during the force field, when applied). We utilized the state estimator model previously deployed in Mathis et al. (1), which was modified from Izawa and Shadmehr (64). We built a discrete dynamical system that models the evolution of the scalar perpendicular deviation of the mouse hand and how sensory prediction error driven signals act upon it. We model the perpendicular displacement of the joystick *x*_*k*_ at a particular position and in trial *k* is the consequence of a motor command *u*_*k*_, the applied perturbation *p*_*k*_, and normally distributed sensorimotor noise 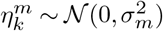, such that:

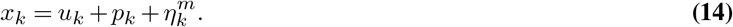

We assume that the mouse internally estimates its actual hand position by 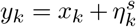 and also modified by sensory noise 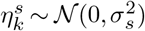. We assume that their internal estimate of the perturbation is given by a discrete dynamical system with retention factor *r* (less or equal to 1; fitted based on data) and memory noise 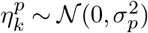:

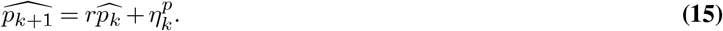

The goal of this modeling is to estimate the internal motor control policy that mice use to adapt. These are then used as proxy variables to measure their effect on neural encoding (both at the single cell and population level). Thus, to model the motor command *u*_*k*_, we assume the mice estimate their hand position 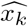 based on the efference copy of their motor signal and their internal estimate of the perturbation 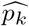 by:

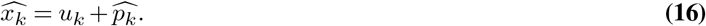

Thus, the optimal behavioral control policy can be derived from the optimal state estimator and the optimal policy (17, 64, 117).

#### Sensory prediction error driven learning

As used in Mathis et al. (1), the state estimator is given by the Kalman filter (18, 64, 118, 119). The generative model – internal model – can be written as a state evolution model:

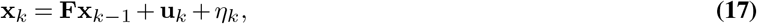

with state transition model 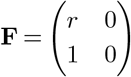, control input model 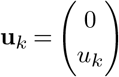, process noise *η*_*k*_ ~ 𝒩 (**0 Q**) with 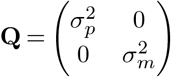 and states 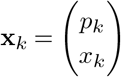. The observer model is given by:

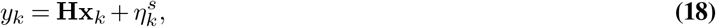

with measurement matrix **H** = (0 1) and sensory noise 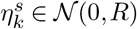 with 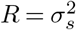.

In this situation the classical results by Kalman imply that the optimal state estimation is given by recursively iterating (118, 119):

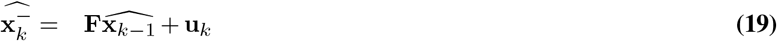

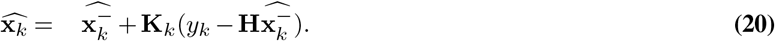

The first equation describes the *a priori* prediction of the hand position and perturbation. The subsequent state estimate is a combination of this *a priori* estimate and the sensory prediction error 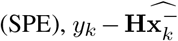, gated by the Kalman gain matrix, **K**_*k*_ (i.e., 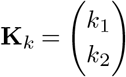). The Kalman gain matrix is updated according to:

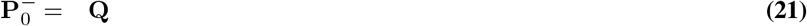

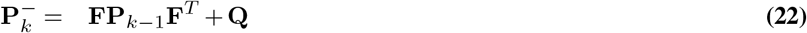

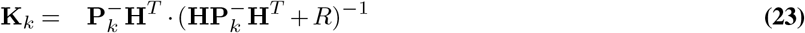

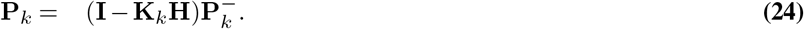

Given this state estimate, the optimal command to compensate for the perturbation is: 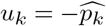. **F** is the model dynamics. *R* is the measurement noise covariance, representing the uncertainty in the measurements. 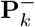 is the predicted (*a priori*) error covariance matrix at time step *k*. It estimates the uncertainty before the measurement update. **F** is the state transition matrix, which models how the state evolves from time step *k* − 1 to *k. I*: The identity matrix.

To fit the model, we supply the perpendicular deviation during the force field per trial, averaged across mice. Then we fit during only the perturbation epoch (trials 76-175) with the differential evolution algorithm (from scipy). We estimated the *p*_*k*_ to be the mean of the first three adaptation trials across sessions.

#### Generalized Linear Modeling (GLM)

One of the primary goals of our study was to establish which features – posture, dynamics (velocity, torque), or muscles – are encoded in individual sensorimotor neurons in the mouse M1 and S1 forelimb areas, as this had not been previously described at this resolution with matched behavioral kinematics. To this end, we used Generalized Linear Models (GLMs) as adapted in prior works (40, 120) to predict the neuronal responses across the sessions.

To examine how much variance of the neural activity could be directly explained by behavioral variables, a GLM with an exponential nonlinearity was used. The deconvolved neural activity was binarized (with a threshold of 0.01). We constructed six design matrices containing different features derived from the different groups (Figure 6a-c, Suppl. Figure S6 and see Table below). Features were individually min-max-scaled to a range (0,1) prior to inputting them to the GLM. Binarized activity was predicted from a single time lag to limit the number of regressors in the design matrix. To note, this is therefore a stringent criteria that will decrease the fraction of neurons considered tuned. The GLM model is then parametrized as

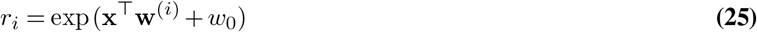

where *r*_*i*_ is the rate of neuron *i* predicted from regressors **x** (including a bias term *w*_0_), and **w**^(*i*)^ are the regression weights.

To compensate for behavioral variability within sessions, all models were trained and tested on five splits of the data (each with 20% test set in three equally spaced blocks). Predictions on the test portion of the dataset are then combined, and used to compute McFadden’s Pseudo-R^2^,

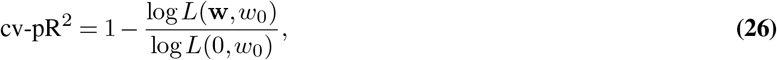

where log *L*(**w**, *w*_0_) is the log-likelihood of the fitted model, and log *L*(0, *w*_0_) is the log-likelihood of the null model (a model with only the intercept). We then used a pseudo-*R*^2^ threshold of 0.025 and only further considered neurons above this threshold. This was determined based on capturing approx. 40-50% of the variability, as used in primate sensorimotor studies (40), and matches prior reports of fraction of tuned neurons in mouse M1 (62, 121).

We fit these GLM models on all six regression groups defined in Table 6. Within each group, we analyze feature importance by further splitting the regressors into subgroups corresponding to body parts, and the type of signal (positions, velocities, joint angles, joint velocities, etc.). We then analyzed the feature importance within each subgroup. We used the regression weights **w**^(*n*)^ ∈ ℝ^*d*^. For a subgroup *G* (here, a subset of indices from 1…*d*) of regressors, we computed the length of the regression weight vector,

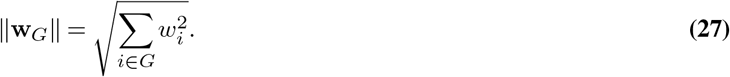

**Table 6.**
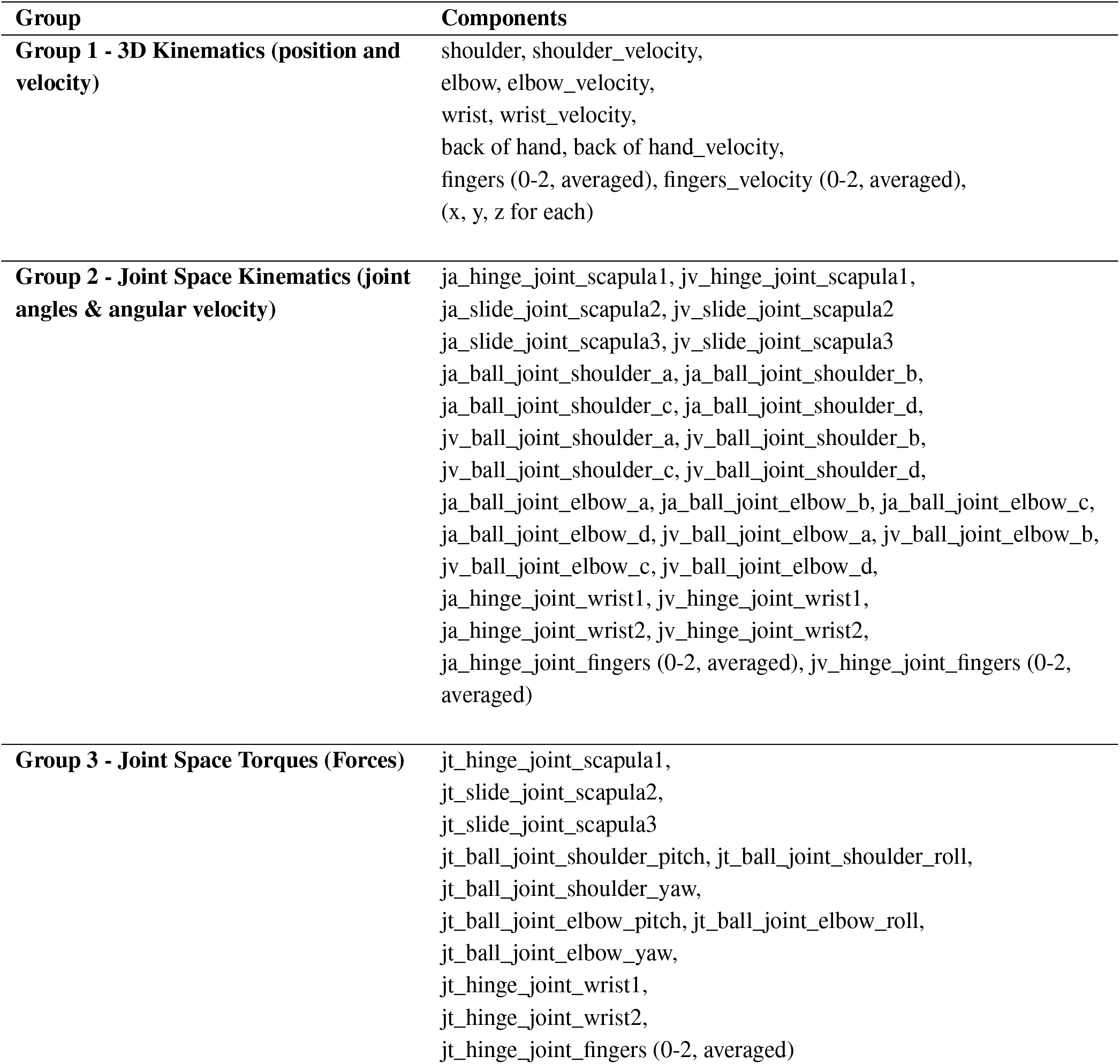
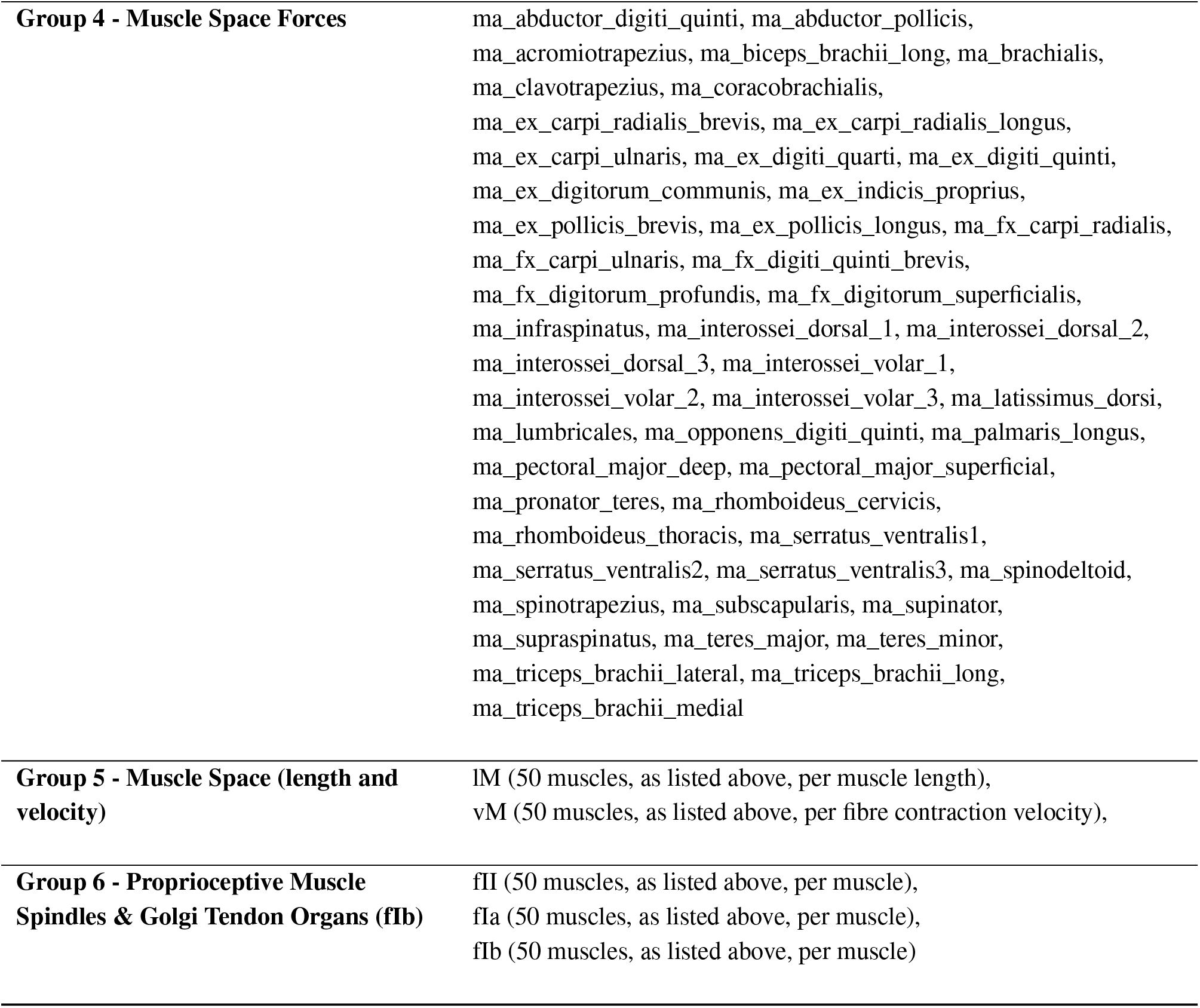
Overview of six different groups of regressors in our GLM analysis.

| Group | Components |
| --- | --- |
| **Group 1 - 3D Kinematics (position and velocity)** | shoulder, shoulder_velocity,<br>elbow, elbow_velocity,<br>wrist, wrist_velocity,<br>back of hand, back of hand_velocity,<br>fingers (0-2, averaged), fingers_velocity (0-2, averaged),<br>(x, y, z for each) |
| **Group 2 - Joint Space Kinematics (joint angles & angular velocity)** | ja_hinge_joint_scapula1, jv_hinge_joint_scapula1,<br>ja_slide_joint_scapula2, jv_slide_joint_scapula2<br>ja_slide_joint_scapula3, jv_slide_joint_scapula3<br>ja_ball_joint_shoulder_a, ja_ball_joint_shoulder_b,<br>ja_ball_joint_shoulder_c, ja_ball_joint_shoulder_d,<br>jv_ball_joint_shoulder_a, jv_ball_joint_shoulder_b,<br>jv_ball_joint_shoulder_c, jv_ball_joint_shoulder_d,<br>ja_ball_joint_elbow_a, ja_ball_joint_elbow_b, ja_ball_joint_elbow_c,<br>ja_ball_joint_elbow_d, jv_ball_joint_elbow_a, jv_ball_joint_elbow_b,<br>jv_ball_joint_elbow_c, jv_ball_joint_elbow_d,<br>ja_hinge_joint_wrist1, jv_hinge_joint_wrist1,<br>ja_hinge_joint_wrist2, jv_hinge_joint_wrist2,<br>ja_hinge_joint_fingers (0-2, averaged), jv_hinge_joint_fingers (0-2, averaged) |
| **Group 3 - Joint Space Torques (Forces)** | jt_hinge_joint_scapula1,<br>jt_slide_joint_scapula2,<br>jt_slide_joint_scapula3<br>jt_ball_joint_shoulder_pitch, jt_ball_joint_shoulder_roll,<br>jt_ball_joint_shoulder_yaw,<br>jt_ball_joint_elbow_pitch, jt_ball_joint_elbow_roll,<br>jt_ball_joint_elbow_yaw,<br>jt_hinge_joint_wrist1,<br>jt_hinge_joint_wrist2,<br>jt_hinge_joint_fingers (0-2, averaged) |
| **Group 4 - Muscle Space Forces** | ma_abductor_digiti_quinti, ma_abductor_pollicis,<br>ma_acromiotrapezius, ma_biceps_brachii_long, ma_brachialis,<br>ma_clavotrapezius, ma_coracobrachialis,<br>ma_ex_carpi_radialis_brevis, ma_ex_carpi_radialis_longus,<br>ma_ex_carpi_ulnaris, ma_ex_digiti_quarti, ma_ex_digiti_quinti,<br>ma_ex_digitorum_communis, ma_ex_indicis_proprius,<br>ma_ex_pollicis_brevis, ma_ex_pollicis_longus, ma_fx_carpi_radialis,<br>ma_fx_carpi_ulnaris, ma_fx_digiti_quinti_brevis,<br>ma_fx_digitorum_profundis, ma_fx_digitorum_superficialis,<br>ma_infraspinatus, ma_interossei_dorsal_1, ma_interossei_dorsal_2,<br>ma_interossei_dorsal_3, ma_interossei_volar_1,<br>ma_interossei_volar_2, ma_interossei_volar_3, ma_latissimus_dorsi,<br>ma_lumbricales, ma_opponens_digiti_quinti, ma_palmaris_longus,<br>ma_pectoral_major_deep, ma_pectoral_major_superficial,<br>ma_pronator_teres, ma_rhomboideus_cervicis,<br>ma_rhomboideus_thoracis, ma_serratus_ventralis1,<br>ma_serratus_ventralis2, ma_serratus_ventralis3, ma_spinodeltoid,<br>ma_spinotrapezius, ma_subscapularis, ma_supinator,<br>ma_supraspinatus, ma_teres_major, ma_teres_minor,<br>ma_triceps_brachii_lateral, ma_triceps_brachii_long,<br>ma_triceps_brachii_medial |
| **Group 5 - Muscle Space (length and velocity)** | IM (50 muscles, as listed above, per muscle length),<br>vM (50 muscles, as listed above, per fibre contraction velocity), |
| **Group 6 - Proprioceptive Muscle Spindles & Golgi Tendon Organs (flb)** | fII (50 muscles, as listed above, per muscle),<br>fIa (50 muscles, as listed above, per muscle),<br>fIb (50 muscles, as listed above, per muscle) |

We obtain this quantity for every neuron in the dataset, and denote the corresponding regression weight lengths 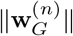 for each neuron *n* in the dataset. For groups with at least two subgroups, we tested the regressor weights on mixed selectivity to both subgroups. The distribution of 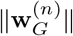 across neurons *n* approximately followed a log-Normal distribution. We first determined the winning subgroup for each neuron by

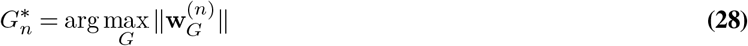

and then computed the mean and standard deviation 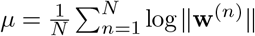 and 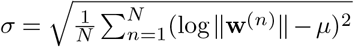 across the regression weights on the five data splits during cross-validation. To consider a neuron to be tuned to the group *G*\*, we required that

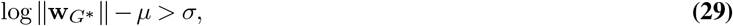

and neurons that cross the significance threshold for multiple groups without fulfilling this criterion are called mixed selective neurons. At the end of this process, we obtained assignment of each neurons to none, one, or multiple subgroups within each of the six groups.

We also assigned neurons to one of the six groups for part of the analysis by a winner-takes-all approach within the neurons exceeding a pseudo-*R*^2^=0.025 on at least one group.

Finally, we ran a variant of GLMs trained on each individual regressor within each group. We performed a sub-selection based on the highest tuned regressors from the multi-regressor analysis by group. By this, we obtained pseudo-*R*^2^ values for every neuron relative to every regressor. For these unique contribution GLMs we additionally considered rewards. We filtered the binarized valve opening time (recorded in hardware TTL) with a Gaussian filter (*σ* = 500) to consider the period when the mice consume the rewards.

#### Neural population analysis with CEBRA

To measure neural representations we used CEBRA, a new self-supervised learning algorithm for obtaining interpretable, **C**onsistent **E**m**B**eddings of high-dimensional **R**ecordings using **A**uxiliary variables to test the relationship of behavioral, learning, and task variables to the neural dynamics. In brief, CEBRA contrasts a reference example **x** to a given positive example (**y**_+_), and multiple negative examples (**y**_*i*_) and minimizes the loss function:

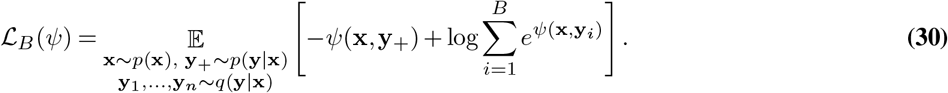

for a positive sample distribution *p*(·|·) and a negative sample distribution *q*(·|·). To measure how well the CEBRA model is fit compared to a null model, we define the goodness of fit (GoF; in bits) as

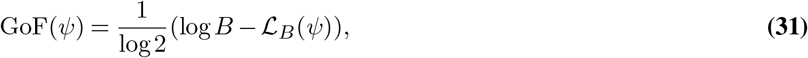

where *B* is the batch size and ℒ the InfoNCE loss of a converged CEBRA model *ψ*. A goodness of fit of 0 bits denotes chance level, higher goodness of fit values are better.

For each adaptation session we fit an individual embedding using CEBRA-Time. Specially, we used the ‘offset10-model-mse’ model, batchsize = 512, learning rate = 5e-5, temperature = 0.01, output_dimension = 3, distance = euclidean, and time_offsets = 10. We set the maximum iterations based on the convergence of the loss function (range: 200-3000).

We trained separate embeddings on 50-trial sections of the data, from baseline (trials 25-75), to early perturbation (trials 76-125), late perturbation (126-175), and washout (trials 176-225) using a continuous auxiliary variable around the trial TTL (red line in Figure 1a), which is also the same position bin as the force-field onset time (also shown as a red line in Figure 2a) to compare the consistency across learning. This continuous variable is computed by assigning values from −100 to 500 samples around trial TTL to the sequence. Neural data from multiple sessions is aligned on the fly based on the trial TTL to learn a single unified embedding (122) across all n=4 adaptation sessions. We used the ‘offset10-model’ model, batchsize = 512, temperature = 0.5, learning rate = 1e-4, time_offset = 10, hidden_dim = 128, output_dim = 10 and trained for 2,500 iterations. We used either all neurons or split the training into M1 and S1 models. To measure changes in the latent space we used the consistency metric described in Schneider et al. (65). To estimate consistency, all embedding points were binned by the trial TTL auxiliary variable, now only considering −1 and 1 sec around the trial time TTL (i.e., force-field onset). We compute the mean embedding per bin, and estimate the consistency metric by fitting a linear regression model and averaging *R*^2^ scores for forward and backward prediction between the embeddings across the four phases.

### Other data analysis methods

#### Identifying pulls and reaches (for muscle correlations)

Identifying stereotyped reaching and pulling movements in the experimental data was performed in two stages, first, determining candidate reach and pull times in the experimental data, and secondly, filtering the candidate reach and pull movements for stereotyped kinematics. To determine candidate reach and pull times, we first isolated the y-axis movements of the mouse hand and joystick, which is the principal axis for reaching and pulling movements. For reaching movements, the mouse right hand DeepLabCut marker was used to identify reach times, and for pulling movements, the joystick marker was used to identify pull times. Next we defined threshold-lines along the y-axis, which, when crossed, suggest a reach from the platform to the joystick or a pull of the joystick has occurred. For reaches, this was done by performing binary classification of the y-axis data as above or below threshold, calculating the derivative to identify points where the threshold was crossed, filtering for movements from the platform to the joystick, and then a secondary filtering removing any partial reaches where the mouse returns to the platform within 300 milliseconds. For reaches, we use −0.75 centimeters along the y-axis as the threshold boundary point to determine candidate moments for reach times. We then require that in the 150 milliseconds before the reach, the mouse and model hand are under the −0.5 centimeters line on the y-axis, and that within 150 milliseconds after the reach time they cross beyond −1.8 centimeters.

The pull times we defined by a TTL pulse that was defined at a positional bin (as shown in Figure 1a b). With the reach and pull times, we analyzed a window from 150 milliseconds before and after the event time. We checked to make sure that for the full 150 milliseconds before the reach or pull time that both the mouse and model hand position was inside the threshold area, and that at some point between the reach or pull time and 150 milliseconds after the mouse and model hand are beyond a secondary threshold line representing a successful pull or reach. When analyzing the pull movements, we also require that the joystick marker is moving with the hand, to ensure that we are isolating actual pulls of the joystick and not including any movements where the hand is simply returning to the platform.

For pulls, we require that all joystick and mouse and model hand positions are beyond −1.85 centimeters in the 150 milliseconds before the pull time, and that within 150 milliseconds after the pull time they cross the −1.5 centimeters line along the y-axis.

#### EMG analysis and cross-correlation calculations

To generate muscle EMG signals, we post-processed the muscle activation signals generated by our model following the steps described in (44). Specifically, we downsampled the signals to 1 kHz, high-pass filtered at 40 Hz, rectified, and convolved with a Gaussian having a 10 ms standard deviation. Using the reach and pull times identified above, we then generated the crosscorrelation matrix using data from 50 milliseconds before the trial time to 100 milliseconds after using the NumPy corrcoef function.

#### Peri-stimulus time histograms

To compute PSTH plots we used the extracted estimated spiking rates signal then extract a window before and after a defined TTL that is defined by a positional bin during the pull. As shown in Figure 1a b, this position is approximately peak velocity of pulls, and as shown in Figure 2a,b the time the force-field comes on during perturbation trials only. To compare across epochs, as in Figure 2 we normalized the activity by 10 timesteps within the pre-TTL period. For computing the max activity in Figure 2g we computed the maximum firing rate across neurons within the time period after the force field onset or the equivalent bin in non-force field trials (i.e., baseline and washout).

#### PCA and hierarchical clustering

For functional clustering across trials in Figure 7a we constructed a vector of neuron x trials across the epochs. Namely, we take the data as shown in Suppl. Figure S1c which is *±*1 sec before the trial TTL (red line as shown in Figure 1a), which is the point of force-field onset during the adaptation task. We normalized activity within the epoch to account for any systematic changes across the session. Then we take the n=1697 M1 an S1 neurons and ran PCA with *n* = 20 components using scikit-learn. Then, we used Agglomerative Clustering algorithm to perform hierarchical clustering. The number of clusters was optimized for non-overlapping functional groupings, and set to *n* = 5.

To assess the statistical significance of the clustering results, we performed a permutation test using 10,000 permutations. For each permutation, the trial activity data was randomly shuffled and PCA (with n=10 components) was applied to the reshuffled data. Next, hierarchical agglomerative clustering with Ward linkage was applied to the PCA-transformed data, and the cluster labels were obtained. The Adjusted Rand Index (ARI) was calculated using the scikit-learn API (adjusted_rand_score) to compare the clustering labels of the observed data with those from each permutation. The p-value was computed as the fraction of permutations where the ARI score was greater than or equal to the ARI of the observed clustering result. A low p-value indicates that the observed clustering is significantly different from random clusterings.

#### Statistical testing and software

All statistical tests were computed in Python 3.8 (or above) with scipy or statsmodels and tests were two-sided. Alpha was preset to 0.05. Non-parametric tests were used when data were not normally distributed. For the permutation test we bootstrapped with n=1000 permutations. We used sklearn to compute the change from baseline per epoch of the Area under the Receiver Operating Characteristic curve (AUC). We used Python, MuJoCo (74), NumPy (123), SciPy (124), Pytorch (100), TensorFlow (125), Scikit-learn (126), DataJoint (101), DeepLabCut (81, 82), and CEBRA (65).

## Supplemental Materials

VIDEO 1: Video showing the 3D lifted DeepLabCut outputs and input video.

VIDEO 2: Video demonstrating MusBioMaus reenacting the main mouse behavior: reach, pull, and return hand to base.

VIDEO 3: Overview of data from an adaptation session with CEBRA embeddings from S1 and M1. Top left: we show a 2D video with DeepLabCut overlay. Bottom left: the joystick movements (top view). Middle: co-recorded M1 and S1 neurons for an example session, and below are joystick and extracted hand coordinates. Right top: S1 embedding, Right bottom: M1 embedding. Epoch as shown in video title.

**Figure S1.**
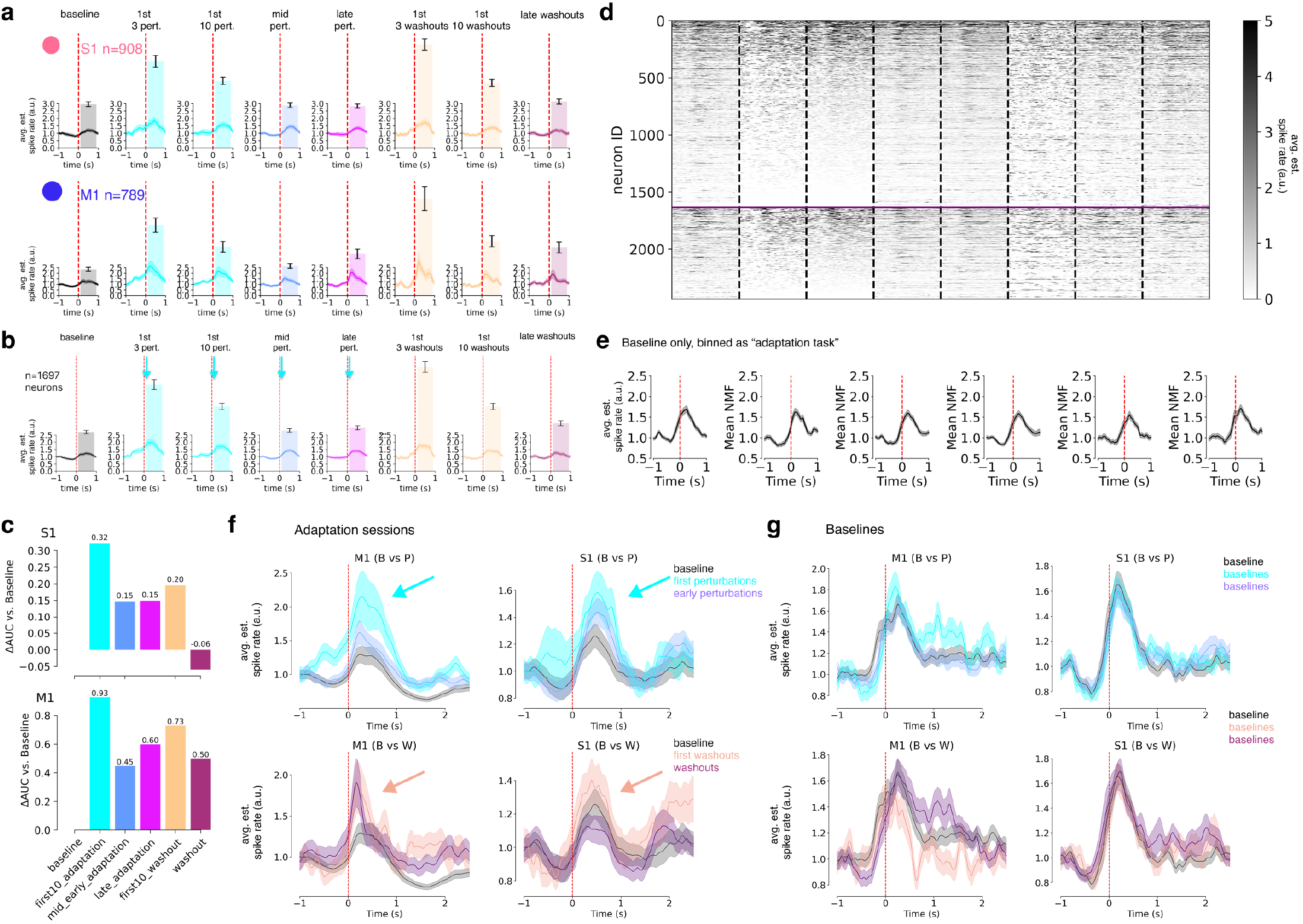
Neural properties during adaptation. **a**, PSTH’s computed based on force-field onset time (delineated by the red line): −1 sec to 1 sec. Note, the force-field was only applied in perturbation trials. M1 or S1 neurons are plotted separately and trials included are denoted in the plot title. The background bar plot is the mean maximum estimated spiking rates computed after the force field onset, which shows the same data trend across blocks. **b**, PSTH plots for all neurons shown in **a** (not split by S1 and M1). **c**, AUC analysis of M1 or S1 neurons. Plot shows a change in AUC per epoch compared to baseline. **d**, Heatmap of all neurons recorded during baseline only sessions (n=11 sessions with >250 trials for comparison) as a control. **e**, PSTH plots of baseline only sessions (n=11 with >250 trials), yet binned as in panel **b & c** as a control. There was no significant change from baseline in any binned block (t-test p-values, baseline vs. other blocks left to right: 0.15, 0.68, 0.33, 0.14, 0.33). **f**, PSTHs showing the difference in activity in M1 or S1 neurons during baseline, early perturbations, mid-perturbations, early washout and washout (as colored in the legend). **g**, Same as in **e** yet for only baseline sessions as a control.

**Figure S2.**
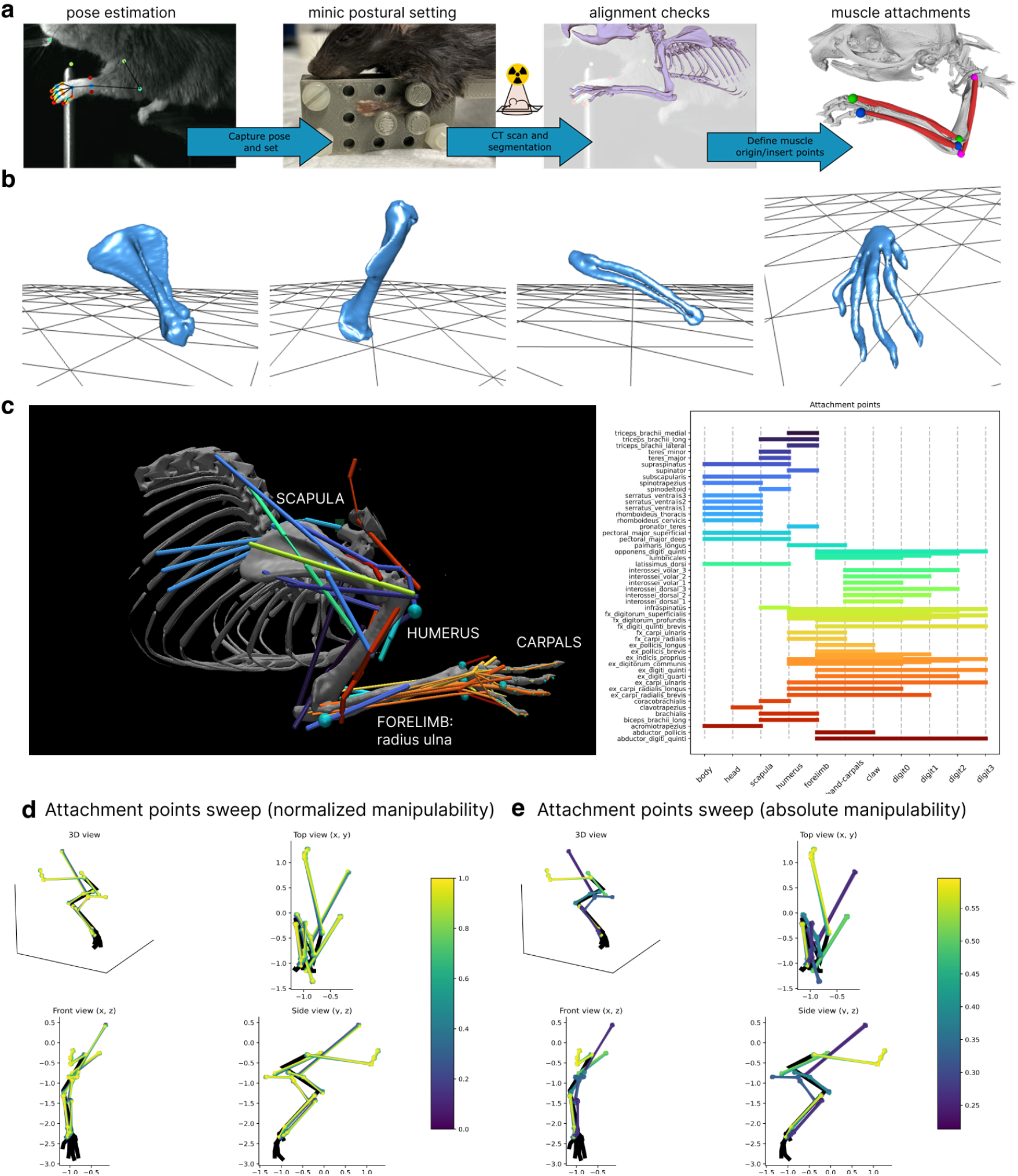
Development and optimization of the adult mouse forelimb model. **a**, overview of posing mice in experimentally realistic postures for CT scanning, with post-hoc alignment and muscle attachment configuration. **b**, Example segmented forelimb showing the scapula, humerus, forelimb (ulna and radius) and hand (here as a “single” image). **c**, Overview of the major groupings (left) attachment points of the 50 muscles modeled (right). **d**, Sample attachment point configurations tested during in the hyperparameter sweep for several muscles. Each muscle’s color represents the resulting manipulability score calculated using Eq. (13), normalized per muscle. Within each set of configurations, small variations can lead to large changes in manipulability. **e**, Sample attachment point configurations tested during in the hyperparameter sweep for several muscles. Each muscle’s color represents the resulting absolute manipulability score. This plot highlights the relative contributions of each of the muscles to the arm’s overall manipulability, calculated using Eq. (13). More proximal muscles, such as the latissimus dorsi or serratus ventralis have relatively low manipulability scores (see Methods for a discussion on the metric) for all configurations, 0.27 and 0.60, respectively. The extensor carpi ulnaris, which spans the elbow and the wrist joints, has several attachment point configurations with relatively high manipulability scores, 0.58. The high manipulability scores for some attachment point sets for extensor carpi ulnaris indicates that its exact placement is the most important of this set of muscles to be able to manipulate the arm as effectively as possibly. The initial attachment point set (placed by hand) achieved a manipulability score of 0.09. Applying the best attachment point sets for each muscle found with a hyperparameter search inside a 0.03^3^ mm box around each initial location achieved a manipulability score of 99.09.

**Figure S3.**
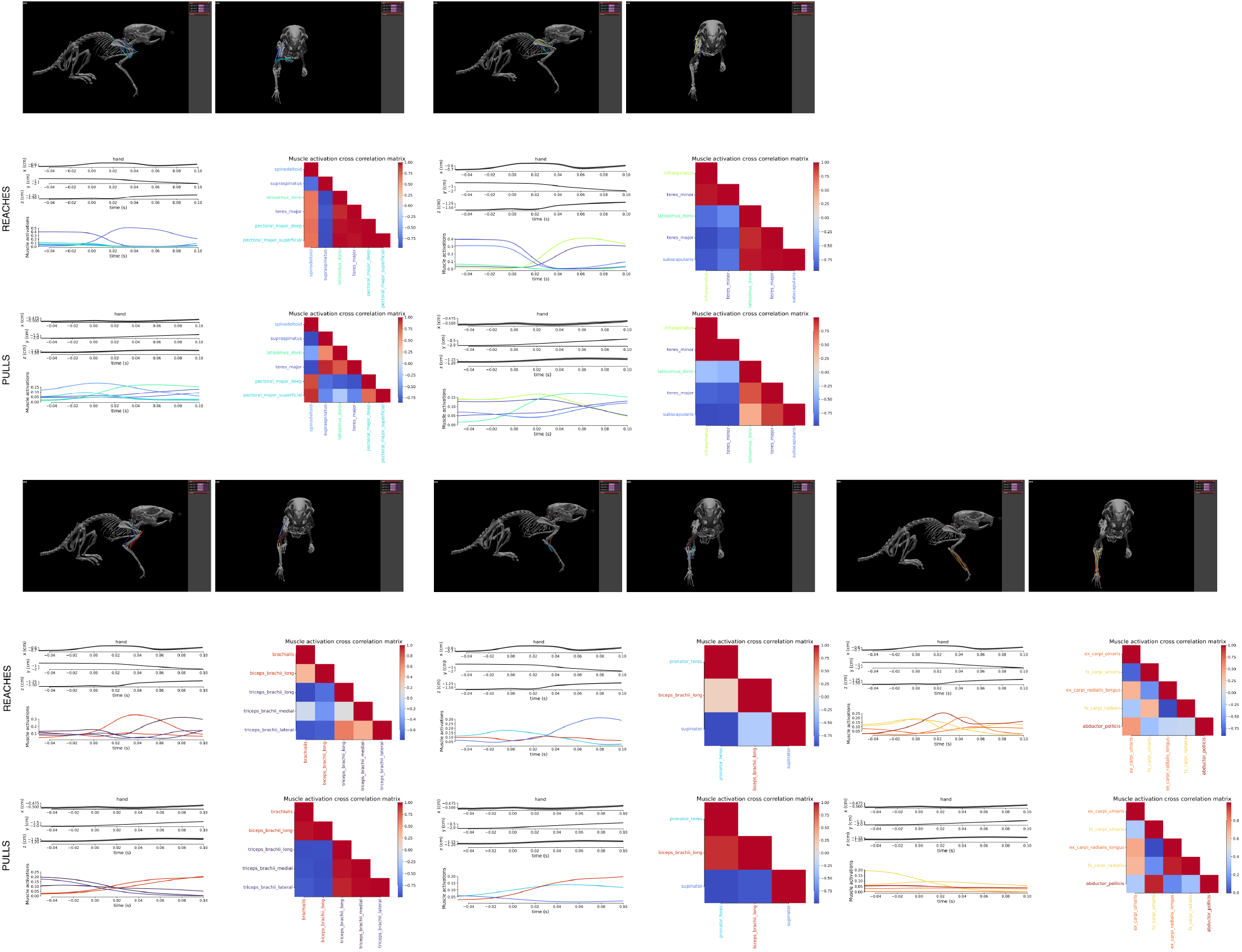
MusBioMaus muscle outputs. **a**, We measured the correlation of muscles during pull (n=435) and reach (n=342) epochs and then computed the average correlation over the time-window shown. Within each panel the hand x, y, and z mean position and 95% confidence intervals are shown on the top left, bottom left is the mean muscle activations (see Methods), and right is the correlation matrix.

**Figure S4.**
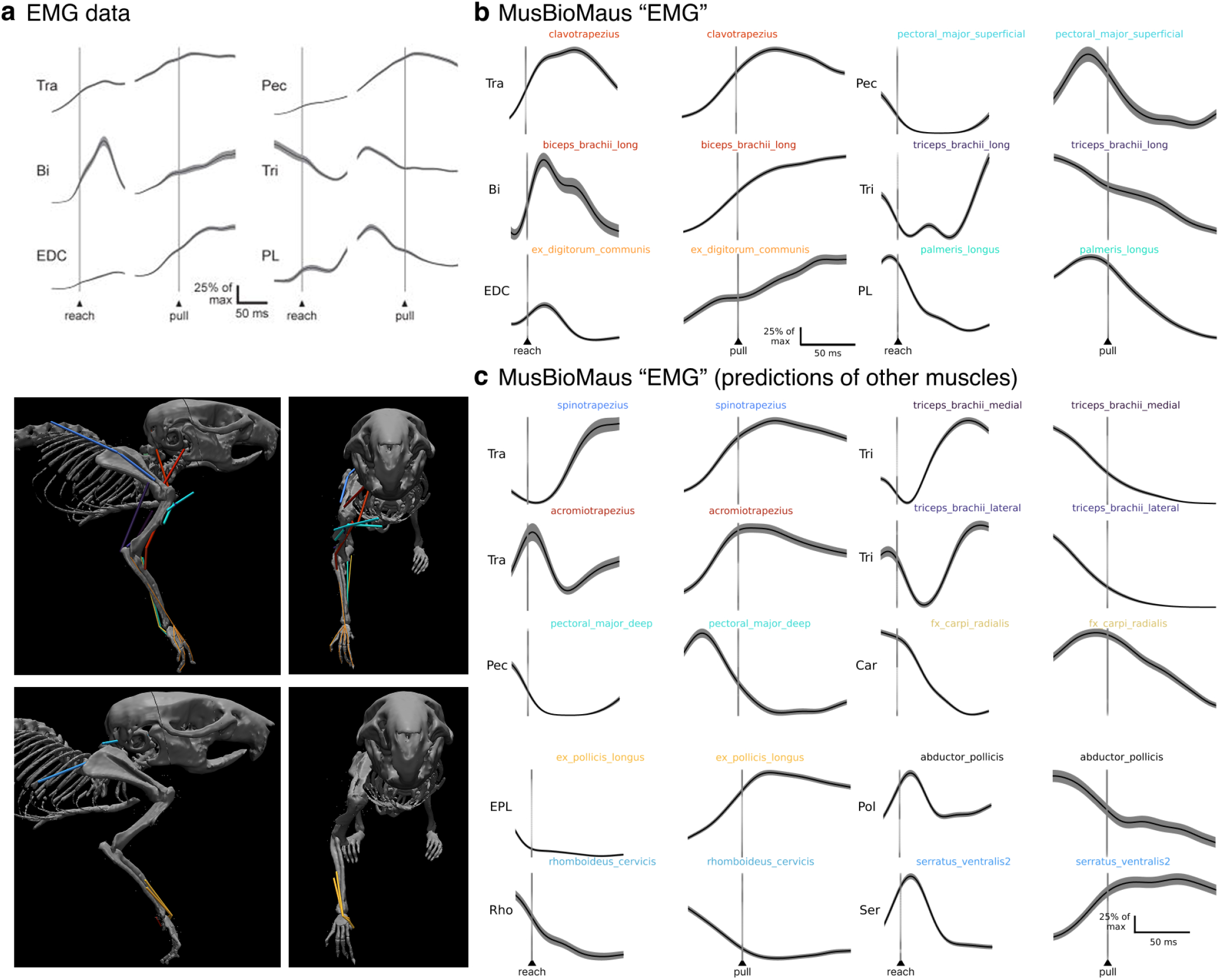
EMG comparisons. **a**, Adapted from Miri et al. (44), where head-fixed mice are reaching and then pulling on a lever. They recorded directly from six muscles: Tra = Trapezius, Tri = triceps, Bi = Biceps, EDC = extensor digitorum communis, and PL = palmaris longus. Note, the kinematics are not identical to our synthetic mouse (as the behavioral tasks are different), but we show in **b** that our general extracted EMG-like signals during “reaching” and “pulling” are reasonably conserved in the 6 muscles Miri et al. (44) recorded from. We calculated the mean and SEM of these EMG-like signals using the muscle activations during pull (n=435) and reach (n=342) epochs. Note, in **c** we highlight we can make other predictions (up to 50 in total), and some are also shown in Figure 5d. **c** shows an additional subset of muscles related to the upper arm, i.e., additional muscles of the Triceps (Tri), Pectoral muscles (Pec) and Trapezius (Tra)); the forearm, i.e., ex. pollicis longus (EPL), abductor pollicis (Pol); and shoulder, i.e., rhomboideus cervicis (Rho) and serratus ventralis(2, Ser).

**Figure S5.**
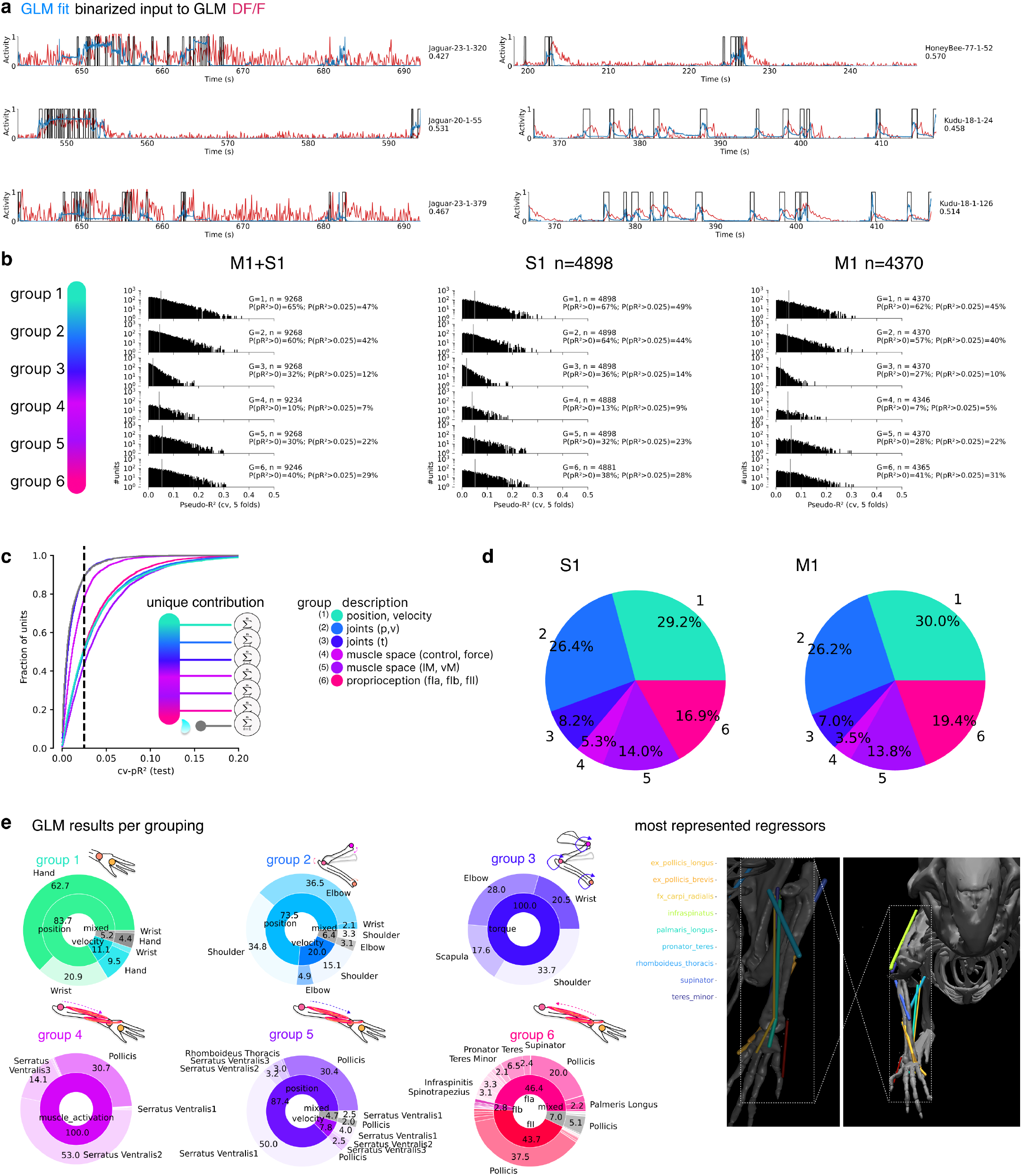
GLM models. **a**, Example GLM model fits of individual neurons on the held-out test dataset. These are selected from the best-fit neurons. pseudo-*R*^2^ is reported on the far right, and each signal is min-max normalized. **b**, Full GLM run results, showing the distribution of all neurons (n) per group (L). The percentage of neurons above a given pseudo-*R*^2^ threshold are inset. Middle and right plots show results for M1 and S1 independently. **c**, GLM results where each neuron is regressed independently against all regressors (see Table 6) then post-hoc placed into a given group. This controls for group-effects. **d**, Pie chart of the tuned neurons (only), for the percentage of neurons with a *cv* − *pR*^2^ above 0.025 for each group (thus, a neuron can be counted in more than one category). **e**, Wheel-plots show within each group the tuning of neurons. In the inner ring is the group factors and the outer ring is the specific features that are significantly tuned. Each ring includes only significantly tuned neurons. Far right: images of the muscles most represented in sensorimotor cortex of our mice during this task.

**Figure S6.**
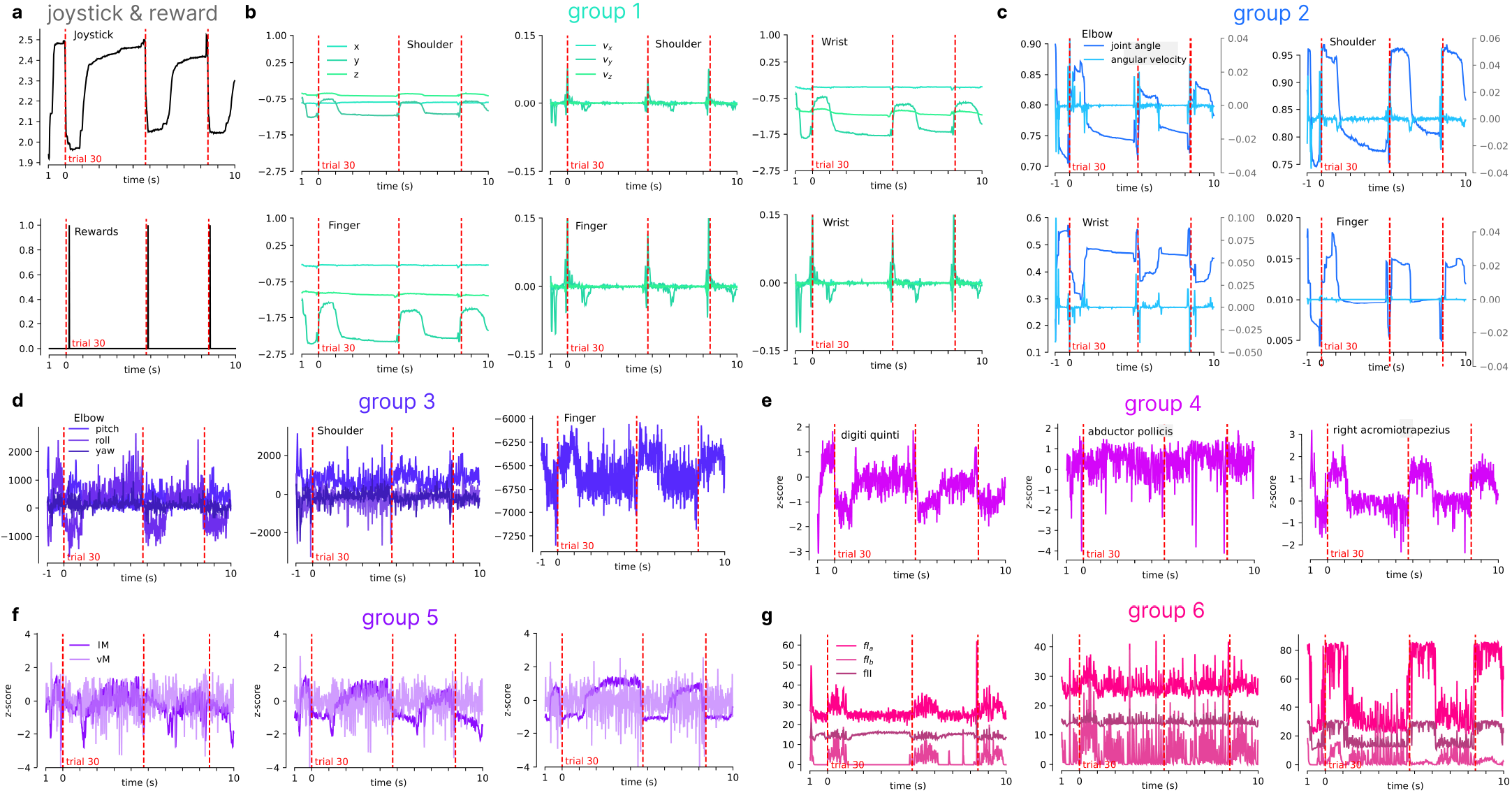
Example GLM regressors. Example temporal traces (10s, corresponding to approximately three trials in the baseline condition for an example session) for kinematic features used as regressors in the GLM analysis. Features are thematically clustered into groups: **a**, external variables such as joystick y-coordinate and reward delivery time; **b**, tracked keypoints 3D position & velocity; **c**, joint position & velocity; **d**, joint torques, **e**, muscle-space activations; **f**, muscle length & velocity, **g**, proprioceptive signals (muscle spindles & Golgi tendon organs).

**Figure S7.**
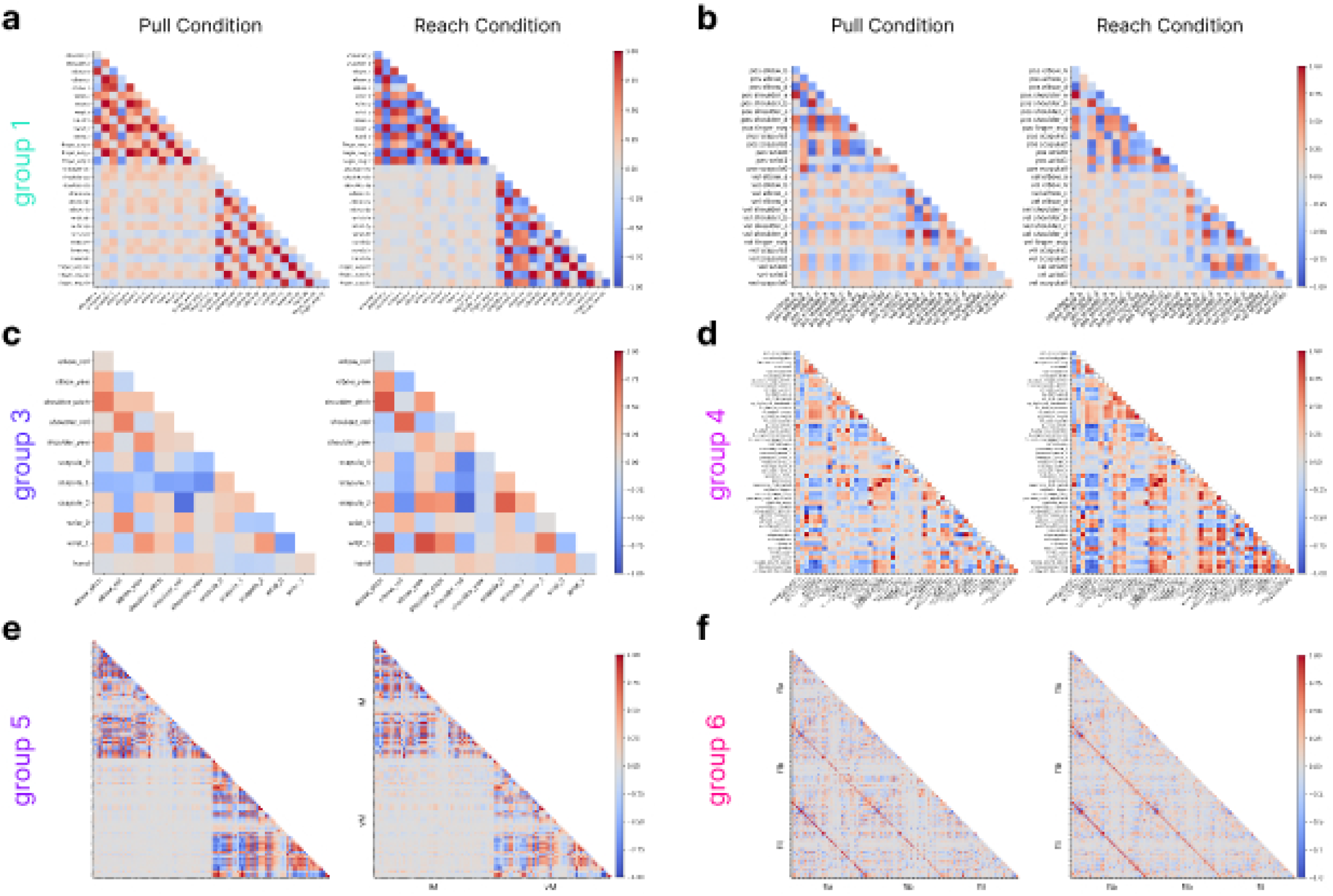
GLM regressors correlations. We measured the correlations between the kinematics features used as regressors in the GLM analysis. We computed the features correlation separately during pull (n=435, left columns) and reach (n=342, right columns) epochs. Features are thematically clustered into groups: **a**, external variables such as joystick y-coordinate and reward delivery; **b**, tracked keypoints 3D position & velocity; **c**, joint position & velocity; **d**, joint torques, **e**, muscle-space activations; **f**, muscle length & velocity, **g**, proprioceptive signals (muscle spindles & Golgi tendon organs).

**Figure S8.**
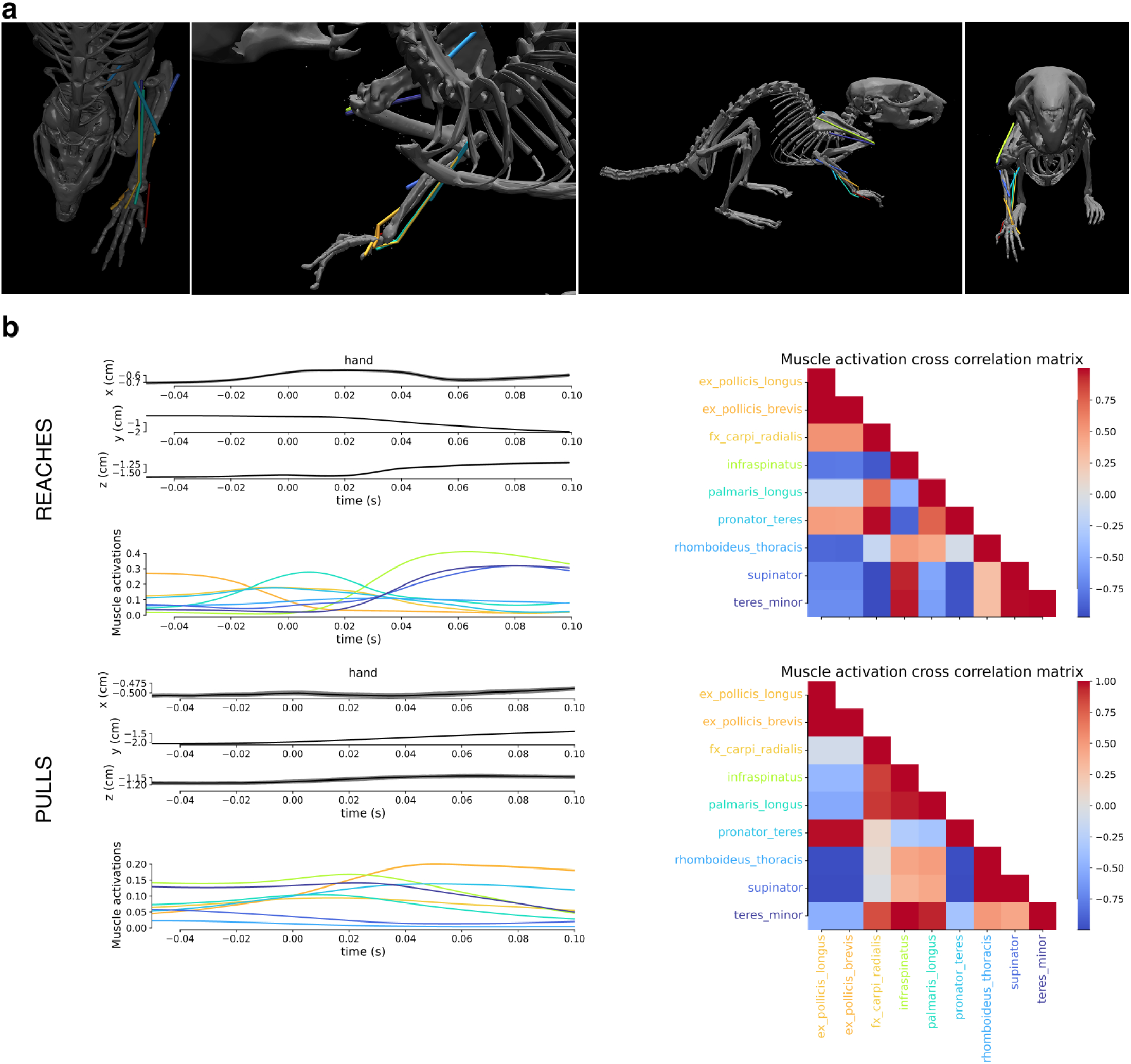
Muscles most tuned to neurons in M1/S1 based on GLMs. **a**, Images of the muscles that were most tuned based on the GLM analysis. **b**, We measured the correlation of muscles during pull (n=435) and reach (n=342) epochs and then computed the average correlation over the time-window shown. Within each panel the hand x, y, and z mean position and 95% confidence intervals are shown on the top left, bottom left is the mean muscle activations (see Methods), and right is the correlation matrix.

**Figure S9.**
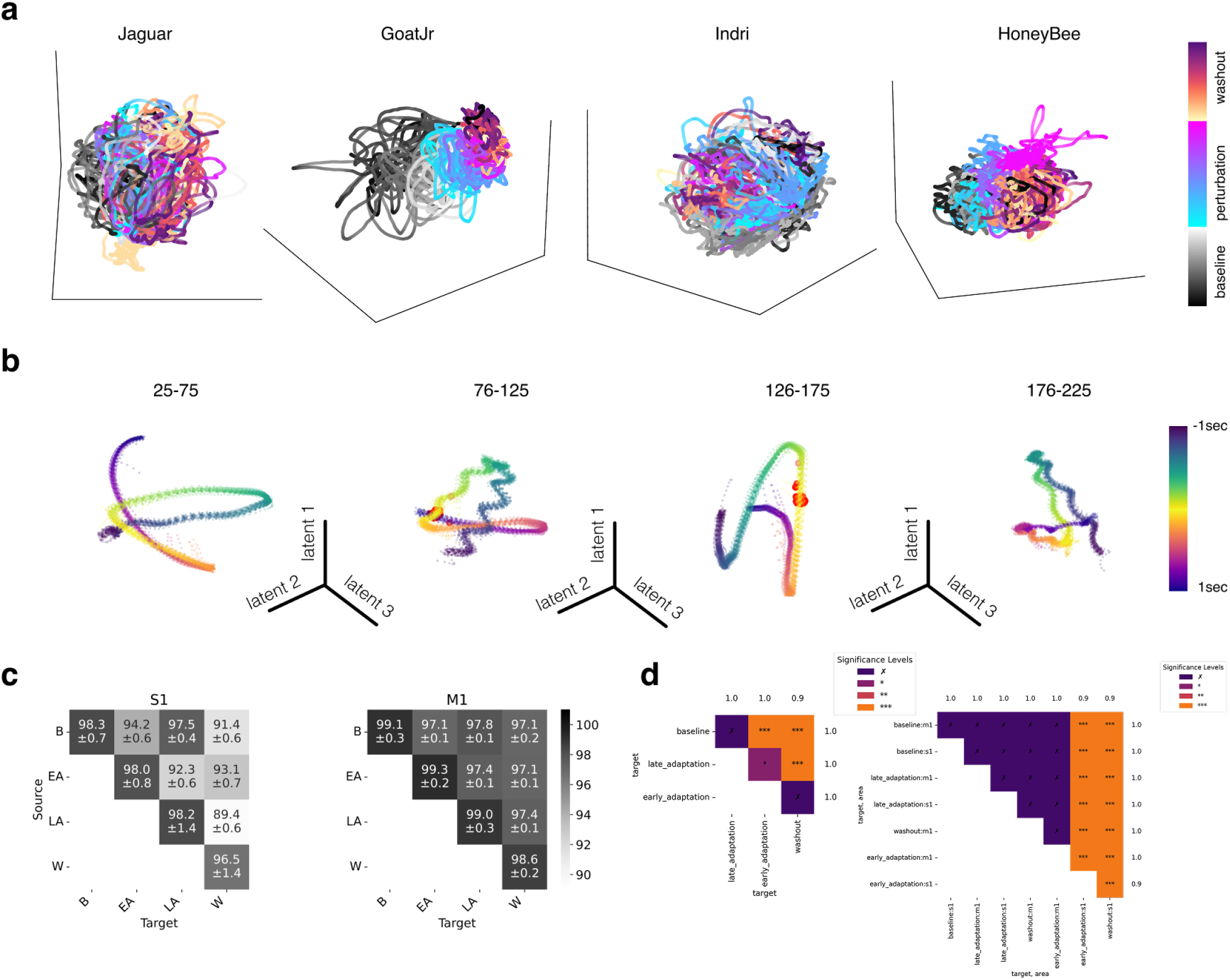
Adaptation CEBRA embeddings. **a**, Individual sessions were fit with CEBRA-Time. Specially, we used the ‘offset10-model-mse’ model, batchsize = 512, learning rate = 5e-5, temperature = 0.01, output_dimension = 3, distance = euclidean, and time_offsets = 10. It is colored by the time within each block. **b**, Using trial-time, we fit CEBRA-Behavior models from 50-trial blocks from baseline, early perturbation, late perturbation, and washout. Shown are the embeddings from 6D model (3 randomly selected are shown), which were used to make quantitative comparisons in Figure 2i. **c**, Consistency of runs across n=10 CEBRA models per epoch. Here, we note that M1 embeddings are more consistent and S1 across any epoch comparison (B is baseline, EA is early adaptation, LA is late adaptation, and W is washout).

**Figure S10.**
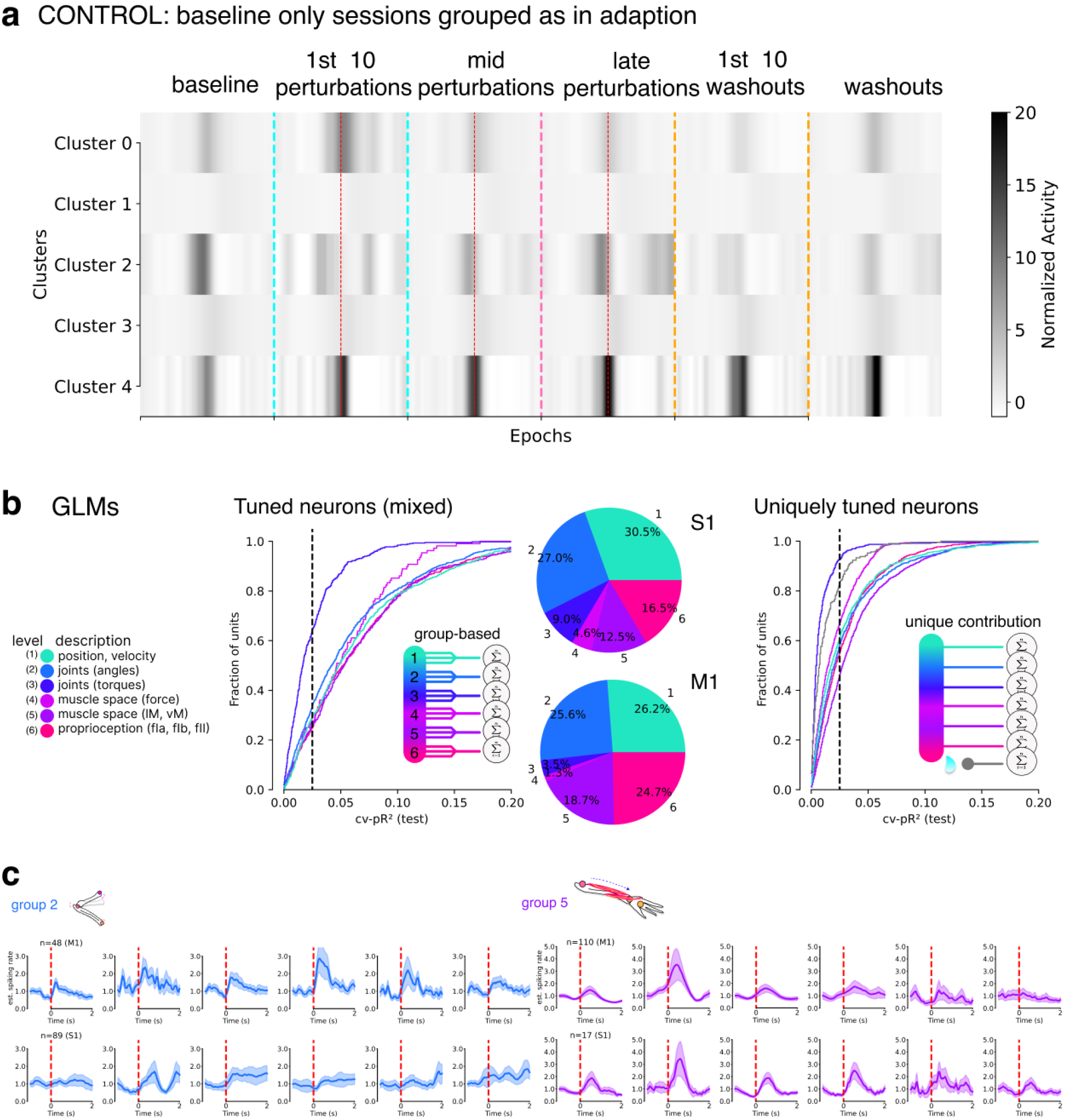
Clustering baseline data, GLMs during adaptation, and group-based PSTHs. **a**, Baseline-only sessions were used as a control for the clustering. Namely, there was no force-field perturbation, but we used the same PCA and hierarchical clustering as in Figure 7a. Here, unlike in the adaptation task, we do not see specific clusters that map to computations like prediction errors. **b**, GLM fits of the neurons recorded during the adaptation task sessions. Pie charts show the distribution of group-tuning (only considering the significantly tuned (*cv* − *pR*^2^ *>* 0.025) neurons). Far right shows the unique contribution. **c**, Neurons within Groups 2 and 5 in S1 or M1 (as noted) computed PSTHs per block, as computed in Figure 7d. Note, groups 3 had less than 15, and group 4 had less than 3 neurons, thus are not included. Statistics (Friedman test and permutation test) are available in Supplemental Tables S2, S3.

**Figure S11.**
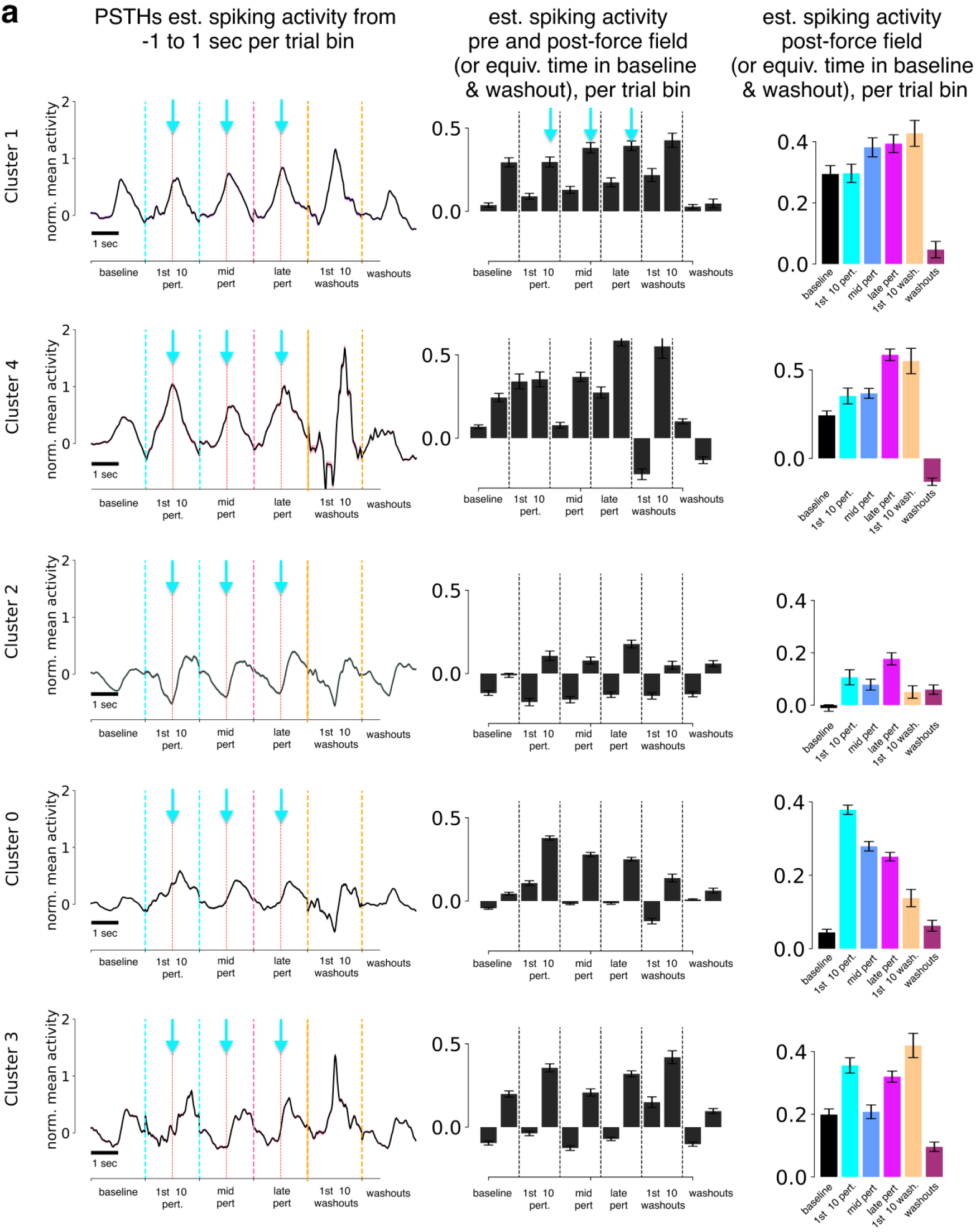
Clustering Adaptation Data. **a**, Left: the PSTH per time bin after clustering. Middle: the mean estimated spiking rates pre and post force-field, or equivalent time bin in baseline and washout. Right: only the postis plotted and re-colored to match the colormap used for adaptation sessions throughout the paper. These plots are then shown in Figure 7b.

## Statistics

**Table S1.**
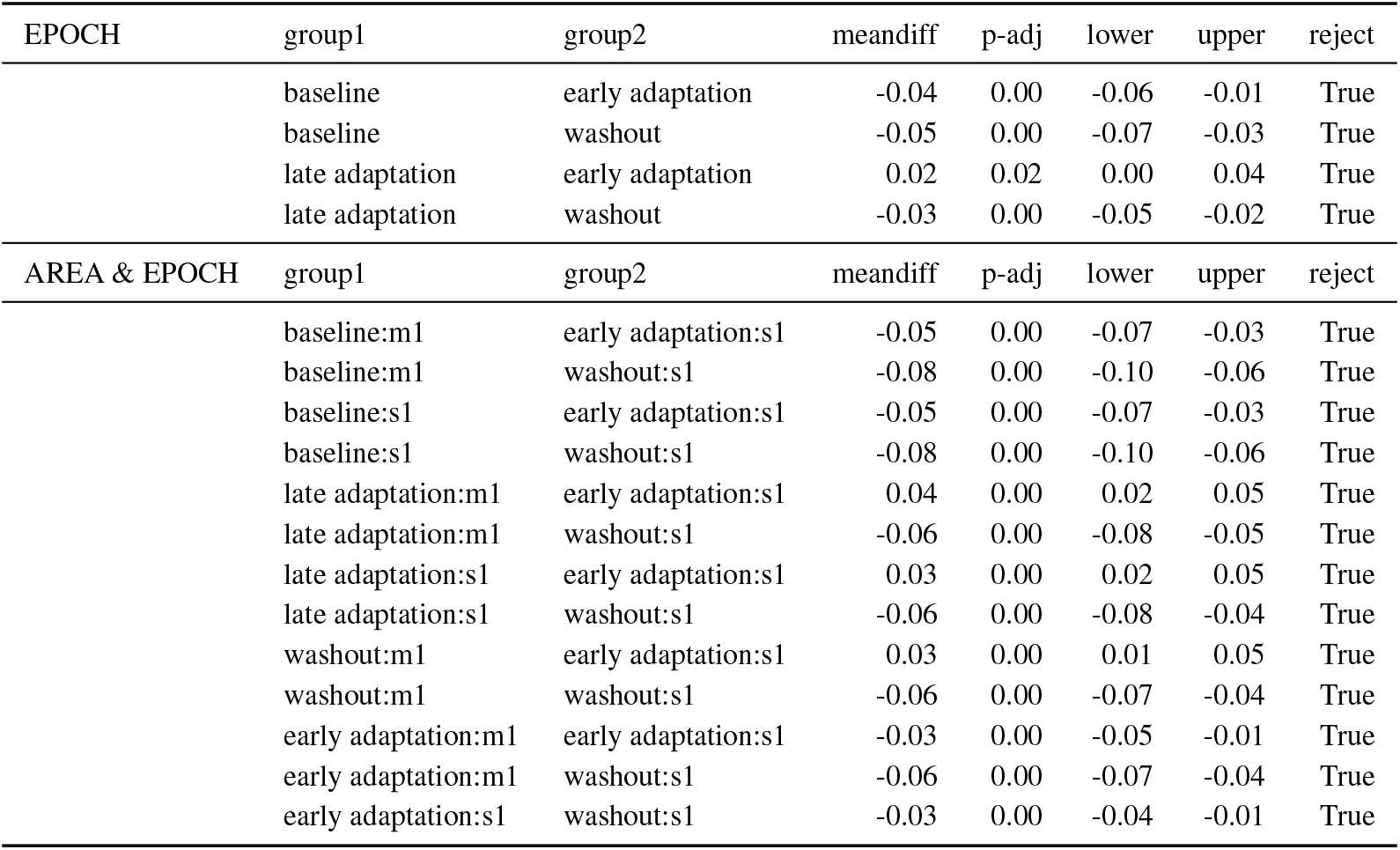
ANOVA with post-hoc Tukey significance testing. See also Suppl. Figure S9d.

| EPOCH | group1 | group2 | meandiff | p-adj | lower | upper | reject |
| --- | --- | --- | --- | --- | --- | --- | --- |
|  | baseline | early adaptation | -0.04 | 0.00 | -0.06 | -0.01 | True |
|  | baseline | washout | -0.05 | 0.00 | -0.07 | -0.03 | True |
|  | late adaptation | early adaptation | 0.02 | 0.02 | 0.00 | 0.04 | True |
|  | late adaptation | washout | -0.03 | 0.00 | -0.05 | -0.02 | True |
| AREA & EPOCH | group1 | group2 | meandiff | p-adj | lower | upper | reject |
|  | baseline:m1 | early adaptation:s1 | -0.05 | 0.00 | -0.07 | -0.03 | True |
|  | baseline:m1 | washout:s1 | -0.08 | 0.00 | -0.10 | -0.06 | True |
|  | baseline:s1 | early adaptation:s1 | -0.05 | 0.00 | -0.07 | -0.03 | True |
|  | baseline:s1 | washout:s1 | -0.08 | 0.00 | -0.10 | -0.06 | True |
|  | late adaptation:m1 | early adaptation:s1 | 0.04 | 0.00 | 0.02 | 0.05 | True |
|  | late adaptation:m1 | washout:s1 | -0.06 | 0.00 | -0.08 | -0.05 | True |
|  | late adaptation:s1 | early adaptation:s1 | 0.03 | 0.00 | 0.02 | 0.05 | True |
|  | late adaptation:s1 | washout:s1 | -0.06 | 0.00 | -0.08 | -0.04 | True |
|  | washout:m1 | early adaptation:s1 | 0.03 | 0.00 | 0.01 | 0.05 | True |
|  | washout:m1 | washout:s1 | -0.06 | 0.00 | -0.07 | -0.04 | True |
|  | early adaptation:m1 | early adaptation:s1 | -0.03 | 0.00 | -0.05 | -0.01 | True |
|  | early adaptation:m1 | washout:s1 | -0.06 | 0.00 | -0.07 | -0.04 | True |
|  | early adaptation:s1 | washout:s1 | -0.03 | 0.00 | -0.04 | -0.01 | True |

**Table S2.** Related to Figures 7d S11c: Friedman Test Results. Each group is tested against all baseline data. Groups 3 and 4 were not considered due to very low numbers of neurons.

| Region | Neuron Count | Group | $\chi^2$ | p-value | Significant |
| --- | --- | --- | --- | --- | --- |
| M1 | 406 | 0 (non-GLM tuned) | 454.387051 | $5.56 \times 10^{-96}$ | True |
| M1 | 101 | 1 | 12.818953 | $2.51 \times 10^{-2}$ | True |
| M1 | 48 | 2 | 16.523810 | $5.50 \times 10^{-3}$ | True |
| M1 | 6 | 3 | 12.380952 | $2.99 \times 10^{-2}$ | True |
| M1 | 2 | 4 | 144.800000 | $1.71 \times 10^{-29}$ | True |
| M1 | 110 | 5 | 2.000000 | $8.49 \times 10^{-1}$ | False |
| M1 | 116 | 6 | 101.073892 | $3.14 \times 10^{-20}$ | True |
| S1 | 425 | 0 (non-GLM tuned) | 35.041008 | $1.48 \times 10^{-6}$ | True |
| S1 | 325 | 1 | 80.570989 | $6.37 \times 10^{-16}$ | True |
| S1 | 89 | 2 | 16.024077 | $6.78 \times 10^{-3}$ | True |
| S1 | 15 | 3 | 1.704762 | $8.88 \times 10^{-1}$ | False |
| S1 | 3 | 4 | 10.949580 | $5.24 \times 10^{-2}$ | False |
| S1 | 17 | 5 | 1.095238 | $9.55 \times 10^{-1}$ | False |
| S1 | 34 | 6 | 16.151261 | $6.43 \times 10^{-3}$ | True |

**Table S3.**
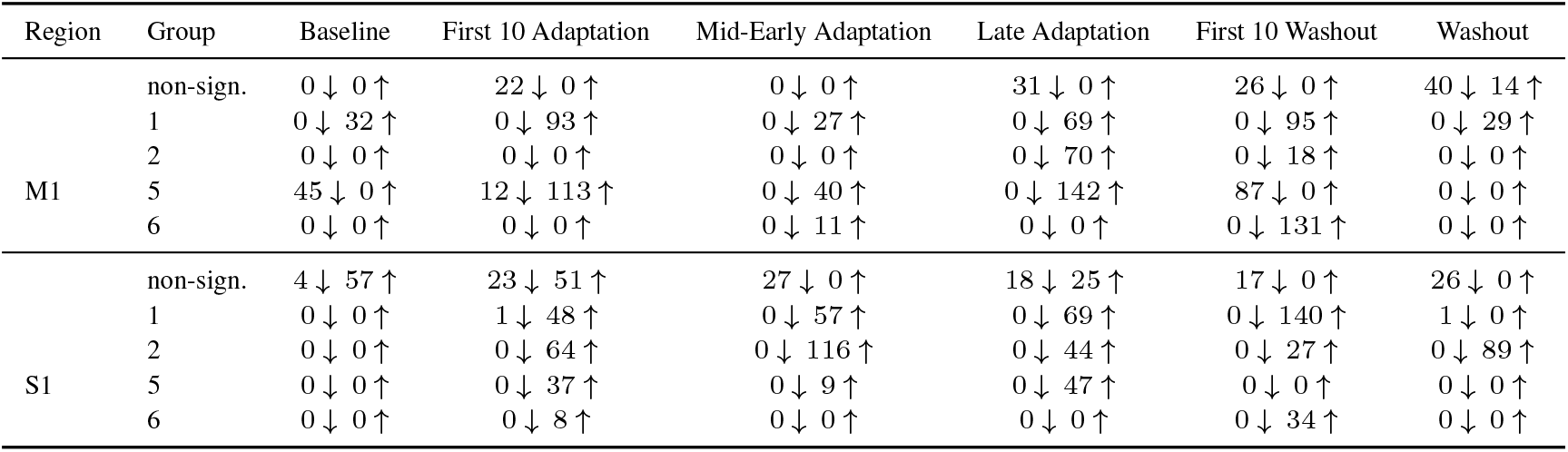
Related to Figures 7d S11c: Two-sided permutation testing (n=1000 bootstrapped samples) per time-bin, against baseline activity of all 1697 neurons. Counts of bins that are decreasing (*↓*) or increasing (*↑*) activity across epochs by region and group. Groups 3 and 4 were not considered due to very low numbers of neurons.

## Footnotes

0 We use the term sensorimotor vs. sensory as we found the prediction error signature in both the motor and somatosensory cortex

## References

1. Mackenzie Weygandt Mathis, Alexander Mathis, and Naoshige Uchida. Somatosensory Cortex Plays an Essential Role in Forelimb Motor Adaptation in Mice. Neuron, 93(6):1493–1503.e6, March 2017. ISSN 0896-6273. doi: 10.1016/j.neuron.2017.02.049.

2. Neeraj Kumar, Timothy F Manning, and David J. Ostry. Somatosensory cortex participates in the consolidation of human motor memory. PLoS Biology, 17, 2019.

3. Shahryar Ebrahimi and David J. Ostry. The human somatosensory cortex contributes to the encoding of newly learned movements. Proceedings of the National Academy of Sciences of the United States of America, 121, 2024.

4. Chiang-Shan Ray Li, Camillo Padoa-Schioppa, and Emilio Bizzi. Neuronal Correlates of Motor Performance and Motor Learning in the Primary Motor Cortex of Monkeys Adapting to an External Force Field. Neuron, 30(2):593–607, May 2001. ISSN 0896-6273. doi: 10.1016/S0896-6273(01)00301-4.

5. Risa Kawai, Timothy Markman, Rajesh Poddar, Raymond Ko, Antoniu L. Fantana, Ashesh K. Dhawale, Adam R. Kampff, and Bence P. Ölveczky. Motor Cortex Is Required for Learning but Not for Executing a Motor Skill. Neuron, 86(3):800–812, May 2015. ISSN 0896-6273. doi: 10.1016/j.neuron.2015.03.024.

6. J. Andrew Pruszynski, Isaac Kurtzer, Joseph Y. Nashed, Mohsen Omrani, Brenda Brouwer, and Stephen H. Scott. Primary motor cortex underlies multi-joint integration for fast feedback control. Nature, 478(7369):387–390, October 2011. ISSN 0028-0836. doi: 10.1038/nature10436.

7. RJ Nudo, GW Milliken, W Jenkins, and M. M. Merzenich. Use-dependent alterations of movement representations in primary motor cortex of adult squirrel monkeys. In Journal of Neuroscience, 1996.

8. Andrew E. Papale and Bryan M. Hooks. Circuit changes in motor cortex during motor skill learning. Neuroscience, 368:283–297, 2018.

9. Stephen H. Scott. The computational and neural basis of voluntary motor control and planning. Trends in Cognitive Sciences, 16(11):541–549, November 2012. ISSN 1364-6613. doi: 10.1016/j.tics.2012.09.008.

10. KA Thoroughman and R Shadmehr. Electromyographic correlates of learning an internal model of reaching movements. J. Neurosci., 19(19):8573–8588, October 1999. ISSN 1529-2401.

11. N. Hogan. An organizing principle for a class of voluntary movements. J. Neurosci., 4(11):2745–2754, November 1984. ISSN 0270-6474.

12. R. Shadmehr and F. A. Mussa-Ivaldi. Adaptive representation of dynamics during learning of a motor task. J. Neurosci., 14(5):3208–3224, May 1994. ISSN 0270-6474, 1529-2401.

13. J. R. Lackner and P. Dizio. Rapid adaptation to Coriolis force perturbations of arm trajectory. Journal of Neurophysiology, 72(1):299–313, July 1994.

14. John W Krakauer and Pietro Mazzoni. Human sensorimotor learning: adaptation, skill, and beyond. Current Opinion in Neurobiology, 21(4):636–644, August 2011. ISSN 0959-4388. doi: 10.1016/j.conb.2011.06.012.

15. Jonathan S. Tsay, Hrach Asmerian, Laura T. Germine, Jeremy B Wilmer, Richard B. Ivry, and Ken Nakayama. Large-scale citizen science reveals predictors of sensorimotor adaptation. Nature human behaviour, 2024.

16. Stephen H. Scott. A functional taxonomy of bottom-up sensory feedback processing for motor actions. Trends in Neurosciences, 39:512–526, 2016.

17. Emanuel Todorov and Michael I. Jordan. Optimal feedback control as a theory of motor coordination. Nat Neurosci, 5(11):1226–1235, November 2002. ISSN 1097-6256. doi: 10.1038/nn963.

18. D. M. Wolpert, R. C. Miall, and M. Kawato. Internal models in the cerebellum. Trends Cogn. Sci. (Regul. Ed.), 2(9):338–347, September 1998. ISSN 1364-6613.

19. Mitsuo Kawato. Internal models for motor control and trajectory planning. Current Opinion in Neurobio, 9(6):718–727, 1999.

20. Michael S. Fine and Kurt A. Thoroughman. Motor Adaptation to Single Force Pulses: Sensitive to Direction but Insensitive to Within-Movement Pulse Placement and Magnitude. Journal of Neurophysiology, 96(2):710–720, August 2006. ISSN 0022-3077, 1522-1598. doi: 10.1152/jn.00215.2006.

21. Stephen H. Scott. Optimal feedback control and the neural basis of volitional motor control. Nat Rev Neurosci, 5(7):532–546, July 2004. ISSN 1471-003X. doi: 10.1038/nrn1427.

22. Reza Shadmehr and John W Krakauer. A computational neuroanatomy for motor control. Exp Brain Res, 185(3):359–381, March 2008. ISSN 1432-1106. doi: 10.1007/s00221-008-1280-5.

23. Daniel M. Wolpert, Jörn Diedrichsen, and J. Randall Flanagan. Principles of sensorimotor learning. Nat Rev Neurosci, 12(12):739–751, December 2011. ISSN 1471-003X. doi: 10.1038/nrn3112.

24. Georg B. Keller, Tobias Bonhoeffer, and Mark Hübener. Sensorimotor Mismatch Signals in Primary Visual Cortex of the Behaving Mouse. Neuron, 74(5):809–815, June 2012. ISSN 0896-6273. doi: 10.1016/j.neuron.2012.03.040.

25. Rebecca Jordan and Georg B. Keller. Opposing influence of top-down and bottom-up input on excitatory layer 2/3 neurons in mouse primary visual cortex. Neuron, 108:1194–1206.e5, 2020.

26. Marcus Leinweber, Daniel R. Ward, Jan M. Sobczak, Alexander Attinger, and Georg B. Keller. A sensorimotor circuit in mouse cortex for visual flow predictions. Neuron, 95(6):1420–1432.e5, 2017.

27. Jonathan Green, Carissa A Bruno, Lisa Traunmuller, Jennifer Ding, Sinia Hrvatin, Daniel E Wilson, Thomas Khodadad, Jonathan Samuels, Michael Eldon Greenberg, and Christopher D. Harvey. A cell-type-specific error-correction signal in the posterior parietal cortex. Nature, 620:366 – 373, 2023.

28. Pawel Zmarz and Georg B. Keller. Mismatch receptive fields in mouse visual cortex. Neuron, 92:766–772, 2016.

29. Georg B. Keller and Philipp Sterzer. Predictive processing: A circuit approach to psychosis. Annual review of neuroscience, 2024.

30. Mackenzie W. Mathis. The neocortical column as a universal template for perception and world-model learning. Nature Reviews Neuroscience, 24:3, 2023.

31. Rebecca Jordan and Georg B Keller. The locus coeruleus broadcasts prediction errors across the cortex to promote sensorimotor plasticity. eLife, 12:RP85111, jun 2023. ISSN 2050-084X. doi: 10.7554/eLife.85111.

32. Nicholas J. Audette, WenXi Zhou, Alessandro La Chioma, and David M. Schneider. Precise movement-based predictions in the mouse auditory cortex. Current Biology, 32(22):4925–4940.e6, November 2022. ISSN 0960-9822. doi: 10.1016/j.cub.2022.09.064.

33. Nicholas J. Audette and David M. Schneider. Stimulus-specific prediction error neurons in mouse auditory cortex. Journal of Neuroscience, 43 (43):7119–7129, 2023. ISSN 0270-6474. doi: 10.1523/JNEUROSCI.0512-23.2023.

34. Emanuel Todorov. Direct cortical control of muscle activation in voluntary arm movements: a model. Nat Neurosci, 3(4):391–398, April 2000. ISSN 1097-6256. doi: 10.1038/73964.

35. Pietro Mazzoni and John W. Krakauer. An Implicit Plan Overrides an Explicit Strategy during Visuomotor Adaptation. J. Neurosci., 26(14):3642–3645, April 2006. ISSN 0270-6474, 1529-2401. doi: 10.1523/JNEUROSCI.5317-05.2006.

36. Michael I. Jordan and David E. Rumelhart. Forward models: Supervised learning with a distal teacher. Cogn. Sci., 16:307–354, 1992.

37. John W. Krakauer, Maria Felice Ghilardi, and Claude Ghez. Independent learning of internal models for kinematic and dynamic control of reaching. Nature Neuroscience, 2:1026–1031, 1999.

38. Alkis M. Hadjiosif, John W. Krakauer, and Adrian M. Haith. Did we get sensorimotor adaptation wrong? implicit adaptation as direct policy updating rather than forward-model-based learning. The Journal of Neuroscience, 41:2747 – 2761, 2021.

39. Apostolos P. Georgopoulos, Andrew B. Schwartz, and RE Kettner. Neuronal population coding of movement direction. Science, 233 4771: 1416–9, 1986.

40. James M. Goodman, Gregg Tabot, Alex S. Lee, Aneesha K. Suresh, Alexander T. Rajan, Nicholas G. Hatsopoulos, and Sliman J. Bensmaia. Postural representations of the hand in the primate sensorimotor cortex. Neuron, 104:1000–1009.e7, 2019.

41. Stephen Scott and John F. Kalaska. Reaching movements with similar hand paths but different arm orientations. i. activity of individual cells in motor cortex. Journal of neurophysiology, 77 2:826–52, 1997.

42. Stephen H. Scott. Inconvenient truths about neural processing in primary motor cortex. The Journal of Physiology, 586, 2008.

43. Yoshikazu Isomura, Rie Harukuni, Takashi Takekawa, Hidenori Aizawa, and Tomoki Fukai. Microcircuitry coordination of cortical motor information in self-initiation of voluntary movements. Nature Neuroscience, 12:1586–1593, 2009.

44. Andrew Miri, Claire L. Warriner, Jeffrey S. Seely, Gamaleldin F. Elsayed, John P. Cunningham, Mark M. Churchland, and Thomas M. Jessell. Behaviorally selective engagement of short-latency effector pathways by motor cortex. Neuron, 95:683–696.e11, 2017.

45. Takaki Komiyama, Takashi R Sato, Daniel H. O’Connor, Ying-Xin Zhang, Daniel Huber, Bryan M. Hooks, Mariano Gabitto, and Karel Svoboda. Learning-related fine-scale specificity imaged in motor cortex circuits of behaving mice. Nature, 464:1182–1186, 2010.

46. Andrew J. Peters, Simon X. Chen, and Takaki Komiyama. Emergence of reproducible spatiotemporal activity during motor learning. Nature, 510:263–267, 2014.

47. Eun Jung Hwang, Jeffrey E. Dahlen, Yvonne Yuling Hu, Karina Aguilar, Bin Yu, Madan Mukundan, Akinori Mitani, and Takaki Komiyama. Disen-gagement of motor cortex from movement control during long-term learning. Science Advances, 5, 2019.

48. Raeed H. Chowdhury, Joshua I. Glaser, and Lee E. Miller. Area 2 of primary somatosensory cortex encodes kinematics of the whole arm. eLife, 9, 2019.

49. Ignacio Alonso, Irina Scheer, Melanie Palacio-Manzano, Noémie Frézel-Jacob, Antoine Philippides, and Mario Prsa. Peripersonal encoding of forelimb proprioception in the mouse somatosensory cortex. Nature Communications, 14, 2023.

50. Alessandro Marin Vargas, Axel Bisi, Alberto S Chiappa, Chris Versteeg, Lee E Miller, and Alexander Mathis. Task-driven neural network models predict neural dynamics of proprioception. Cell, 187(7):1745–1761, 2024.

51. Hermano Igo Krebs, Neville Hogan, Mindy L. Aisen, and Bruce T. Volpe. Robot-Aided Neurorehabilitation. IEEE Trans Rehabil Eng, 6(1):75–87, March 1998. ISSN 1063-6528.

52. Mohsen Omrani, Chantelle D. Murnaghan, J. Andrew Pruszynski, and Stephen H. Scott. Distributed task-specific processing of somatosensory feedback for voluntary motor control. eLife, 5:e13141, April 2016. ISSN 2050-084X. doi: 10.7554/eLife.13141.

53. Saurabh Vyas, Daniel J. O’Shea, Stephen I. Ryu, and Krishna V. Shenoy. Causal role of motor preparation during error-driven learning. Neuron, 2020.

54. Matthew I. Becker, Dylan J. Calame, Julia Wrobel, and Abigail L. Person. Online control of reach accuracy in mice. Journal of neurophysiology, 2020.

55. Wuzhou Yang, Harsh Kanodia, and Silvia Arber. Structural and functional map for forelimb movement phases between cortex and medulla. Cell, 186:162–177.e18, 2023.

56. Kelly A. Tennant, DeAnna L. Adkins, Nicole A. Donlan, Aaron L. Asay, Nagheme J. Thomas, Jeffrey A. Kleim, and Theresa A. Jones. The organization of the forelimb representation of the c57bl/6 mouse motor cortex as defined by intracortical microstimulation and cytoarchitecture. Cerebral cortex, 21 4:865–76, 2011.

57. Karin Morandell and Daniel Huber. The role of forelimb motor cortex areas in goal directed action in mice. Scientific Reports, 7, 2017.

58. D. Huber, D. A. Gutnisky, S. Peron, D. H. O’Connor, J. S. Wiegert, L. Tian, T. G. Oertner, L. L. Looger, and K. Svoboda. Multiple dynamic representations in the motor cortex during sensorimotor learning. Nature, 484(7395):473–478, April 2012. ISSN 0028-0836, 1476-4687. doi: 10.1038/nature11039.

59. Mark J. Wagner, Joan Savall, Tony Hyun Kim, Mark J. Schnitzer, and Liqun Luo. Skilled reaching tasks for head-fixed mice using a robotic manipulandum. Nature Protocols, 15:1237 – 1254, 2020.

60. Spencer Bowles, Jordan Hickman, Xiaoyu Peng, William R. Williamson, Rongcheng Huang, Kayden A. Washington, Dane C Donegan, and Cristin G. Welle. Vagus nerve stimulation drives selective circuit modulation through cholinergic reinforcement. Neuron, 110:2867–2885.e7, 2022.

61. John Martin Barrett, Martinna G. Raineri Tapies, and Gordon M. G. Shepherd. Manual dexterity of mice during food-handling involves the thumb and a set of fast basic movements. PLoS ONE, 15, 2019.

62. Riichiro Hira, Fuki Ohkubo, Katsuya Ozawa, Yoshikazu Isomura, Kazuo Kitamura, Masanobu Kano, Haruo Kasai, and Masanori Matsuzaki. Spatiotemporal dynamics of functional clusters of neurons in the mouse motor cortex during a voluntary movement. The Journal of Neuroscience, 33:1377 – 1390, 2013.

63. Nicholas James Sofroniew, Daniel Flickinger, Jonathan King, and Karel Svoboda. A large field of view two-photon mesoscope with subcellular resolution for in vivo imaging. eLife, 5, 2016.

64. Jun Izawa and Reza Shadmehr. Learning from Sensory and Reward Prediction Errors during Motor Adaptation. PLoS Comput Biol, 7(3):e1002012, March 2011. doi: 10.1371/journal.pcbi.1002012.

65. Steffen Schneider, Jin Hwa Lee, and Mackenzie W. Mathis. Learnable latent embeddings for joint behavioural and neural analysis. Nature, 617:360 – 368, 2023.

66. Mark Churchland et al. Neural population dynamics during reaching. Nature, 2012.

67. Mark M. Churchland and Krishna V Shenoy. Preparatory activity and the expansive null-space. Nature reviews. Neuroscience, 2024.

68. Bryce Chung, Muneeb Zia, Kyle A. Thomas, Jonathan A. Michaels, Amanda L. Jacob, Andrea R. Pack, Matt Williams, Kailash Nagapudi, Lay Heng Teng, Eduardo Arrambide, Logan Ouellette, Nicole F. Oey, Rhuna Gibbs, Philip Anschutz, Jiaao Lu, Yu Ping Wu, Mehrdad Kashefi, Tomomichi Oya, Rhonda Kersten, Alice C. Mosberger, Sean O’Connell, Runming Wang, Hugo Marques, Ana Rita Mendes, Constanze Lenschow, Gayathri Kondakath, Jeong Jun Kim, William P Olson, Kiara N. Quinn, Pierce Perkins, Graziana Gatto, Ayesha Thanawalla, Susan K. Coltman, Taegyo Kim, Trevor Smith, Benjamin I. Binder-Markey, Martin Zaback, Christopher K. Thompson, Simon F. Giszter, Abigail L. Person, Martyn D. Goulding, Eiman Azim, Nitish V. Thakor, Daniel H. O’Connor, Barry Andrew Trimmer, Susana Q. Lima, Megan R. Carey, Chethan Pandarinath, Rui M. Costa, J. Andrew Pruszynski, Muhannad S. Bakir, and Samuel J. Sober. Myomatrix arrays for high-definition muscle recording. eLife, 12, 2023.

69. Eunice Chace Greene. Anatomy of the rat. Transactions of the American Philosophical Society, 27:iii–370, 1935.

70. Takamasa Iwaki, Hiroshi Yamashita, Toshiyuki Hayakawa, et al. Color atlas of sectional anatomy of the mouse. Braintree Scientific Inc, 2005.

71. Albert King. A review of biomechanical models. Journal of biomechanical engineering, 106 2:97–104, 1984.

72. Emilie Mathieu, Sylvain Crémoux, David Duvivier, David Amarantini, and Philippe Pudlo. Biomechanical modeling for the estimation of muscle forces: toward a common language in biomechanics, medical engineering, and neurosciences. Journal of NeuroEngineering and Rehabilitation, 20, 2023.

73. Scott L. Delp, Frank C. Anderson, Allison S. Arnold, Peter Loan, Ayman Habib, Chand T. John, Eran Guendelman, and Darryl G. Thelen. Opensim: Open-source software to create and analyze dynamic simulations of movement. IEEE Transactions on Biomedical Engineering, 54:1940–1950, 2007.

74. Emanuel Todorov, Tom Erez, and Yuval Tassa. Mujoco: A physics engine for model-based control. In 2012 IEEE/RSJ international conference on intelligent robots and systems, pages 5026–5033. IEEE, 2012.

75. Kai J. Sandbrink, Pranav Mamidanna, Claudio Michaelis, Matthias Bethge, Mackenzie W. Mathis, and Alexander Mathis. Contrasting action and posture coding with hierarchical deep neural network models of proprioception. eLife, 12, 2023.

76. Olivier Codol, Jonathan A. Michaels, Mehrdad Kashefi, J. Andrew Pruszynski, and Paul L. Gribble. Motornet: a python toolbox for controlling differentiable biomechanical effectors with artificial neural networks. eLife, 2024.

77. Muhammad Noman Almani, John Lazzari, Andrea Chacon, and Shreya Saxena. µsim: A goal-driven framework for elucidating the neural control of movement through musculoskeletal modeling. bioRxiv, 2024.

78. Timothy P Lillicrap and Stephen H Scott. Preference distributions of primary motor cortex neurons reflect control solutions optimized for limb biomechanics. Neuron, 77(1):168–179, 2013.

79. Alberto Silvio Chiappa, Pablo Tano, Nisheet Patel, Abigaïl Ingster, Alexandre Pouget, and Alexander Mathis. Acquiring musculoskeletal skills with curriculum-based reinforcement learning. bioRxiv, 2024. doi: 10.1101/2024.01.24.577123.

80. Shravan Tata Ramalingasetty, Simon M. Danner, Jonathan Arreguit, Sergey N. Markin, Dimitri Rodarie, Claudia Kathe, Grégoire Courtine, Ilya A. Rybak, and Auke Jan Ijspeert. A whole-body musculoskeletal model of the mouse. IEEE access : practical innovations, open solutions, 9:163861 – 163881, 2021.

81. Alexander Mathis, Pranav Mamidanna, Kevin M Cury, Taiga Abe, Venkatesh N Murthy, Mackenzie Weygandt Mathis, and Matthias Bethge. Deeplabcut: markerless pose estimation of user-defined body parts with deep learning. Nature neuroscience, 21(9):1281–1289, 2018.

82. Tanmay Nath, Alexander Mathis, An Chi Chen, Amir Patel, Matthias Bethge, and Mackenzie W. Mathis. Using deeplabcut for 3d markerless pose estimation across species and behaviors. Nature Protocols, 14:2152 – 2176, 2019.

83. Markus Frey, Wesley Monteith-Finas, Travis DeWolf, Steffen Schneider, and Mackenzie W. Mathis. MouseArmTransformer: a transformer-based 3D lifting module for an adult mouse arm. Zenodo, July 2024. doi: 10.5281/zenodo.12673173.

84. Alain Frigon, Yann Thibaudier, and Marie-France Hurteau. Modulation of forelimb and hindlimb muscle activity during quadrupedal tied-belt and split-belt locomotion in intact cats. Neuroscience, 290:266–278, 2015.

85. Quanxin Wang, Song-Lin Ding, Yang Li, Joshua J. Royall, Hongkui Zeng, and Lydia Ng. The allen mouse brain common coordinate framework: A 3d reference atlas. Cell, 181:936–953.e20, 2020.

86. Eric R Kandel, James H Schwartz, and Thomas M Jessell. Principles of neural science. McGraw-Hill, Health Professions Division, New York, 2000. ISBN 0-8385-7701-6 978-0-8385-7701-1.

87. Diego E. Aldarondo, Josh Merel, Jesse D. Marshall, Leonard Hasenclever, Ugne Klibaite, Amanda Gellis, Yuval Tassa, Greg Wayne, Matthew M. Botvinick, and Bence P. Ölveczky. A virtual rodent predicts the structure of neural activity across behaviors. Nature, 2024.

88. Maurice A Smith, Ali Ghazizadeh, and Reza Shadmehr. Interacting Adaptive Processes with Different Timescales Underlie Short-Term Motor Learning. PLoS Biol, 4(6):e179, May 2006. doi: 10.1371/journal.pbio.0040179.

89. Ian E. Brown and James M. Bower. The Influence of Somatosensory Cortex on Climbing Fiber Responses in the Lateral Hemispheres of the Rat Cerebellum after Peripheral Tactile Stimulation. J. Neurosci., 22(15):6819–6829, August 2002. ISSN 0270-6474, 1529-2401.

90. Mohsen Omrani, Mohsen Omrani, Chantelle Dawn Murnaghan, J. Andrew Pruszynski, J. Andrew Pruszynski, and Stephen H. Scott. Distributed task-specific processing of somatosensory feedback for voluntary motor control. eLife, 5, 2016.

91. Mackenzie W. Mathis and Steffen Schneider. Motor control: Neural correlates of optimal feedback control theory. Current Biology, 31:R356–R358, 2021.

92. Matthew T. Kaufman, Mark M. Churchland, Stephen I. Ryu, and Krishna V. Shenoy. Cortical activity in the null space: permitting preparation without movement. Nature Neuroscience, 17(3):440–448, 2014. ISSN 1546-1726. doi: 10.1038/nn.3643.

93. David Eriksson, Mona Heiland, Artur Schneider, and Ilka Diester. Distinct dynamics of neuronal activity during concurrent motor planning and execution. Nature Communications, 12, 2021.

94. Barbara Feulner, Matthew G. Perich, Raeed H. Chowdhury, Lee E. Miller, Juan Alvaro Gallego, and Claudia Clopath. Small, correlated changes in synaptic connectivity may facilitate rapid motor learning. Nature Communications, 13, 2022.

95. Abigail A. Russo, Ramin Khajeh, Sean R. Bittner, Sean M. Perkins, John P. Cunningham, Laurence F. Abbott, and Mark M. Churchland. Neural trajectories in the supplementary motor area and motor cortex exhibit distinct geometries, compatible with different classes of computation. Neuron, 107:745–758.e6, 2020.

96. Abigail A. Russo, Sean R. Bittner, Sean M. Perkins, Jeffrey S. Seely, Brian M. London, Antonio H. Lara, Andrew Miri, Najja J. Marshall, Adam Kohn, Thomas M. Jessell, L. F. Abbott, John P. Cunningham, and Mark M. Churchland. Motor cortex embeds muscle-like commands in an untangled population response. Neuron, 97:953–966.e8, 2018.

97. Mackenzie W. Mathis, Adriana Perez Rotondo, Edward F. Chang, Andreas S. Tolias, and Alexander Mathis. Decoding the brain: From neural representations to mechanistic models. Cell, 2024.

98. Steffen Schneider, Rodrigo González Laiz, Anastasiia Filippova, Markus Frey, and Mackenzie W Mathis. Time-series attribution maps with regularized contrastive learning. In The 28th International Conference on Artificial Intelligence and Statistics, 2025.

99. Tony Hyun Kim, Yanping Zhang, Jérôme A. Lecoq, Juergen C. Jung, Jane Li, Hongkui Zeng, Cristopher M. Niell, and Mark J. Schnitzer. Long-term optical access to an estimated one million neurons in the live mouse cortex. Cell reports, 17 12:3385–3394, 2016.

100. Adam Paszke, Sam Gross, Francisco Massa, Adam Lerer, James Bradbury, Gregory Chanan, Trevor Killeen, Zeming Lin, Natalia Gimelshein, Luca Antiga, Alban Desmaison, Andreas Köpf, Edward Yang, Zach DeVito, Martin Raison, Alykhan Tejani, Sasank Chilamkurthy, Benoit Steiner, Lu Fang, Junjie Bai, and Soumith Chintala. Pytorch: An imperative style, high-performance deep learning library. ArXiv, abs/1912.01703, 2019.

101. Dimitri Yatsenko, Jacob Reimer, Alexander S. Ecker, Edgar Y. Walker, Fabian H Sinz, Philipp Berens, Andreas Hoenselaar, R. James Cotton, Athanassios S. Siapas, and Andreas Savas Tolias. Datajoint: managing big scientific data using matlab or python. bioRxiv, 2015.

102. J. Alexander Bae, Mahaly Baptiste, Agnes L. Bodor, Derrick Brittain, JoAnn Buchanan, D. Bumbarger, Manuel A. Castro, Brendan Celii, Erick Cobos, Forrest Collman, Nuno Maçarico da Costa, Sven Dorkenwald, Leila Elabbady, Paul G. Fahey, Tim Fliss, Emmanouil Froudakis, Jay Gager, Clare Gamlin, Akhilesh Halageri, James Hebditch, Zhen Jia, Chris S. Jordan, Daniel Kapner, Nico Kemnitz, Sam Kinn, Selden Koolman, Kai Kuehner, Kisuk Lee, Kai Li, Ran Lu, Thomas Macrina, Gayathri Mahalingam, Sarah McReynolds, Elanine Miranda, Eric Mitchell, Shanka Subhra Mondal, Merlin Moore, Shang Mu, Taliah Muhammad, Barak Nehoran, Oluwaseun Ogedengbe, Ch. Papadopoulos, Stelios Papadopoulos, Saumil S. Patel, Xaq Pitkow, Sergiy Popovych, Anthony Ramos, R. Clay Reid, Jacob Reimer, Casey M. Schneider-Mizell, H. Sebastian Seung, Benjamin Silverman, William M. Silversmith, Amy Robinson Sterling, Fabian H Sinz, Cameron L. Smith, Shelby K. Suckow, Zheng Huan Tan, Andreas Savas Tolias, Russel Torres, Nicholas L. Turner, Edgar Y. Walker, Tianyu Wang, Grace Williams, Susana Williams, Kyle Patrick Willie, Ryan Willie, William Wong, Jingpeng Wu, Chris Xu, Runzhe Yang, Dimitri Yatsenko, Fei Ye, Wenjing Yin, and Szi chieh Yu. Functional connectomics spanning multiple areas of mouse visual cortex. Nature, 640:435 – 447, 2025.

103. B. Srinivasa Reddy and Biswanath N. Chatterji. An fft-based technique for translation, rotation, and scale-invariant image registration. IEEE transactions on image processing : a publication of the IEEE Signal Processing Society, 5 8:1266–71, 1996.

104. Andrea Giovannucci, Johannes Friedrich, Pat Gunn, Jérémie Kalfon, Brandon L Brown, Sue Ann Koay, Jiannis Taxidis, Farzaneh Najafi, Jeffrey L. Gauthier, Pengcheng Zhou, Baljit S. Khakh, David W. Tank, Dmitri B. Chklovskii, and Eftychios A. Pnevmatikakis. Caiman an open source tool for scalable calcium imaging data analysis. eLife, 8, 2019.

105. Eftychios A. Pnevmatikakis, Daniel Soudry, Yuanjun Gao, Timothy A. Machado, Josh Merel, David Pfau, Thomas R. Reardon, Yu Mu, Clay Lacefield, Weijian Yang, Misha B. Ahrens, Randy M. Bruno, Thomas M. Jessell, Darcy S. Peterka, Rafael Yuste, and Liam Paninski. Simultaneous denoising, deconvolution, and demixing of calcium imaging data. Neuron, 89:285–299, 2016.

106. James P Charles, Ornella Cappellari, Andrew J Spence, Dominic J Wells, and John R Hutchinson. Muscle moment arms and sensitivity analysis of a mouse hindlimb musculoskeletal model. Journal of anatomy, 229(4):514–535, 2016.

107. Pierre Kibleur, Shravan R Tata, Nathan Greiner, Sara Conti, Beatrice Barra, Katie Zhuang, Melanie Kaeser, Auke Ijspeert, and Marco Capogrosso. Spatiotemporal maps of proprioceptive inputs to the cervical spinal cord during three-dimensional reaching and grasping. IEEE Transactions on Neural Systems and Rehabilitation Engineering, 28(7):1668–1677, 2020.

108. H.M. Hudson and R.S. Larkin. Accelerated image reconstruction using ordered subsets of projection data. IEEE Transactions on Medical Imaging, 13(4):601–609, 1994. doi: 10.1109/42.363108.

109. Andriy Fedorov, Reinhard Beichel, Jayashree Kalpathy-Cramer, Julien Finet, Jean-Christophe Fillion-Robin, Sonia Pujol, Christian Bauer, Dominique Jennings, Fiona Fennessy, Milan Sonka, et al. 3d slicer as an image computing platform for the quantitative imaging network. Magnetic resonance imaging, 30(9):1323–1341, 2012.

110. Arthur Prochazka and Monica Gorassini. Ensemble firing of muscle afferents recorded during normal locomotion in cats. The Journal of physiology, 507(1):293–304, 1998.

111. Arthur Prochazka and Monica Gorassini. Models of ensemble firing of muscle spindle afferents recorded during normal locomotion in cats. The Journal of physiology, 507(1):277–291, 1998.

112. Eduardo Martin Moraud, Marco Capogro:wsso, Emanuele Formento, Nikolaus Wenger, Jack DiGiovanna, Gregoire Courtine, and Silvestro Micera. Mechanisms underlying the neuromodulation of spinal circuits for correcting gait and balance deficits after spinal cord injury. Neuron, 89(4):814– 828, 2016.

113. Arthur Prochazka. Quantifying proprioception. Progress in brain research, 123:133–142, 1999.

114. Fanwang Meng, Michael Richer, Alireza Tehrani, Jonathan La, Taewon David Kim, Paul W. Ayers, and Farnaz Heidar-Zadeh. Procrustes: A python library to find transformations that maximize the similarity between matrices. Computer Physics Communications, 276(108334):1–37, 2022. ISSN 0010-4655. doi: 10.1016/j.cpc.2022.108334.

115. Tsuneo Yoshikawa. Manipulability of robotic mechanisms. The international journal of Robotics Research, 4(2):3–9, 1985.

116. Microsoft. Neural Network Intelligence. GitHub, 1 2021.

117. R. S Sutton and A. G Barto. Reinforcement learning: An introduction, volume 1. Cambridge Univ Press, 1998.

118. Max Berniker and Konrad Kording. Estimating the sources of motor errors for adaptation and generalization. Nat Neurosci, 11(12):1454–1461, December 2008. ISSN 1097-6256. doi: 10.1038/nn.2229.

119. R. E. Kalman. A New Approach to Linear Filtering and Prediction Problems. J. Basic Eng., 82(1):35–45, March 1960. ISSN 0098-2202. doi:10.1115/1.3662552.

120. Simon Musall, Matthew T. Kaufman, Ashley L. Juavinett, Steven Gluf, and Anne K. Churchland. Single-trial neural dynamics are dominated by richly varied movements. Nature neuroscience, 22:1677 – 1686, 2019.

121. Andrew J. Peters, Simon X. Chen, and Takaki Komiyama. Emergence of reproducible spatiotemporal activity during motor learning. Nature, 510 (7504):263–267, June 2014. ISSN 0028-0836. doi: 10.1038/nature13235.

122. Celia Benquet, Hossein Mirzaeri, Steffen Schneider, and Mackenzie W. Mathis. Unified cebra encoders for integrating neural recordings via behavioral alignment. bioRxiv, 2025.

123. Charles R. Harris, K. Jarrod Millman, Stéfan van der Walt, Ralf Gommers, Pauli Virtanen, David Cournapeau, Eric Wieser, Julian Taylor, Sebastian Berg, Nathaniel J. Smith, Robert Kern, Matti Picus, Stephan Hoyer, Marten H. van Kerkwijk, Matthew Brett, Allan Haldane, Jaime Fern’andez del R’io, Marcy Wiebe, Pearu Peterson, Pierre G’erard-Marchant, Kevin Sheppard, Tyler Reddy, Warren Weckesser, Hameer Abbasi, Christoph Gohlke, and Travis E. Oliphant. Array programming with numpy. Nature, 585:357 – 362, 2020.

124. Pauli Virtanen, Ralf Gommers, Travis E. Oliphant, Matt Haberland, Tyler Reddy, David Cournapeau, Evgeni Burovski, Pearu Peterson, Warren Weckesser, Jonathan Bright, Stéfan J. van der Walt, Matthew Brett, Joshua Wilson, K. Jarrod Millman, Nikolay Mayorov, Andrew R. J. Nelson, Eric Jones, Robert Kern, Eric Larson,J J Carey, İlhan Polat, Yu Feng, Eric W. Moore, Jake VanderPlas, Denis Laxalde, Josef Perktold, Robert Cimrman, Ian Henriksen, E. A. Quintero, Charles R Harris, Anne M. Archibald, Antônio H. Ribeiro, Fabian Pedregosa, Paul van Mulbregt, and SciPy 1.0 Contributors. SciPy 1.0: Fundamental algorithms for scientific computing in python. Nature Methods, 17(3):261–272, March 2020. doi: 10.1038/s41592-019-0686-2.

125. TensorFlow Developers. Tensorflow. Zenodo, 2017. doi: 10.5281/zenodo.12726004.

126. F. Pedregosa, G. Varoquaux, A. Gramfort, V. Michel, B. Thirion, O. Grisel, M. Blondel, P. Prettenhofer, R. Weiss, V. Dubourg, J. Vanderplas, Passos, D. Cournapeau, M. Brucher, M. Perrot, and E. Duchesnay. Scikit-learn: Machine learning in Python. Journal of Machine Learning Research, 12:2825–2830, 2011.

